# Probing the scalability of ultra stable catch bond complexes

**DOI:** 10.64898/2026.01.08.695900

**Authors:** Zarah Walsh–Korb, Sean Boult, Rosario Vanella, Intizar Ali Tunio, Jiajun Li, Vanni Doffini, Inam Ul Ahad, Michael A. Nash

## Abstract

The catch bond complex between serine-aspartate repeat protein G (SdrG) from *Staphylococcus epidermidis* and the beta chain of fibrinogen (Fgβ) exhibits two distinct rupture populations when dissociated under tensile force. Such complexes present exciting possibilities for developing dynamic biomaterials due to their unique response to shear force. However, the environmental responsiveness of this complex and its influence on adhesion behaviour in multi-valent systems remain underexplored. Using AFM-single molecule force spectroscopy (AFM-SMFS) and spinning disk adhesion (SDA) assays, we examined how protein orientation, mutations, and environment influence the stability and scaling behaviour of this catch bond system. Our findings confirmed that anchor point location (i.e., the direction from which the protein is pulled) strongly influences catch bond behaviour, while an S338H mutation in the binding domain destabilised the interaction in both single-molecule and multi-valent adhesion assays. This research examines how catch bond behaviour translates from the nanoscale to microscale using single molecule and multi-valent cell adhesion measurements and provides a toolkit for exploiting catch bonds towards macroscale material applications.

## Main

In recent years, interest in shear-responsive materials has increased significantly. This interest is primarily directed at shear-thinning materials for applications in 3D-printing and drug delivery,^1–3^ ensuring that materials can be printed or injected in a targeted manner and re-stabilise after introduction. However, there are also a number of applications in which shear-strengthening materials would be advantageous, e.g., materials designed to stem uncontrolled blood loss (haemorrhaging) must withstand high shear forces that would damage shear-thinning materials.^4^ These materials could be enhanced by the use of multi-regime or dilatant fluids,^5^ allowing placement of materials and high-performance under stress (shear-strengthening), a feature ultra stable catch bonds are particularly suited to provide.^6^

Catch bonds defy the common perception of how intermolecular interactions behave under shear stress, becoming more stable and resisting rupture under the influence of force. In contrast, classical ‘slip’ bonds are destabilised and break more easily under the influence of shear force. Although discussed theoretically since the 1980’s,^7^ experimental evidence for catch bonds first emerged 20 years ago, which coincided with methodological improvements in single-molecule analysis techniques.^8^ Catch bond complexes modulate a number of physiological processes, e.g., leukocyte rolling,^9^ desmosomal adhesion,^10^ sperm-egg adhesion,^11^ and cellular transport through dynein-microtubule interactions.^12^ They are also suspected to stabilise the power stroke in myosin II-actin interactions.^13,14^ Catch bond forming proteins have also been identified among the cell surface proteins of pathogenic bacteria, playing a role in the infection and colonisation of host organisms. For example, SdrG from *Staphylococcus epidermidis*,^15,16^, SdrD *from Staphylococcus aureus,* SpsD and SpsL from *Staphylococcus pseudintermedius*,^17^ BBK32 from *Borrelia burgdorferi,*^18,19^ *and* FimH from *Escherichia coli,*^20^ form catch bonds with fibrinogen, fibronectin (SpsD, SpsL, BBK32), and mannosylated proteins, respectively.

Many investigations have focused on identifying catch bonds,^15,16,21^ quantifying their maximum rupture forces,^16,22^ as well as exploring their role in disease and/or physiological processes.^13,23,24^ The extraordinary response of catch bonds to shear stress makes them important candidates in the development of shear-responsive materials. Several attempts have been made to design and optimise biological catch bonds and catch bond cross-linked materials *in silico* and *in vitro*,^25–32^ but applications are limited by an incomplete understanding of their stability and scalability. Research on the SdrG:Fgβ catch bond has explored thermal sensitivity and the Ca^2+^ mediated stability of the adjacent B-domains.^15,33,34^ However, the extent of tunability based on mechanical, environmental and molecular geometric perturbations, and how this translates across length scales is unclear.

Recent work from this group has compared AFM-SMFS with SDA assays to correlate molecular interactions with cell adhesion strength.^35^ SMFS provides detailed insight into the behaviour of molecular interactions and how they change as a function of tensile stress, but it neglects multi-valency, which is often relevant for adhesive proteins like SdrG. In SDA assays shear flow is applied and the adhesion profile of a cell population is measured, providing insight into cooperative multi-valent interactions that govern the adhesion strength at the cellular scale. We further exploited these techniques, alongside numerical modelling to study the mechanical stability of the SdrG:Fgβ complex (Figure **1**) as a function of chemical environment, shear stress, pulling geometry, binding pocket mutations, and protein/peptide ligands (i.e., the full fibrinogen (Fg) protein *vs.* Fgβ peptide).

**Figure 1:**
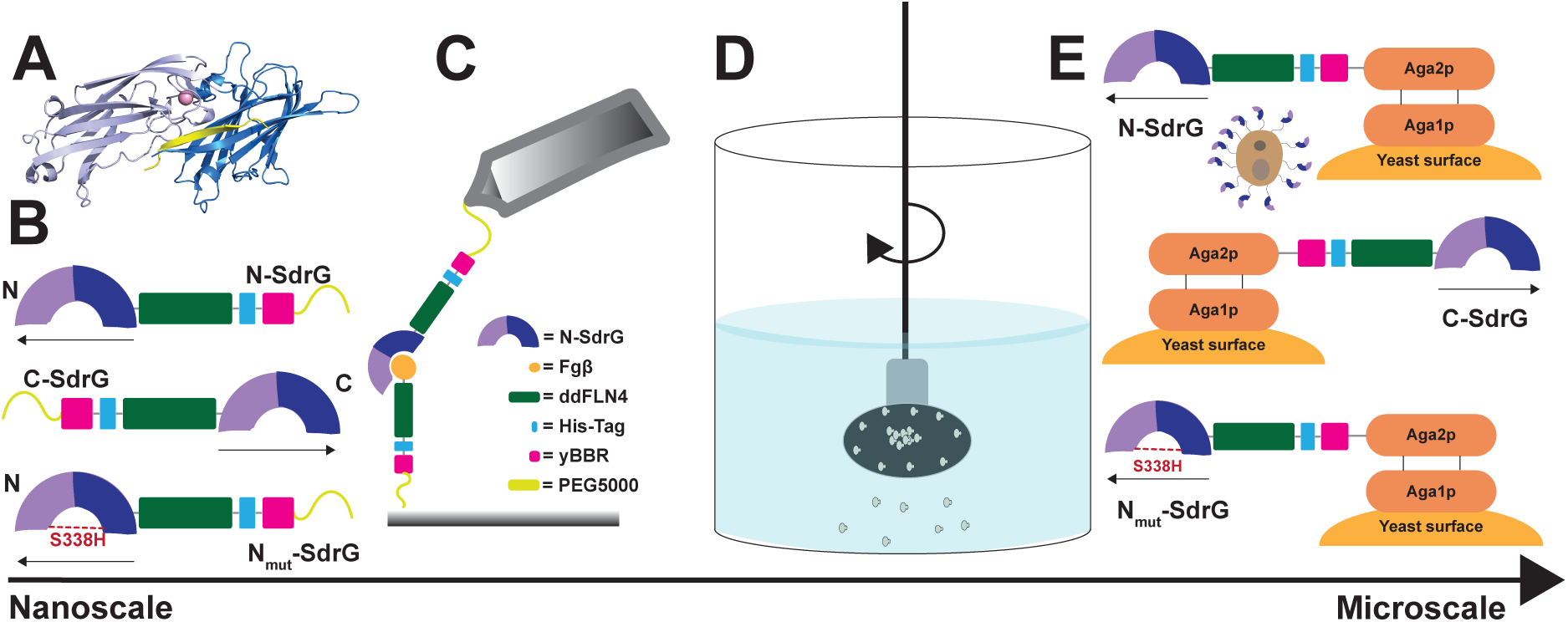
Examining mechano-response of SdrG:FgB interactions from the nano- to the micro-scale. ***Nanoscale*** (**A**) Crystal structure of the SdrG:Fgβ complex (PDB 1R17), (**B**) Graphical representation of the three protein constructs under investigation: the native pulling geometry (N-SdrG), the non-native pulling geometry (C-SdrG), and the native pulling geometry with a binding site mutation S338H (Nmut-SdrG). (**C**) Schematic of the AFM-SMFS setup. A cantilever is functionalised with the SdrG variant, and the substrate is modified with the Fgβ peptide or full-length fibrinogen. Constant speed AFM is then used to measure the rupture forces. ***Microscale***. (**D**) Graphical illustration of the spinning disk adhesion assay in which an inverted functionalised coverglass with adherent yeasts is spun. The spatial distribution of cells after spinning is used to quantify adhesion strength. (**E**) Graphical illustration of the three yeast surface display constructs under investigation: N-SdrG (insert representation of the decorated yeast surface), C-SdrG, and Nmut-SdrG, in which the native pulling geometries of the N-SdrG and Nmut-SdrG are achieved by expressing the protein constructs at the N-terminus of the Aga2p domain using a pCHA expression vector, and the non-native configuration of the C-SdrG is achieved by expressing the construct at the C-terminus of the Aga2p domain using the pYD1 vector.

By examining both single-molecule interactions and multi-valent adhesion behaviour after displaying the protein constructs on the surface of *Saccharomyces cerevisiae*, we determined which factors more strongly influence behaviour at each length scale and provide insight into how the mechanostability of the SdrG:Fgβ complex manifests at different length scales. This combination of techniques allows us to characterise this complex in detail and facilitates the future design of tailored shear-modulating materials mediated by catch bond formation.

## Results

We studied in total six pairs of protein complexes comprising one of three SdrG constructs (N-SdrG, C-SdrG and N_mut_-SdrG) bound to either the Fgβ peptide or full-length fibrinogen (Fg). N-SdrG (Figure **1A** and **1B**) represents the native pulling configuration in which the C-terminus is tethered to the AFM cantilever or Aga2p domain on the yeast surface and the N-terminus remains free. C-SdrG represents the non-native pulling configuration where the N-terminus was anchored to the cantilever or yeast surface. In the case of N_mut_-SdrG, the protein was tethered in the native configuration at the C-terminus and contained a mutation (S338H)^36^ that inhibits binding to Fgβ/Fg (Figure **1B**). The SdrG constructs for the SDA assay (Figure **1D**) displayed on the surface of *S. cerevisiae* (Figure **1E**) are also denoted N-SdrG, C-SdrG and N_mut_-SdrG with the same naming conventions and were anchored via the Aga2p surface display anchor domain (Figure **1F**). Our experimental approach aimed to study how changes in the chemical environment (i.e., Ca^2+^ concentration) and the choice of target ligand (Fgβ vs. Fg) influenced the force/shear stress response in these protein constructs. We were also interested in comparing differences in response between single protein complexes (in a defined nanoscale environment) measured by SMFS and multiple protein complexes tested in parallel (i.e., a defined microscale environment) using the yeast adhesion assay.

### Ca^2+^-dependence of SdrG:Fgβ interactions

Previous studies highlighted the role of Ca^2+^ in enhancing and stabilising the mechanical properties of the SdrG:Fgβ complex, as well as the adjacent B domains.^15^ The N2/N3 domains of SdrG contain a Ca^2+^ binding site, and its role in the catch bonding mechanism is not completely clear. Recent studies on the related protein SdrD identified that Ca^2+^ is critical for the mechanical stability of the binding pocket and the B-domains, and here we identify a similar behaviour in SdrG.^37^ To measure the calcium sensitivity of the SdrG complexes, we examined the behaviour of the N-SdrG:Fgβ complex using SMFS in TBS buffer containing 1 mM, 2.5 mM, 6.25 or 10 mM Ca^2+^ at 22°C. Using a dynamic force spectroscopy methodology similar to our previously reported studies,^11,38,39^ N-SdrG was brought into contact with an Fg- or Fgβ-functionalised surface, and retracted at pulling speeds of 400, 800, 1600 and 3200 nm/s (Figure **2**). As pathogenic bacteria move along physiological surfaces, having the cantilever was functionalised with SdrG to more closely reconstruct this scenario. As *S. epidermidis* is found on the skin and in mucus membranes,^40^ we hypothesised that optimal behaviour of SdrG may be related to physiological conditions.^37^ While physiological Ca^2+^ concentrations in blood and mucus are ∼1.8 mM, the concentration of Ca^2+^ in the skin consists of a gradient ranging from 0 mM to 4 mM depending on the skin layer, where *S. epidermidis* is also commonly found.^40,41^

**Figure 2:**
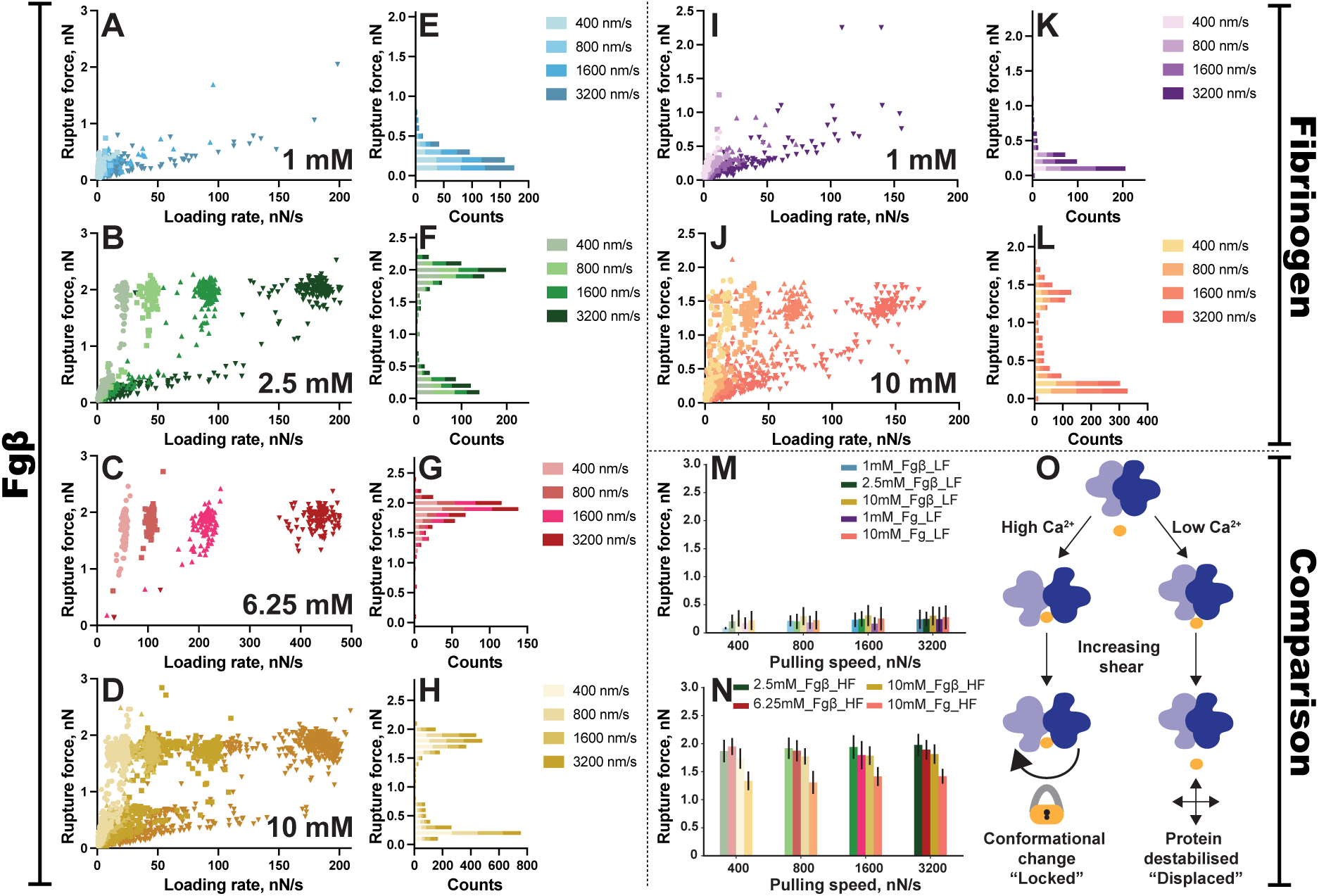
Investigation of single molecule mechanostability as a function of Ca^2+^ concentration and target ligand. Plots of rupture force (nN) vs. loading rate (nN/s) in constant speed AFM-SMFS measurements on the N-SdrG:Fgβ complex (N-SdrG on cantilever, Fgβ on substrate) in TBS buffer with (**A**) 1 mM, (**B**) 2.5 mM, (**C**) 6.25 mM and (**D**) 10 mM CaCl2. Histograms of the rupture force distributions from the same experiments are shown in panels **E-H**. Plots of rupture force (nN) vs. loading rate (nN/s) in constant speed AFM-SMFS measurements on the N-SdrG:Fg complex (N-SdrG on cantilever, Fg on surface) in TBS buffer with (**I**) 1 mM and (**J**) 10 mM CaCl2. Histograms of the rupture force distributions from the same experiments are shown in panels **K-L**. Comparison of median rupture forces across complex type and TBS buffer Ca^2+^ concentration for low force (**M**) and high force (**N**) ruptures. (**O**) Graphical illustration of the mechanism by which a low or high force rupture occurs. For all conditions, n = 3 independent datasets.

The most salient result is the bimodal distribution of rupture forces that is characteristic of the catch bond for this complex, particularly at intermediate calcium levels (Figures **2B, 2F**).^42^ The median rupture forces increase slightly with increasing pulling speed, and at one point there is a clear shift in rupture event population from a low force to a high force regime. Most importantly, we observe a calcium dependency as the Ca^2+^ concentration increases from 1 mM to 10 mM. At 1 mM there are few observable high force rupture events with median rupture forces in the 200-500 pN range (Figures **2A, 2E**). At 2.5 mM we see bimodal behaviour characteristic of the catch bond, and at 6.25 mM we no longer observe the low force rupture events (Figures **2C, 2G**). Surprisingly, at 10 mM Ca^2+^ we found that the low force population returned (Figures **2D, 2H**). The reasons for this remain unclear. More extensive data sets are shown in the ESI (Section S2). From these data, we conclude that 1 mM Ca^2+^ is likely too low to completely saturate the Ca^2+^ binding site within SdrG. This could result in the complex being destabilised and unable to activate to the high stability state, resulting in only low force rupture events at 1mM Ca^2+^ (Figure **2O**).

When changing the target ligand from the Fgβ peptide to full-length fibrinogen (Fg), we see the same trend (Figures **2I-L** and Figure **2O**). Mechanical shear has some effect on the ability of the complex to form a catch bond but Ca^2+^ concentration in the measurement buffer can hinder or enhance the formation of the complex to a much larger degree. Once again at 1 mM, we primarily see low force interactions with a median rupture force of 200 pN across all pulling speeds. At 10 mM a more sizable high force population is visible with a median rupture force of approximately 1.4 nN (Figures **2M, 2N**). Changing the substrate to the full Fg chain also allowed us to visualise the effect of non-specific interactions, indicated by the overall lowering of the maximum rupture forces. This was particularly noticeable in the high force populations. Non-specific interactions between the Fg chain and the cantilever result in a reduction of the median high force rupture by almost 25%.

As a negative control, a variant of the N-SdrG was prepared that contained a point mutation at the S338H position, denoted N_mut_-SdrG. In a TBS buffer solution containing 10 mM Ca^2+^ against both Fg and Fgβ, few specific interactions (8-35 in 2000 measurements) were observed over a 12 h measurement period, indicating this variant is incapable of binding the target. This negative control data is provided in the ESI (Section S2, Figures **S2.8** and **S2.9**).

### Pulling direction influences force activation

An important consideration for exploiting catch bonds in macroscopic materials is the direction that force is applied to the complex (i.e., the pulling point). This property can be defined through site-specific surface immobilisation or, in the case of the SDA assays, by the topology of the surface display protein anchor. As such, the C-SdrG:Fgβ interaction was investigated in the same way as the N-SdrG:Fgβ and N-SdrG:Fg interactions previously, with the exception that only 1 mM Ca^2+^ and 10 mM Ca^2+^ were explored (Figures **S2.10-S2.13**).

In contrast to N-SdrG:Fgβ, no significant difference was observed between force spectra for C-SdrG:Fgβ obtained at 1 mM or 10 mM Ca^2+^ (Figures **S2.10** and **S2.11**). Notably, there is a distinct absence of high force populations, regardless of pulling speed. The histograms of the rupture force distribution (Figures **S2.10** and **S2.11**) confirm that the distribution is monomodal and rupture forces are concentrated in the low force pathway. High force ruptures were not fully eliminated from any population. However, the likelihood of catch bond formation was significantly reduced, in fact almost negligible, and rupture forces were lower than those observed for N-SdrG:Fgβ.

Repeating these experiments with Fg, in place of Fgβ, a similar trend is observed. The plots of loading rate vs rupture force show a slight increase in the number of high force ruptures with 1 mM and 10 mM Ca^2+^ (Figures **S2.12A** and **S2.13A**, respectively), as do the related histograms (Figures **S2.12B** and **S2.13B**). Comparing the histograms from the Fg series with those obtained in the Fgβ series (Figures **S2.12B** and **S2.13B** vs Figures **S2.10B** and **S2.11B**) it seems maximum average rupture forces are more dependent on force than in the Fgβ series (Figure **S2.10B** and **S2.11B**), although the increase is small. This suggests that the non-specific interactions introduced by other amino acids in the Fg chain may have a greater effect on the overall stability of the complex in this orientation than for the native pulling geometry. Despite this minor change, the strength of the interaction is still significantly reduced and the high force pathway almost inaccessible.

### Deciphering binding modes with Monte Carlo simulations

As we see in Figure **2**, there is a clear difference in the observed interactions of the N-SdrG:Fgβ and N-SdrG:Fg complexes as a function of environmental Ca^2+^ concentration. To better understand how environment impacts the interactions between N-SdrG and Fg/Fgβ, we extrapolated the kinetic data from the plots of rupture force *vs.* loading rate using the Bell-Evans equation (ESI, Section S3).^11,43^

Previous investigation of binding pathways in cohesin-dockerin and Izumo1-Juno catch bonds published by this group verified a multi-pathway kinetic model for these catch bonds using Monte Carlo (MC) simulations.^11,39^ MC simulations are useful for statistically exploring possible kinetic models and hidden energy barriers in complex biological interactions. Given the bimodal distribution of forces observed here, and literature proposing that N-SdrG:Fgβ undergoes a conformational change to activate the characteristic high force rupture, a two-pathway kinetic model was once again assumed. We probed the data using three different models to decipher how the high force pathway is activated and how sensitive it is to its environment. The reversible transition two-pathway (rttp) model assumes there are two pathways, and both pathways are accessible to each interaction, while the irreversible transition two-pathway (ittp) model, assumes that interactions that follow the high force rupture pathway cannot transition back to the low force rupture pathway. The third model employed was a multiple binding modes two-pathway (mbmtp) model, in which it is assumed that there are at least 2 pathways and transitions can be made between them, Figures **S3.1-S3.16**.^11,34,39,44–46^ However, while the rttp model fit the 1 mM Ca^2+^ dataset well, it failed for higher Ca^2+^ concentrations, which were better fit with the mbmtp model. The mbmtp also correlates well with the established Dock-Lock-Latch mechanism and MD simulations of similar interactions. However, we observed that no single model could correctly predict the behaviour observed in the experimental data based on the kinetic data alone, (ESI S3). It is clear from the experimental data that Ca^2+^ influences catch bond behaviour, and that mechanical shear does not induce a catch bond if Ca^2+^ concentration is too low. This is highlighted in Figure **2M** and **2N**, which show the ratios of high force:low force ruptures, as a function of Ca^2+^ concentration. Modifying the mbmtp model to include a parameter linking experimental outcome to mechanical shear and Ca^2+^ concentration (Ca-mbmtp), resulted in more accurate predictions of the conformational ensemble (Figures **3A-D**).

**Figure 3:**
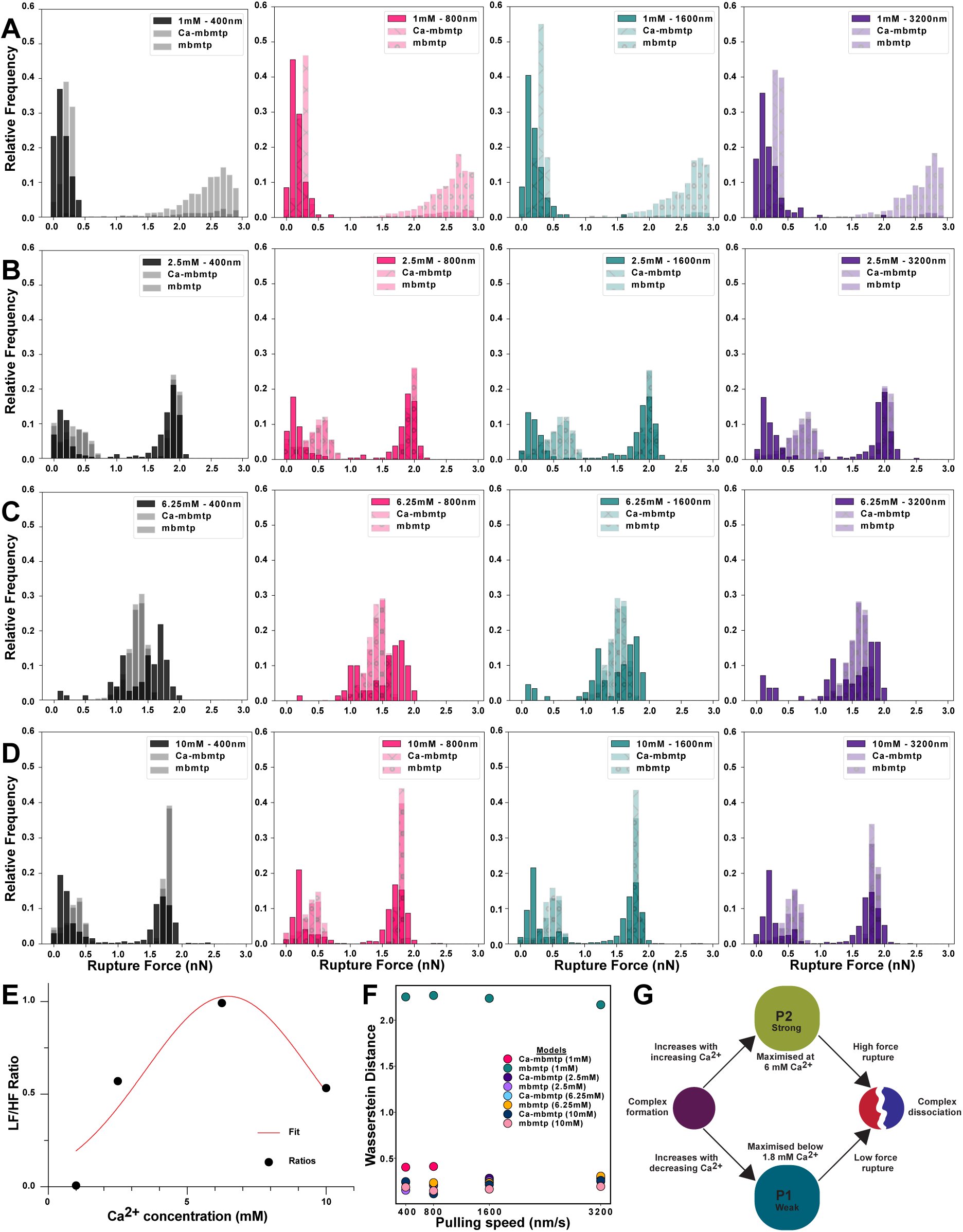
Deciphering the role of the chemical environment in the binding pathways of the N-SdrG:Fgβ catch bond. Histograms of rupture force (nN) vs frequency count of the experimental data obtained from the single molecule force spectroscopy measurements on the N-SdrG:Fgβ complex overlaid with Monte Carlo simulations of both the multiple binding mode two-pathway (mbmtp) and Ca-dependent multiple binding mode two-pathway (Ca-mbmtp) models at when Ca^2+^ concentration is set at (**A**) 1 mM, (**B**) 2.5 mM, (**C**) 6.25 mM and (**D**) 10 mM. Through each line, black refers to data collected at 400 nm/s, pink at 800 nm/s, teal at 1600nm/s and purple at 3200 nm/s. (**E**) Plot of Ca^2+^ concentration (mM) vs low force/high force rupture ratio used to adjust the mbmtp model to account for Ca^2+^ concentration. (**F**) Scatter plot of the Wasserstein distance as a function of pulling speed for the mbmtp and Ca-mbmtp models. (**G**) Graphical illustration of the two binding pathway model for a Ca-dependent binding model for this complex.

The so-called Ca-mbmtp model relies on the distribution of pathways in the system being an intrinsic or deterministic property that is primarily affected by the chemical environment, rather than applied shear forces. Using the Wasserstein distance, a non-parametric test to determine similarity between two probability distributions, we estimated the validity of this approach (Figure **3C**).^47,48^ We see that the mbmtp and Ca-mbmtp models showing a low Wasserstein compared to all other models except at the 1 mM condition at all loading rates. This highlights to define the binding behaviour N-SdrG:Fgβ, the calcium concentration must be considered. This suggests that, contrary to previous hypotheses, the catch bond behaviour of the N-SdrG:Fgβ complex is more dependent on chemical environment than applied force, at least at the nanoscale, illustrated in Figure **3G**.

### Understanding resistance of multivalent interactions to shear forces

Nanoscale investigation of SdrG:Fg and SdrG:Fgβ complexes identified chemical, mechanical and geometrical parameters essential to stable activation of individual high force catch bonds. To create and control macroscale catch bond-mediated materials, we must also understand how SdrG:Fg or SdrG:Fgβ interactions cooperate (or perhaps compete) with one another at the microscale. Recent work from this group compared nanoscale measurements of monomeric streptavidin (mSA)-biotin complexes to their cellular scale adhesion using SDA assays.^35^ This permitted correlations between defined nanoscale interactions and multiple interactions (up to 50,000 potential simultaneous interactions)^49^ in a defined microscale space, although without the formation of a catch bond.

By surface displaying N-SdrG, C-SdrG and N_mut_-SdrG on *S. cerevisiae*, multivalent interactions of N-SdrG:Fg/Fgβ, C-SdrG:Fg/Fgβ and N_mut_-SdrG:Fg/Fgβ were examined, under the same environmental conditions as SMFS, using the SDA assay. Spinning was performed at 3000 and 4000 rpm in TBS buffer containing 1 mM Ca^2+^ or 10 mM Ca^2+^, to investigate the influence of both shear stress and environment on cooperative catch bond interactions at the microscale. Each Fg/Fgβ-coated glass disk is seeded with 3 million yeast cells displaying the construct of interest prior to spinning and cells remaining after spinning are analysed using an in-house developed python workflow to extract both positional and force-related information, see ESI **S5** and **S6**. Heatmaps of the distribution of cells on both Fg and Fgβ-functionalised substrates incubated with N-SdrG (Figures **4A**,**4C** with Fg, Figures **4I,4K** with Fgβ), and C-SdrG displayed cells (Figures **4B**,**4D** with Fg, Figures **4J,4L** with Fgβ) were analysed. The cell populations are not normalised to visualise how cell distribution is dependent on shear force and catch bond orientation.

**Figure 4:**
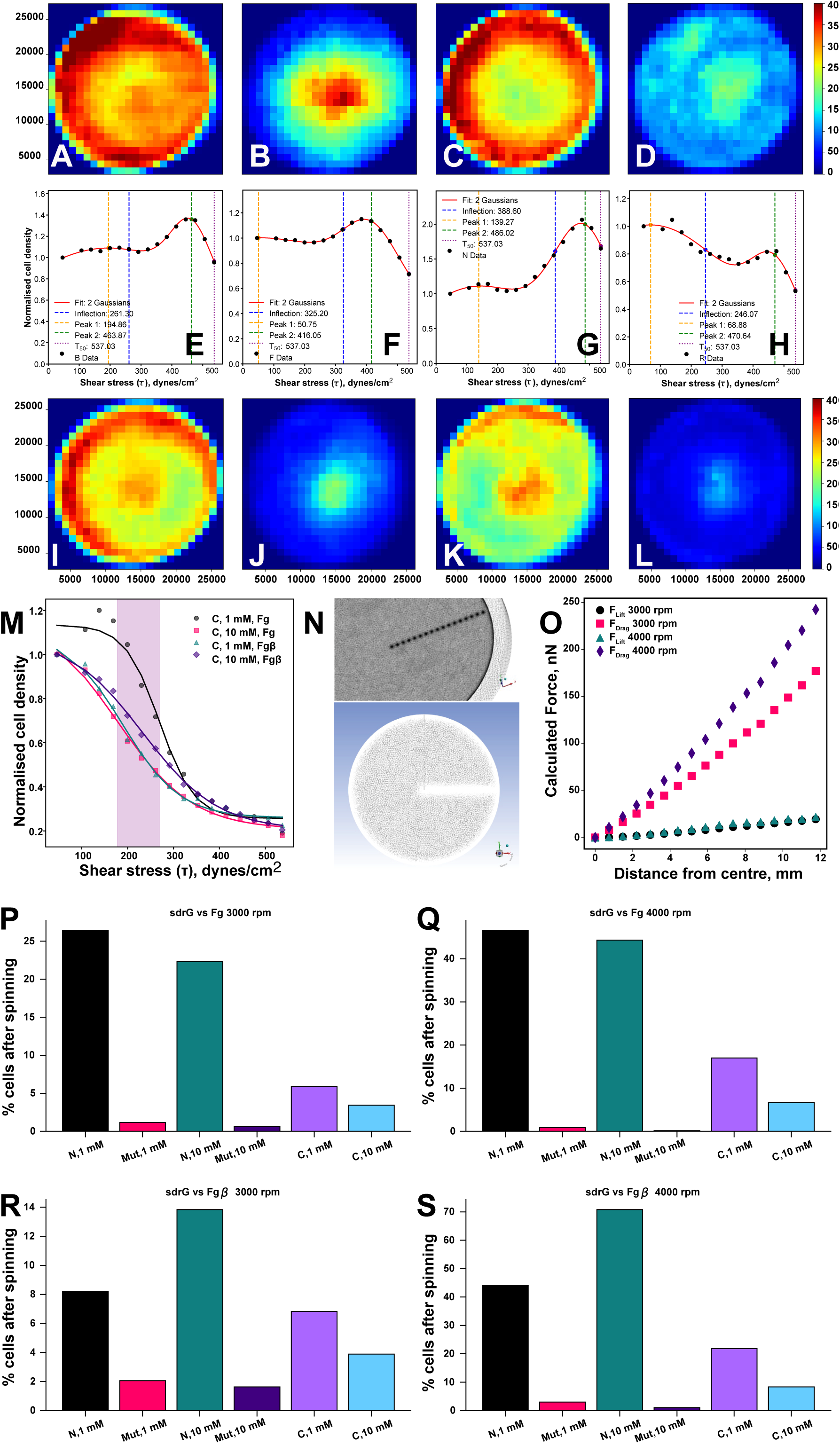
Analysis of SdrG-mediated yeast cell adhesion under shear stress. Cumulative density (CD) plots of cell distribution on 25 mm diameter Fg-coated glass disks after spinning at 4000 rpm with (**A**) N-SdrG at 1 mM Ca^2+^, (**B**) C-SdrG at 1 mM Ca^2+^, (**C**) N-SdrG at 10 mM Ca^2+^, (**D**) C-SdrG at 10 mM Ca^2+^. Fitting of catch bonding datasets using a double gaussian fitting model to extract low and high force binding and the onset of the high force catch bond for N-SdrG at 4000 rpm on (**E**) Fg at 1 mM Ca^2+^, (**F**) Fg at 10 mM Ca^2+^, (**G**) Fgβ at 1 mM Ca^2+^, (**H**) Fgβ at 10 mM Ca^2+^. CD plots of cell distribution on 25 mm diameter Fgβ-coated glass disks after spinning at 4000 rpm with (**I**) N-SdrG at 1 mM Ca^2+^, (**J**) C-SdrG at 1 mM Ca^2+^, (**K**) N-SdrG at 10 mM Ca^2+^, (**L**) C-SdrG at 10 mM Ca^2+^. (**M**) Fitting of non-catch bonding datasets (C-SdrG) using a sigmoidal fit to extract τ50 (indicated by the purple band) at 4000 rpm on both Fg and Fgβ substrates. (**N**) Computational fluid dynamics (CFD) model of the glass disk surface with one row of cells originating at the centre of the disk, with 17 points in total (top) zoom, (side) overview. (**O**) Radial cell position on the disk surface in CFD in mm vs calculated hydrodynamic force (nN) to detach cell in both the y (drag) and z (lift) directions. (**P-S**) Bar charts of % cells remaining after spinning for N-SdrG, C-SdrG and Nmut-SdrG at both 1 mM and 10 mM Ca^2+^ at (**P**) 3000 rpm on Fg, (**Q**) 4000 rpm on Fg, (**R**) 3000 rpm on Fgβ, and (**S**) 4000 rpm on Fgβ.

Depending on spinning speed, substrate and Ca^2+^ concentration, 8-70% of N-SdrG displayed cells remain on the surface post-spinning, dropping to 4-20% for C-SdrG displayed cells. Only 2% of cells displayed with the non-binding mutant, N_mut_-SdrG, remain adhered across all conditions showing no dependence on shear force or Ca^2+^, Figures **4P-S** and Figures **S6.1**, **S6.2**, **S6.3**. Interestingly, C-SdrG adheres better at 1 mM Ca^2+^ than 10 mM Ca^2+^, irrespective of substrate coating. However, adhesion increases as a function of shear force, from ∼8% at 3000 rpm to ∼20% at 4000 rpm for 1 mM Ca^2+^, and from ∼4% to ∼10% at 10 mM, a 2.5-fold increase across both conditions. Despite the non-optimal orientation, it appears to be possible to activate some catch bonds through shear, as seen in the SMFS previously.^50^

As with the SMFS data, the N-SdrG datasets show more complex behaviour, in which multiple dependencies are evident. On Fg-coated substrates, we see that at 4000 rpm (Figure **4Q**) more cells remain adhered post-spinning than at 3000 rpm (Figure **4P**), correlating with force activation of the catch bond. The difference in adhesion between the 1 mM and 10 mM Ca^2+^ concentrations is also minimal, in contrast to the data obtained by SMFS. On the Fgβ-coated substrates, however, there is a substantial difference between cell adhesion as a function of speed (an increase from 8% to 45% at 1 mM and 14% to 72% at 10 mM Ca^2+^), and between the 1 mM and 10 mM Ca^2+^ datasets (8 vs 14% at 3000 rpm and 45 vs 72% at 4000 rpm) (Figures **4R** and **4S**). We hypothesise that the non-specific interactions and the presence of two Fgβ domains in close proximity^51^ on the full Fg chain help maintain the cells at the disk surface long enough so that the catch bond can be activated. However, this non-specific adhesion interferes with the Ca-dependent behaviour of the displayed N-SdrG construct. When Fgβ is used, the applied shear at 3000-rpm does not appear to properly activate the catch bond across the entire contact area at 1 mM Ca^2+^, but the 10 mM Ca^2+^ does increase adhesion by 57%, correlating strongly with the SMFS data. At 4000 rpm, the increased applied shear appears sufficient to activate the catch bond even at low Ca^2+^ but increasing the Ca^2+^ concentration does result in a 62% increase in cell adhesion post-spinning. This highlights both the shear and Ca-dependent nature of the catch bonds at the microscale, in contrast to their strongly Ca-dependent nature at the nanoscale.

### Cooperativity contributes to cell adhesion

More than differences in cell adhesion, the distribution of cells across the disk surface as a function of protein orientation, protein/peptide or mutant merits closer examination. For all C-SdrG datasets, and N-SdrG at 3000 rpm with 1 mM Ca^2+^ and the Fgβ substrate, we observe classic slip bond behaviour. In SDA assays, this is observed as higher cellular adhesion at the centre of the disk, where shear forces are lower, in comparison with the edges where the shear forces are higher, Figure **S6.1**. A common analysis parameter in SDA assays is the tau_50_ (τ_50_) value, i.e., the shear stress that detaches 50% of the cells. This can provide information on the cooperative adhesion of the surface expressed proteins. For the slip bonding datasets, a sigmoidal fit or a simple x at half y-max extraction (two datasets have a more convex shape that gave a low fitting accuracy, see ESI Section **S6**) were used to extract this parameter, Table 1. Due to low adhesion post-spinning of N_mut_-SdrG-displayed cells (less than 2% of incubated cells), no meaningful fitting of the datasets could be done, this highlights that the binding site mutation is effective at hindering binding even in a cooperative scenario.

**Table 1:**
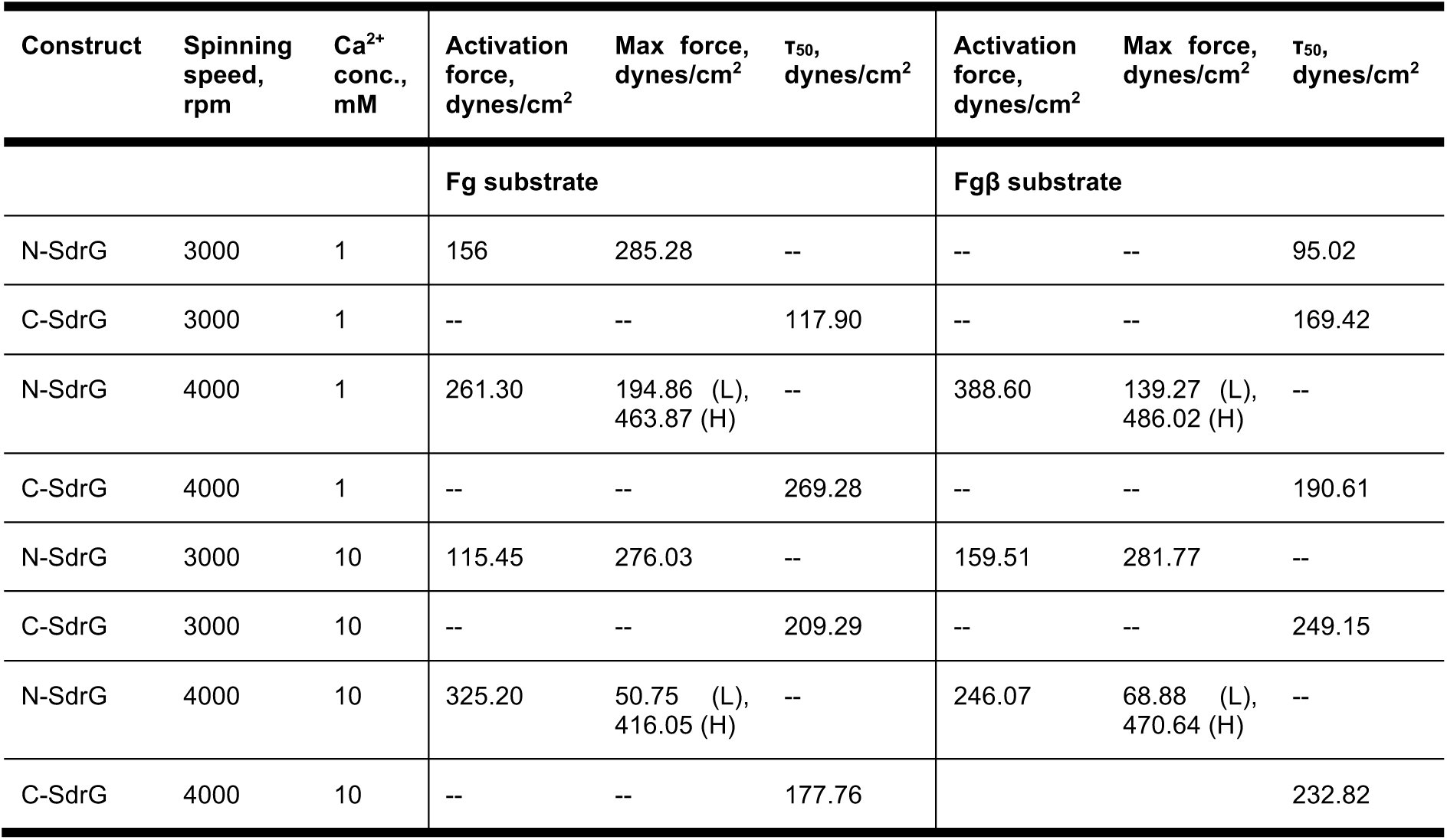
Activation force, maximum force and τ_50_, all in dynes/cm^2^ obtained from spinning disk microscopy for both N-SdrG, C-SdrG, and N_mut_-SdrG constructs against Fgβ and Fg substrates.

While τ_50_ provides meaningful insight on slip bonding interactions, it fails to fully explain the complexity of catch bonding interactions at the microscale. Due to the nature of the SDA assay, apparent forces increase from the centre to the edges of the disk. As catch bonds do not bind strongly at low shear, there may be regions of high cell density at no shear (origin), low cell adhesion (low shear region) and then increasing cell adhesion towards the edges where shear is highest, as seen in Figures **4A, 4C, 4I, 4K** above.

Moreover, the spinning speed had a significant impact on the pattern observed. Data collected at 3000 rpm (except for dataset N-SdrG:Fgβ with 1 mM Ca^2+^, which exhibited slip bond behaviour) could be fitted with a mixed sigmoid/gauss function, data collected at 4000 rpm could only be fit with a double gaussian. This is interesting because it gives insight into intrinsic behaviours of the catch bond cooperativity. The combination of sigmoid and gaussian distributions suggests that slip and catch bonds are present simultaneously, as observed by SMFS. Viewed from a different perspective, it suggests the applied shear forces are adequate to activate some of the population but not all. The double gaussian however, suggests that there are two force dependent populations. This could be two conformations of the complex, as has been suggested for some catch bond complexes at the nanoscale, or, at the microscale this may indicate that the catch bond and cooperative adhesion create two distinct behaviours. Comparing the single gaussian in the 3000 rpm datasets to the double gaussian in the 4000 rpm datasets, we see that the single gaussian exists between 276-285 dynes/cm^2^ with an onset visible between 115-159 dynes/cm^2^. The double gaussian datasets show an initial peak at 50-68 dynes/cm^2^ (Fgβ) or 139-194 dynes/cm^2^ (Fg), suggesting initial cooperative adhesion stabilised by force, which appears to be enhanced by non-specific interactions of the Fg substrate. These stabilised populations are then fixed in place to undergo a conformational change to access the high force adhesion regimes, which occurs between 246-388 dynes/cm^2^, resulting in a second gaussian at 416-486 dynes/cm^2^.

### Estimating loading rate at the microscale

To decipher the correlation between the nanoscale and microscale behaviours of the catch bond complexes, we used computational fluid dynamics (CFD) with ANASYS to simulate the disk surface (Figure **4N**). This allowed us to determine wall shear stress across the disk when no particles (cells) are present (see Figure **S4.1**) and drag and lift forces acting on the surface at 3000 and 4000 rpm (Figure **4O**) with cells present. We simulated a row of cells (17 in total) at regular intervals along the radius of the disk starting at the origin (Figure **4N**) to determine applied shear at each position, which is a combination of wall shear stresses across the surface, around the cells and their nearest neighbours (see Figure **S4.1**). With this information we could calculate the maximum loading rate in nN/s (as a function of the spinning speed) per position to facilitate comparison with SMFS data. At 3000 rpm the max loading rate at the edge of the disk is 177.01 nN/s, while at 4000 rpm it is calculated to be 242.3 nN/s. The values obtained from the CFD model are similar to those from the 3200 nm/s pulling speed datasets of the SMFS (approximately 200 nN/s), indicating that the SDA assay is only comparable with the higher pulling speed datasets. Given the high level of surface display of the constructs on the yeast cells (Figures **S5.3, S5.4**) achieved with this system, spinning at less than 3000 rpm failed to remove enough cells for adequate analysis, thus it was not possible to compare the SDA assay results to the lower pulling speed datasets from SMFS.

The CD plots for the 10 mM Ca^2+^ concentration at 4000 rpm in Figures **4A-D** and **4I-L** (see ESI **S6** for other conditions), show that cells are concentrated at 0-4 mm and 10-12 mm from the centre, the low shear and activated catch bonds, respectively. Taking the median SMFS adhesion noted above of 1.9 nN and the calculated forces per distance in Figure **4O**, indicates that in the low shear region 28-39 catch bonds adhere per yeast cell, while in the high shear region ∼93 (>3-fold) catch bonds are activated at 3000 rpm, and ∼127 (>3-fold) at 4000 rpm. The patterns of adhesion look extremely similar between 3000 and 4000 rpm for N-SdrG, yet the numbers of cells remaining after spinning are very dependent on the spinning speed, Ca^2+^ concentration and the nature of the substrate. This suggests it may not be the actual number of activated catch bonds per cell, but the kinetics and cooperativity of their activation that plays the strongest role. This is the subject of further computational investigation, as experimental determination of shear-dependence in binding kinetics is limited by the inability to introduce shear forces into standard methods, such as isothermal titration calorimetry.

## Discussion

Exploring SdrG:Fg(β) interactions under a variety of environmental conditions and orientations over two length-scales reveals several behaviours related to the scalability of catch bond complexes. At the single molecule scale we see a rather deterministic relationship with tensile force in all datasets, suggesting that complex formation resulting in a high force rupture or ‘catch’ is an intrinsic property of the SdrG:Fgβ interaction with a higher sensitivity to environmental calcium than applied force. Monte Carlo simulations also indicated that the best fitting model (Ca-mbmtp) was not only force dependent but included a Ca-dependent parameter. Our observations show that catch bond behaviour of SdrG:Fg is highly dependent on Ca^2+^ concentration and orientation of the SdrG with respect to the ligand (Fg/Fgβ). The ligand itself also plays a role, but to a much lesser extent. At low Ca^2+^ concentrations in the non-native pulling geometry (C-SdrG), catch bond behaviour is almost eliminated. At high Ca^2+^ catch bonds re-appear but never reach the same median rupture force of the native orientation, indicating that the conformational change necessary to produce the high force rupture is not accessible from all pulling orientations. Mutation, by replacing serine at position 338 of the SdrG with histidine to block entry to the binding domain, prevents catch bond formation, or binding in general, at any Ca^2+^ concentration, as expected.

However, some nanoscale observations disappear as we transition to the microscale, while other trends emerge. At the nanoscale force activation is highly Ca-dependent, with almost no high force ruptures observed in any orientation at low Ca^2+^ concentration. At the microscale, likely due to cooperation between adjacent complexes, this dependence changes. Catch bond behaviour can be observed in both N-SdrG:Fg and N-SdrG:Fgβ at low Ca^2+^. However, the calcium stabilisation of the individual cells seems to impact both the maximum force achievable by the system and the number of cells that remain adhered after spinning. Furthermore, under high Ca^2+^ and high shear stress, even the C-SdrG:Fgβ complex can be induced to adhere to both substrates, which was not seen at the molecular scale and is likely only possible due to cooperative interactions between adjacent proteins. The influence of ligand type, Ca^2+^ concentration and shear force are more complex at the microscale, with highly specific ligand interactions (with Fgβ), high Ca^2+^ and high shear required to ensure adequate activation of the catch bond. CFD investigations of the fluid parameters of the SDA assay identified key shear stresses across the surface suggesting an equilibrium behaviour in which approximately 28-39 complexes per cell are activated, which transitions to 93 complexes at 3000 rpm and 127 at 4000 rpm. However, whether these are cooperative interactions of the original complexes or really the activation of 93-127 complexes is not yet fully determined and is the subject of further computational analysis.

This study highlighted the transition of catch bond behaviours from the nanoscale to the microscale and how traditionally defined catch bond behaviour that is truly force dependent is likely a cellular, rather than a molecular, feature. This is in line with physiological studies outlining a Dock-Lock-Latch mechanism as one by which *Staphylococcus epidermidis* colonises a host organism, at the cellular scale. However, it was assumed that this phenomenon must exist on the molecular scale of the surface proteins too. Our study posits that truly force dependent catch bond activation of SdrG:Fgβ, exists only on the cellular scale, and derives from the cumulative effect of individual SdrG:Fg complexes in concert with one another, at the nanoscale this behaviour is highly driven by the chemical environment. These investigations shed new light on catch bond activation at multiple length scales and will be essential to the determination of design parameters for scaling these fascinating complexes to the macroscale.

## Online Methods

### Design and expression of SdrG and Fgβ protein constructs

#### Plasmid design

Using SnapGene (v6.02) 4 constructs were designed for this study; (1) N-SdrG, in which SdrG is oriented to be pulled in the native geometry, *i.e.*, from the N-terminus, (2) C-SdrG, in which the SdrG is pulled in the non-native geometry (from the C-terminus), (3) N_mut_-SdrG, in which SdrG is in the native orientation but a mutation is introduced at S338H to prevent binding to fibrinogen, and (4) Fgβ, the 15 amino acid sequence that fits in the SdrG binding domain, all sequences are given in Table **S1.1**. Three additional sequences were added to the pulling terminus of each construct, the entire composition of which is provided in the electronic supplementary information (Table **S1.1**); (1) His-Tag, to simplify purification, (2) ddFLN4, to improve accuracy in SMFS data analysis, and (3) yBBR, to facilitate surface functionalisation of the AFM cantilever.

#### Protein expression and purification

The designed SdrG constructs were cloned in a pET28 expression vectors by Gibson Assembly using a BioXP 3250 (Telesis Bio, CA, USA), transformed in Top 10 cells (New England Biolabs, MA, USA). After sequence verification with Sanger sequencing (MicroSynth, CH), the constructs were transformed again in *E. coli* BL21 cells (New England Biolabs), then cultured in 500 ml Luria Bertani (LB) medium (10 g/L tryptone, 5 g/L yeast extract, 10 g/L NaCl, all Sigma Aldrich, Buchs, CH, in 1 L milli-Q UltraPure water, Millipore, MA, USA) with 50 μl kanamycin (Thermo Fisher Scientific, Basel CH) at 37°C until an optical density at 600 nm (OD_600_) of 0.7 had been reached. Expression was then induced with 300 μM isopropyl β-D-1-thiogalactopyranoside (IPTG, Thermo Fisher Scientific) and the temperature reduced to 20°C for 20 h.

Expressed proteins were centrifuged in 1 L Nalgene flasks at 8,000 rpm using an RC6+ centrifuge (Thermo Fisher Scientific) for 30 min to precipitate the cells. The supernatant was decanted. Cell pellets were resuspended in tris-buffered saline(TBS) containing 1 mM CaCl_2_ (TBS/Ca buffer) (24.228 g/L tris-base, 87.66 g/L NaCl, 11.09 mg/L CaCl_2_, adjusted to pH 7.2 with 6N HCl, all Sigma Aldrich) and lysed using an ultrasonic probe (Digital Sonifier, Branson, UK) at 25% amplitude for 2 s on and 2 s off, for a total of 5 min active sonication. The sonicated mixture was centrifuged again in 50 ml Nalgene flasks at 15,000 rpm for 30 min to pellet the cell debris and recover the lysate. Each 50 ml portion of cell lysate was mixed with 3 ml of a 50 wt.% mixture of Ni-NTA resin (Thermo Fisher Scientific) in water and shaken on ice for 1 h. The mixture was centrifuged for 5 min at 500 rpm (5920 R, Eppendorf, NJ, USA) in 50 ml Falcon tubes to precipitate the protein bound resin and the supernatant was recovered. The process was repeated with the supernatant, two further times. After each resin recovery, the resin was re-suspended in TBS/Ca buffer and shaken on ice for a further 1 h to remove any unbound protein. After centrifugation, the washed resin was transferred to a 5 ml polypropylene tube with a frit (Gravity Flow Purification Kit, Pierce, Thermo Fisher Scientific), and the washing solution was filtered through the resin bed with bound protein. A top frit was inserted into the column to stabilise the bed, and the resin was washed with 3 column volumes (CVs) of TBS/Ca buffer containing 5 mM imidazole (34 mg/L, Sigma Aldrich) and then with 3 CVs of TBS/Ca buffer containing 10 mM imidazole (68 mg/L) to remove any non-specifically bound proteins. The column was eluted using 5 CVs of TBS/Ca buffer containing 250 mM imidazole (17.02 g/L), followed by at least 1 CV of TBS/Ca buffer.

The column washing was transferred to a VivaSpin 20 ml centrifuge concentrator (Cytiva, Grens, CH) with 10,000 Da molecular weight cut-off (MWCO) and concentrated to less than 1 ml volume. The concentrate was then diluted up to 20 ml at least once with TBS/Ca buffer, and concentrated down to less than 1 ml again, to ensure the final volume of imidazole in the purified product was less than 1 mM. To ensure a high purity product, the concentrated protein was purified using size exclusion chromatography (SEC) with an ÄKTA Pure SEC from GE Healthcare (Chicago, IL, USA) with TBS/Ca as the running buffer and re-concentrated using spin columns, as before. The presence of the desired protein in the fractions of the SEC was verified with SDS-PAGE using the Mini-PROTEAN electrophoresis kit from BioRad (CA, USA) and in-house prepared acrylamide/bisacrylamide gels of 4 wt.% (stacking) and 12 wt.% (separation). After concentration, the proteins were diluted 1:2 with a 50 v/v % glycerol/MQ water solution and stored at -20°C before use.

### Single molecule force spectroscopy

#### Surface functionalisation with SdrG constructs and Fgβ peptide

Surface functionalisation before AFM experiments follows a three-step procedure described previously by this group.^41^ Briefly, 25 mm diameter round glass slides (Menzel Gläser, Braunschweig, Germany) and AFM cantilevers (Biolever Mini AC40TS, Olympus, Tokyo, Japan) are cleaned before use. AFM cantilevers were treated with an ultraviolet ozone cleaner (Novascan, IA, USA) for 40 min, while glass slides were incubated in piranha solution (1:1 (v/v) 30% H_2_O_2_: concentrated H_2_SO_4_) for 30 min. The cleaned surfaces are immersed in a 50 vol.% solution of 3-aminopropyltriethoxysilane (APTES, ABCR, Karlsruhe, Germany) in isopropanol with 0.5 vol.% H_2_O for 5 min. Treated cantilevers were washed with pure toluene, pure ethanol and pure water for 10 s in succession, while treated glass slides were washed 3 times in water. The treated surfaces were placed in an oven at 80°C for 1 h to anneal. After annealing, the surfaces were incubated in a 125 mg/ml solution of a 5,000 Da maleimide-poly(ethylene glycol)-N-hydroxysuccinimide (Mal-PEG5000-NHS, Rapp Polymere, Tübingen, Germany) linker in 4-(2-hydroxyethyl)-1-piperazineethanesulfonic acid buffer (HEPES, 50 mM HEPES, 150 mM NaCl, all Sigma Aldrich) buffer at pH 7.5 for 1 h at RT. The surfaces were washed with ultra-pure water and dried with compressed air. The exposed maleimide groups on the cantilevers and glass slides were reacted with a 200 μM solution of coenzyme A (CoA) in Mal-coupling buffer (50 mM Na_2_HPO_4_, 50 mM NaCl, 10 mM C_10_H_16_N_2_O_8_, adjusted to pH 7.2) for 1 h at RT to facilitate the sulfhydryl reaction at the maleimide chain terminus. Surfaces are again washed in ultrapure water and dried with compressed air. For the AFM cantilevers no further preparation is necessary before the final protein immobilisation stage, as described below. In the case of the cover glasses, slides are attached to a standard microscope slide (VWR) using a 3D printed well to allow measurement under fluid. The last step is the reaction of the ybbR-tagged protein construct with 1.5 μM 4’-phosphopantetheinyl transferase (sfp) and 10 mM MgCl_2_ in TBS/Ca buffer for 2 h at RT to facilitate the enzymatic cross-linking of the protein to the CoA.^52,53^ The surfaces are then washed in the TBS/Ca buffer and stored in the same buffer at 4°C and used as quickly as possible (maximum 3 days of storage, before hydrolysis of the maleimide linkage is observed).

#### Glass slide functionalisation with Fibrinogen

All AFM-SMFS experiments were performed with the SdrG on the AFM cantilever, as above. Experiments performed on Fg surfaces were done using a condensation reaction due to the lack of yBBR handle on the full Fg chain. For the condensation reaction, epoxysilane functionalisation of glass slides was required in place of aminosilane functionalisation. The cleaning and silanisation procedure is the same as above, with the exception that APTES was replaced by (3-glycidyloxypropyl)trimethoxysilane (ABCR). After epoxidation of the glass, 90 μl of a 0.01 mg/ml solution of fibrinogen (Human fibrinogen, plasminogen, vWR and fibronectin depleted, Milan Analytica AG, Rheinfelden, CH) in 0.1 M HEPES buffer at pH 8 was sandwiched between two epoxidised glass slides and incubated for 45 min. The slides were then washed with MQ water, dried with compressed air, and placed on a support inside a petri dish. 600 μl of a 5 wt.% solution of bovine serum albumin (BSA) in phosphate buffered saline (PBS, 8 g/L NaCl, 0.2 g/L KCl, 1.44 g/L Na_2_HPO_4_, 0.24 g/L KH_2_PO_4_ in MQ water, pH adjusted to 7.2) was added to each surface to coat any unfunctionalised epoxy residues on the glass surface. Substrates are stored in TBS/Ca buffer until further use, which is always within 48 h of preparation.

#### SMFS protocol

Single molecule force spectroscopy was performed on a Force Robot AFM (JPK Instruments, Berlin, Germany). Cantilevers were calibrated used a contact free method, with spring constants ranging from 0.1-0.19 N/m. The cantilever was brought into contact with the surface at a force setpoint of 0.18 nN for 0.2 s and then retracted at constant speeds of 400, 800, 1600 and 3200 nm/s. All measurements are collected in liquid at a constant temperature of 22°C. Between retractions the cantilever moves horizontally over the surface by 100 nm until 24 curves at each pulling speed have been attempted. In the absence of interactions over 24 attempts, the surface is moved by 1000 nm. These movements are programmed using an in-house designed script based on Python. A typical experiment was run for 16 h yielding up to 20,000 measurements over this period. TBS/Ca was the measurement buffer used for all SMFS experiments, with the concentration of Ca varied per measurement as described in the text.

#### SMFS Data Analysis

The SMFS data is extracted and smoothed using the JPK Data Analysis software and exported as a .txt file into an in-house designed software (Biomechanical Analyser, BMA) for peak picking and analysis of the rupture forces and contour lengths. After the spectra have been visually assessed they are exported as .tsv files for statistical analysis using python, R Studio and Matlab. This data processing is carried out as described in detail by Boult, et al.^11^

### Yeast expression and display

#### N-SdrG, C-SdrG and N_mut_-SdrG sequence cloning

The gene coding for each of the three SdrG constructs was obtained from the pET28a plasmids described above and optimised for expression in *S. cerevisiae* using SnapGene v6.02. The sequence was cloned by Gibson assembly into a pCHA plasmid (N-terminus constructs N-SdrG and N_mut_-SdrG) or after restriction digestion with NheI and PmeI into a pYD1 plasmid (C-terminus construct, C-SdrG). The sequence was then confirmed by Sanger sequencing. Plasmids pCHA-N-SdrG, pCHA_N_mut_-SdrG and pYD1_C-SdrG were then transformed through a lithium acetate transformation protocol^54^ in the yeast strain EBY100 and the positive colonies selected on synthetic defined (SD) agar 2% (wt/vol) glucose plates lacking tryptophan (-Trp). The sequence of the three yeast plasmids can be found in the ESI S4.

#### Expression and surface display of N-SdrG, C-SdrG and N_mut_-SdrG

Yeast colonies were pre-cultured in -TRP liquid medium with 2 wt.% glucose for 24 h at 30°C to an OD_600_ = 8 with continuous shaking at 224 rcf in an incubator/shaker (Innova 44, New Brunswick Scientific, NJ, USA). Protein expression and display were induced by transferring the culture at a starting OD_600_ = 0.4 to fresh liquid medium containing 0.2 wt.% glucose, 1.8 wt.% galactose and 100 mM citrate/phosphate buffer (4.7 g/L C_6_H_8_O_7_ and 9.2 g/L Na_2_HPO_4_, all Sigma Aldrich, adjusted to pH 3, 5 and 7 to optimise culture, pH 7 was eventually selected see Figure S5.3) to facilitate the best expression conditions. Yeast cells were expressed at 20°C under continuous shaking (224 rcf) for 48 h, before being pelleted by centrifugation at 3920 rcf for 4 min. (5920 R, Eppendorf, NJ, USA). Cells were washed with PBS containing 0.1 wt.% BSA (PBS 0.1 wt.% BSA), re-suspended in 0.5 ml of the same buffer and stored at 4°C until required.

#### Antibody staining for surface displayed N-SdrG, C-SdrG and N_mut_-SdrG

After display, a volume of each of the resuspended cells containing 2M total cells was transferred from each of the storage vials to a separate clean 1.5 ml Eppendorf tube. The tubes were centrifuged at 13,000 rpm for 1 min (5420, Eppendorf, NJ, USA) to precipitate the cells. The supernatant from each tube was then removed. Cells were then incubated at RT for 30 min in 100 μl of a 0.2 μg/μl solution of a primary anti-6x-His mouse monoclonal antibody (Invitrogen, Waltham, MA, USA) in PBS 0.1 wt.% BSA. After incubation, cells were washed twice with PBS 0.1 wt.% BSA and re-suspended at the same concentration of a secondary goat anti-mouse antibody conjugated with Alexa Fluor 594 (Invitrogen). Cells were incubated at 4°C for 30 min. Following incubation, cells were washed twice with cold PBS 0.1 wt.% BSA and re-suspended to achieve a final cell concentration of 2 M cells/300 µl of buffer. Surface expression of each of the stained constructs was confirmed by flow cytometry on an Attune NxT flow cytometer (Thermo Scientific) equipped with 488 and 561 nm lasers. Attune NxT software v.3.2.1 (Life Technologies) was used to analyse the cell populations of interest. Measurements were performed in triplicate, and 10,000 cells were recorded and analysed per construct.

### Spinning Disk Adhesion Assays

#### Protocol

After staining to confirm expression a stock solution of cells at a concentration of 3 M SdrG-displayed cells per 300 μl in TBS/Ca measurement buffer with the appropriate Ca^2+^ concentration. Functionalised glass surfaces, prepared as described in the SMFS section above, were placed into in-house designed metal coverglass holders and seeded with 300 μl of a stock cell solution in measurement buffer and incubated for 30 mins. After incubation, the cell solutions were carefully removed and replaced with 300 μl measurement buffer until spinning, which took place up to 45 min later. The buffer was removed before spinning, and the slide transferred carefully from the metal holder onto the measurement plate on the spinning disk microscope. The cell-laden disks were spun at 3000 or 4000 rpm with an initial ramp rate of 100 rpm/s, holding at the target speed and then ramping down at 100 rpm/s. The entire protocol, including ramping, lasted 308 s regardless of target speed, thus the hold step lasted 20 s less at 4000 rpm than at 3000 rpm. After spinning, glass slides were carefully transferred back to the slide holder and 300 μl of measurement buffer was added. The surface was imaged using a macro within the CellSens software that allows for multiple images across the surface to be stitched, to visualise the cell adhesion distribution across the complete functionalised glass surface.

#### Data Analysis

Numbers of adhered cells and their coordinates were extracted from microscope images using PyCharm and a segmentation script updated for use with Python but previously described by Santos et al.^36^ Both the number and area of the cells was extracted along with their coordinates. This data was processed using an in-house developed computational analysis workflow. This consists of (1) localising the coordinates of the cells from n=3+ measurements on a combined density plot to determine the profile of the cell distribution over the cover slide, (2) using the relative distribution of the cells from the origin to create a histogram of distribution, (3) using the results from 1 and 2 to determine if the plots exhibit attributes of a catch bond forming complex, (4) plotting median cells per bin against shear rate, (5) fitting the median cells vs shear rate with a sigmoidal fit (slip bonds), a combined sigmoidal/gaussian fitting function or a double gaussian, and finally (6) obtaining the first and second derivative of the sigmoidal/gaussian fit to extract the inflection and turning points that correspond to the onset, and maxima of the catch bond. All Jupyter notebooks containing raw and analysed data, including the fitting protocol and extraction of the catch bond activation and rupture forces are available on GitHub.

## Data availability

Datasets will be made available for download upon request.

## Code availability

Python scripts developed for the analysis and visualisation of data will be made available upon request.

## Acknowledgements

Z.W.K. acknowledges funding from the European Union’s Horizon 2020 research and innovation programme under the Marie Skłodowska-Curie grant agreement No. 842043. IUA and IAT acknowledge financial support from the Research Ireland grant no. 21/RC/10295_P2 and from Erasmus+ International Credit Mobility programme 2022-1-IE02-KA171-HE-000073430. This work was further supported by the University of Basel, ETH Zurich, the SNF-NCCR in Molecular Systems Engineering, and an SNSF Grant (200021_191962) to M.A.N.

## Author Affiliations

*Zarah Walsh–Korb*

Department of Chemistry, University of Basel, Mattenstrasse 22, 4058 Basel, Switzerland *and* Department of Biomedical Engineering, University of Basel, Hegenheimermattweg 167B, 4123 Allschwil, Switzerland

*Sean Boult and Rosario Vanella*

Department of Chemistry, University of Basel, Mattenstrasse 22, 4058 Basel, Switzerland *and* Department of Biosystems Science and Engineering, ETH Zurich, Klingelbergstrasse 48, 4056 Basel, Switzerland

*Vanni Doffini, and Michael A. Nash*

Department of Chemistry, University of Basel, Mattenstrasse 22, 4058 Basel, Switzerland *and* Department of Biosystems Science and Engineering, ETH Zurich, Klingelbergstrasse 48, 4056 Basel, Switzerland *and* Swiss Nanoscience Institute, Klingelbergstrasse 86, 4056 Basel, Switzerland

*Inam Ul Ahad*

DCU Institute of Advanced Processing Technology, School of Mechanical & Manufacturing Engineering, Dublin City University, Glasnevin Campus, Dublin 9, D09 V209, Ireland

*Intizar Ali Tunio*

School of Mechanical & Manufacturing Engineering, Dublin City University, Glasnevin Campus, Dublin 9, D09 V209, Ireland *and* Department of Mechanical Engineering, Mehran University of Engineering & Technology, Jamshoro, Pakistan

## Contributions

**ZWK** - conceptualisation, methodology, validation, formal analysis, resources, data curation, writing (original draft), visualisation, project administration, supervision, funding acquisition; **SB** - methodology, software, formal analysis; **JL** - methodology, software; **IAT** - methodology, software, formal analysis, investigation, visualisation; **RV** - methodology, investigation; **VD** - methodology, software, formal analysis; **IUA** - supervision, funding acquisition; **MAN** - conceptualisation, supervision, resources, funding acquisition. All authors reviewed and edited the manuscript.

## Ethics declarations

The authors declare no competing interests.

