## Supplemental Information for "Probing the scalability of ultra stable catch bond complexes"

### Electronic Supplementary Information for “Probing the scalability of ultra stable catch bond complexes”

The following sections provide additional information on the preparation of the protein constructs, additional SMFS data, an expanded dataset from the Monte Carlo simulations, an expanded dataset of SDA assays, information on yeast expression and display, and computation fluid dynamics experimental set-up.

#### S1. Plasmid maps for SMFS protein constructs

Plasmid maps from SnapGene for the 4 engineered protein constructs (N-SdrG, C-SdrG, $N_{mut}$-SdrG and Fgβ) for the SMFS experiments are shown in Figures **S1.1**-**S1.4**.

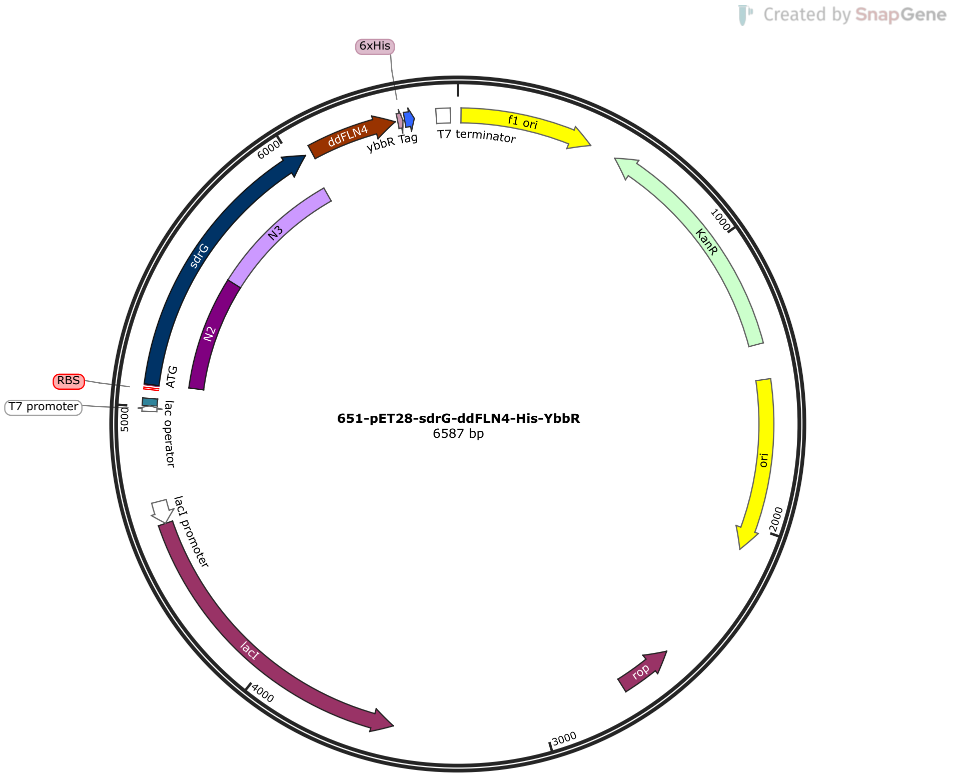


**Figure S1.1:** Plasmid map for the N-SdrG construct with ddFLN4, His-tag and yBBR domains in a pET28 expression vector created in SnapGene

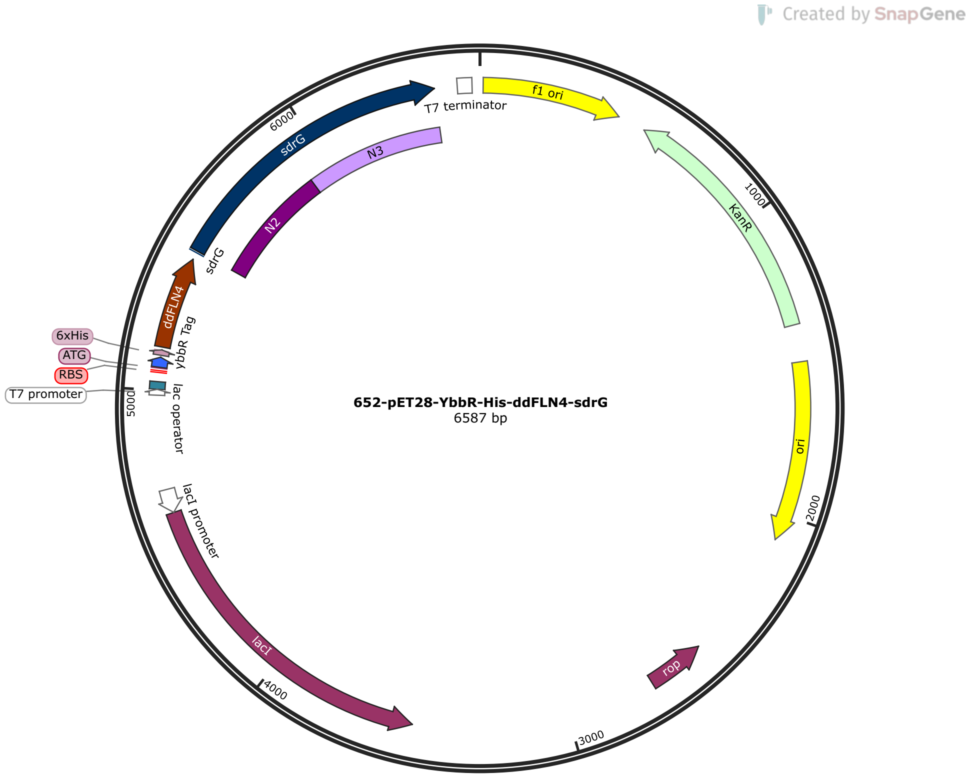


**Figure S1.2:** Plasmid map for the C-SdrG construct with ddFLN4, His-tag and yBBR domains in a pET28 expression vector created in SnapGene

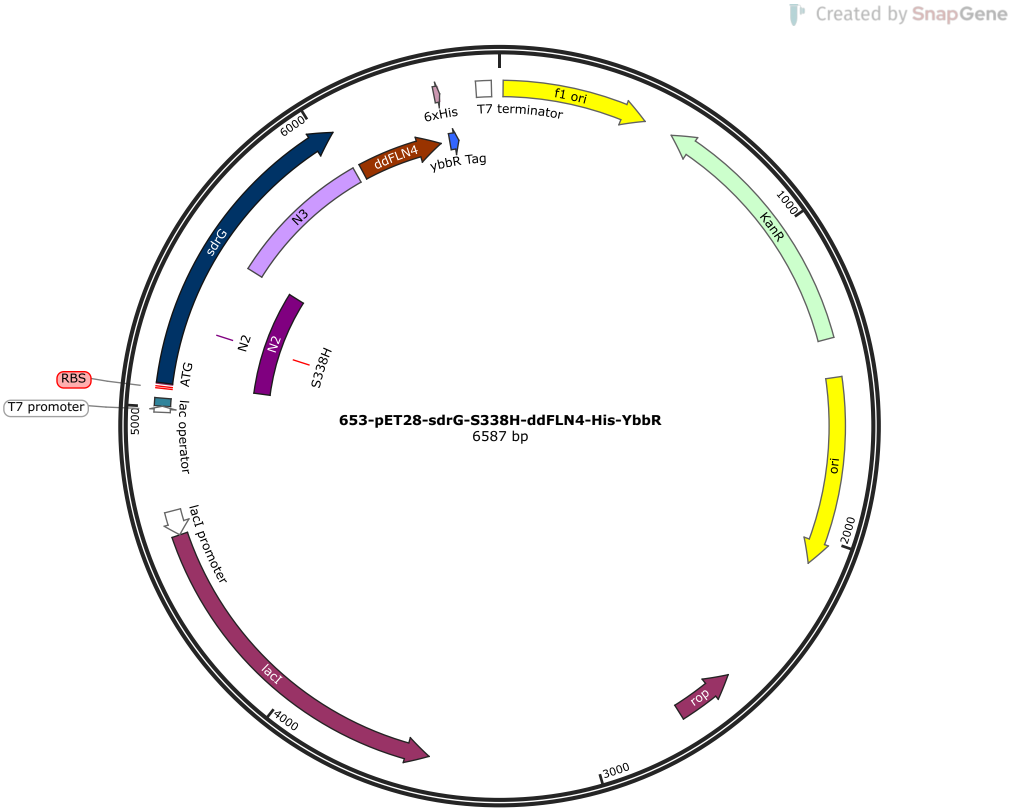


**Figure S1.3:** Plasmid map for the N$_{mut}$-SdrG construct with ddFLN4, His-tag and yBBR domains in a pET28 expression vector created in SnapGene

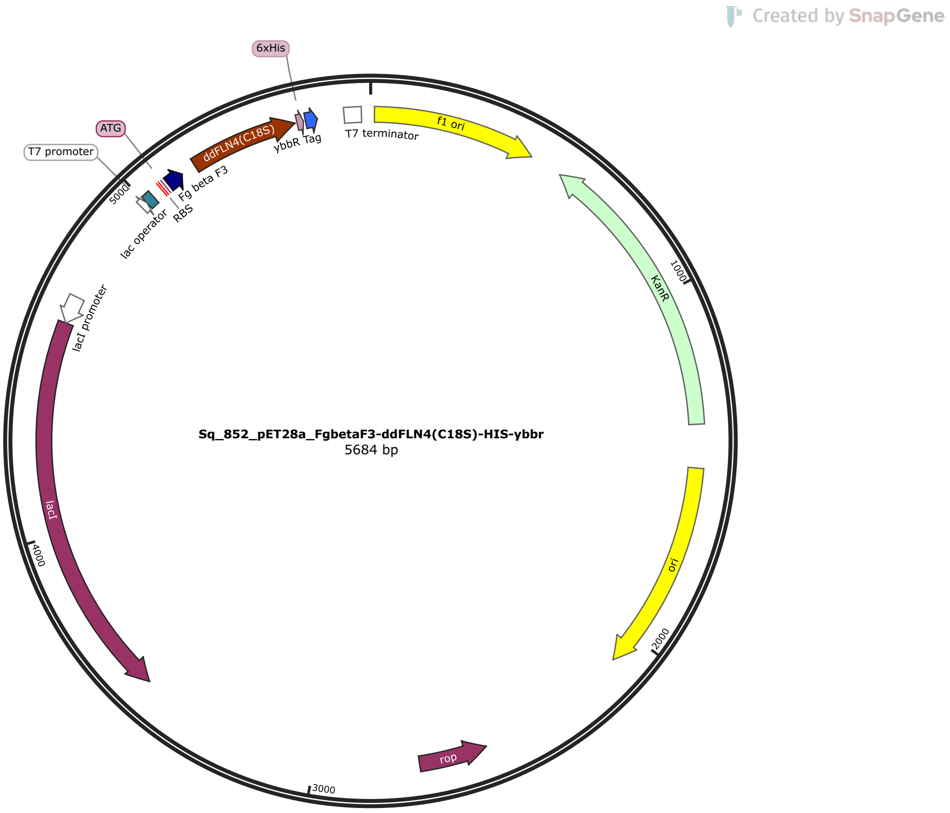


**Figure S1.4:** Plasmid map for the Fgβ construct with ddFLN4, His-tag and yBBR domains in a pET28 expression vector created in SnapGene

**Table S1.1:** Amino acid sequences for each part of the protein constructs under investigation

| Fragment | Sequence |
|---|---|
| N-SdrG | MEQGS NVNHL IKVTD QSITE GYDDS DGIIK AHDAE NLIYD VTFEV DDKVK SGDTM TVNID KNTVP SDLTD SFAIP KIKDN SGEII ATGTY DNTNK QITYT FTDYV DKYEN IKAHL KLTSY IDKSK VPNNN TKLDV EYKTA LSSVN KTITV EYQKP NENRT ANLQS MFTNI DTKNH TVEQT IYINP LRYSA KETNV NISGN \par GDEGS TIIDD STIIK VYKVG DNQNL PDSNR IYDYS ETEDV TNDDY AQLGN NNDVN INFGN IDSPY IIKVI SKYDP NKDDY TTIQQ TVTMQ TTINE YTGEF RTASY DNTIA FSTSS GQGQG DLPPE |
| C-SdrG | EQGSN VNHLI KVTDQ SITEG YDDSD GIIKA HDAEN LIYDV TFEVD DKVKS GDTMT VNIDK NTVPS DLTDS FAIPK IKDNS GEIIA TGTYD NTNKQ ITYTF TDYVD KYENI KAHLK LTSYI DKSKV PNNNT KLDVE YKTAL SSVNK TITVE YQKPN ENRTA NLQSM FTNID TKNHT VEQTI YINPL RYSAK ETNVN ISGNG DEGST IIDDS TIIKV YKVGD NQNLP DSNRI YDYSE TEDVT NDDYA QLGNN NDVNI NFGNI DSPYI IKVIS KYDPN KDDYT TIQQT VTMQT TINEY TGEFR TASYD NTIAF STSSG QGQGD LPPE |
| $N_{mut}$-SdrG | MEQGS NVNHL IKVTD QSITE GYDDS DGIIK AHDAE NLIYD VTFEV DDKVK SGDTM TVNID KNTVP HDLTD SFAIP KIKDN SGEII ATGTY DNTNK QITYT FTDYV DKYEN IKAHL KLTSY IDKSK VPNNN TKLDV EYKTA LSSVN KTITV EYQKP NENRT ANLQS MFTNI DTKNH TVEQT IYINP LRYSA KETNV NISGN \par GDEGS TIIDD STIIK VYKVG DNQNL PDSNR IYDYS ETEDV TNDDY AQLGN NNDVN INFGN IDSPY IIKVI SKYDP NKDDY TTIQQ TVTMQ TTINE YTGEF RTASY DNTIA FSTSS GQGQG DLPPE |
| Fgβ | NEEGF FSARG HRPLD |
| His-tag | HHHHH H |
| ddFLN4 | ADPEK SYAEG PGLDG GECFQ PSKFK IHAVD PDGVH RTDGG DGFVV TIEGP APVDP VMVDN GDGTY DVEFE PKEAG DYVIN LTLDG DNVNG FPKTV TVKPA P |
| yBBR | DSLEF IASKL A |

#### S2. AFM-Single molecule force spectroscopy

AFM-single molecule force spectroscopy (AFM-SMFS) was performed on three representative surfaces under each condition, and the experiment was allowed to run for approximately 16 h. Data from each experiment was aggregated and analysed together to give the datasets shown in the following figures.

##### *S.2.1 N-SdrG:Fgβ expanded rupture force data*

These experiments are conducted to understand the effect of the $Ca^{2+}$ concentration on the specific catch bond interaction. The experiment is performed at four pulling speeds simultaneously – 400, 800, 1600, and 3200 nm/s using an N-SdrG-coated cantilever and an Fgβ-coated substrate in 1 mM (Figure **S2.1**), 2.5 mM (Figure **S2.2**), 6.25 mM (Figure **S2.3**), and 10 mM (Figure **S2.4**) $Ca^{2+}$ in TBS buffer.

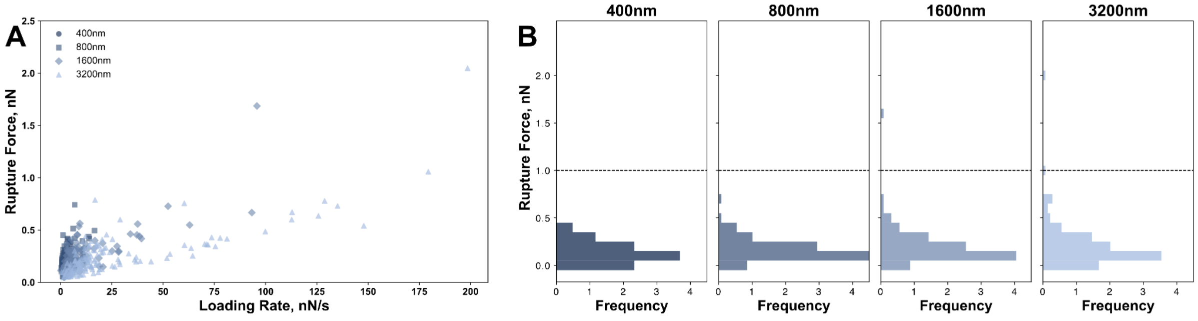


**Figure S2.1:** (**A**) Scatter plot of loading rate (nN/s) vs rupture force (nN) for each of the 4 pulling speeds (400, 800, 1600, 3200 nm/s), and (**B**) histograms of frequency vs rupture force at each pulling speed for an N-SdrG-coated cantilever against an Fgβ-coated substrate in TBS buffer with 1 mM $Ca^{2+}$

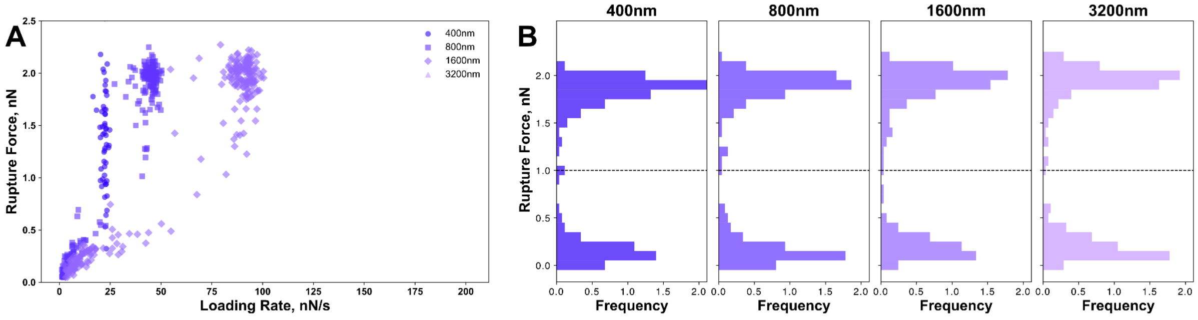


**Figure S2.2:** (**A**) Scatter plot of loading rate (nN/s) vs rupture force (nN) for each of the 4 pulling speeds (400, 800, 1600, 3200 nm/s), and (**B**) histograms of frequency vs rupture force at each pulling speed for an N-SdrG-coated cantilever against an Fgβ-coated substrate in TBS buffer with 2.5 mM $Ca^{2+}$

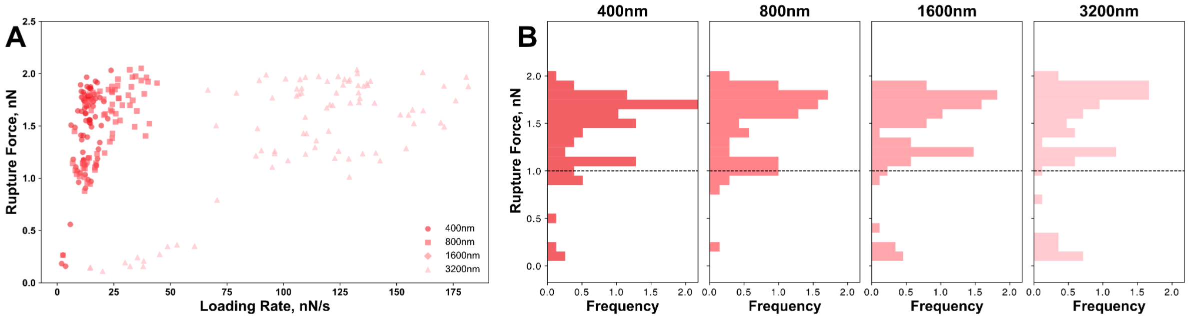


**Figure S2.3:** (**A**) Scatter plot of loading rate (nN/s) vs rupture force (nN) for each of the 4 pulling speeds (400, 800, 1600, 3200 nm/s), and (**B**) histograms of frequency vs rupture force at each pulling speed for an N-SdrG-coated cantilever against an Fgβ-coated substrate in TBS buffer with 6.25 mM $Ca^{2+}$

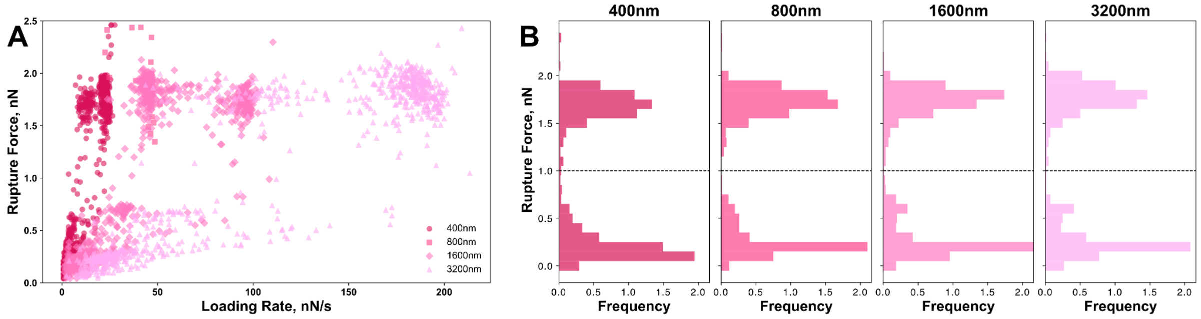


**Figure S2.4:** (**A**) Scatter plot of loading rate (nN/s) vs rupture force (nN) for each of the 4 pulling speeds (400, 800, 1600, 3200 nm/s), and (**B**) histograms of frequency vs rupture force at each pulling speed for an N-SdrG-coated cantilever against an Fgβ-coated substrate in TBS buffer with 10 mM $Ca^{2+}$

#### *S.2.2 N-SdrG:Fg expanded rupture force data*

These experiments are conducted to understand both the effect of the $Ca^{2+}$ concentration on the catch bond interaction and how the specificity of the substrate impacts this behaviour. The experiment is performed at four pulling speeds simultaneously – 400, 800, 1600, and 3200 nm/s using an N-SdrG-coated cantilever and an Fg-coated substrate in 1 mM (Figure **S2.5**), 6.25 mM (Figure **S2.6**), and 10 mM (Figure **S2.7**) $Ca^{2+}$ in TBS buffer.

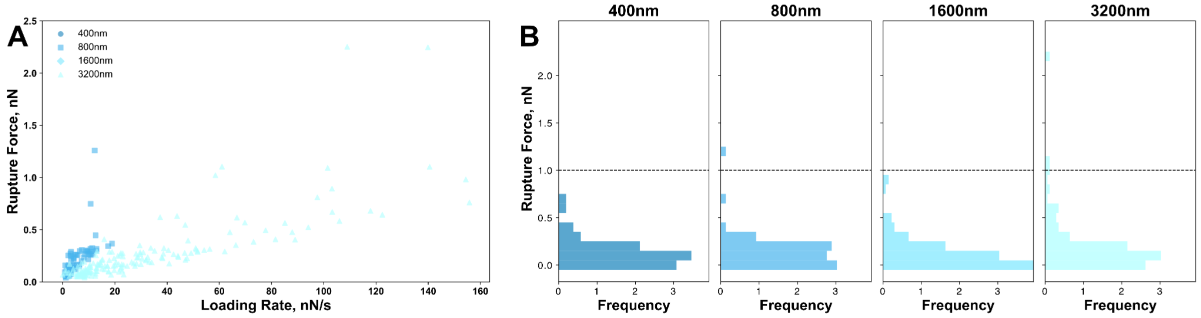


**Figure S2.5:** (**A**) Scatter plot of loading rate (nN/s) vs rupture force (nN) for each of the 4 pulling speeds (400, 800, 1600, 3200 nm/s), and (**B**) histograms of frequency vs rupture force at each pulling speed for an N-SdrG-coated cantilever against an Fg-coated substrate in TBS buffer with 1 mM $Ca^{2+}$

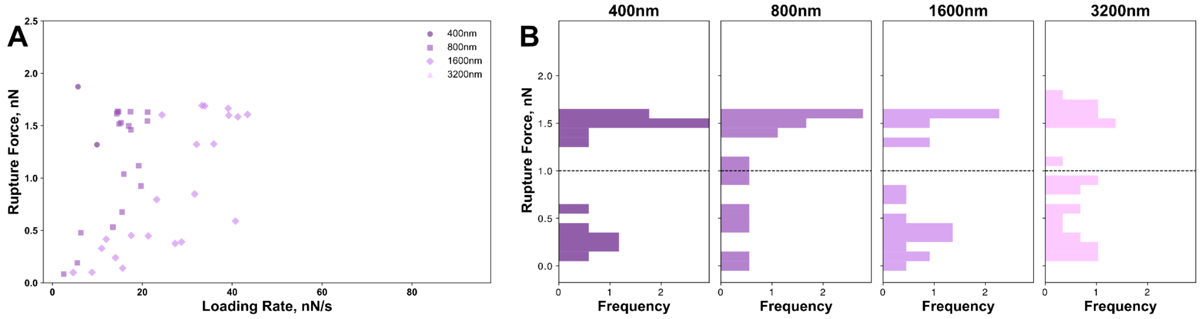


**Figure S2.6:** (**A**) Scatter plot of loading rate (nN/s) vs rupture force (nN) for each of the 4 pulling speeds (400, 800, 1600, 3200 nm/s), and (**B**) histograms of frequency vs rupture force at each pulling speed for an N-SdrG-coated cantilever against an Fg-coated substrate in TBS buffer with 6.25 mM $Ca^{2+}$

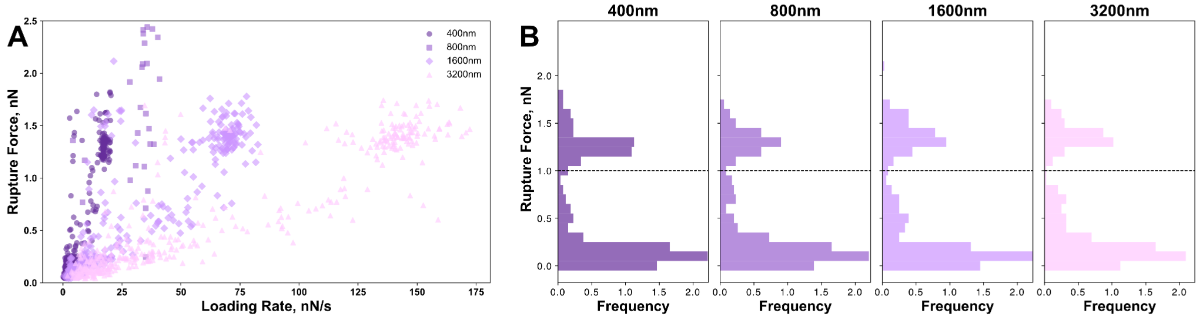


**Figure S2.7:** (**A**) Scatter plot of loading rate (nN/s) vs rupture force (nN) for each of the 4 pulling speeds (400, 800, 1600, 3200 nm/s), and (**B**) histograms of frequency vs rupture force at each pulling speed for an N-SdrG-coated cantilever against an Fg-coated substrate in TBS buffer with 10 mM $Ca^{2+}$

#### *S.2.3 $N_{mut}$-SdrG:Fgβ expanded rupture force data*

These experiments are conducted to understand the effect of the binding pocket mutation on catch bond formation, essentially as a control to ensure that the effects observed in other experiments can be validated. This experiment is performed at four pulling speeds simultaneously – 400, 800, 1600, and 3200 nm/s using an $N_{mut}$-SdrG-coated cantilever and an Fgβ-coated substrate in 10 mM $Ca^{2+}$ in TBS buffer (Figure **S2.8**).

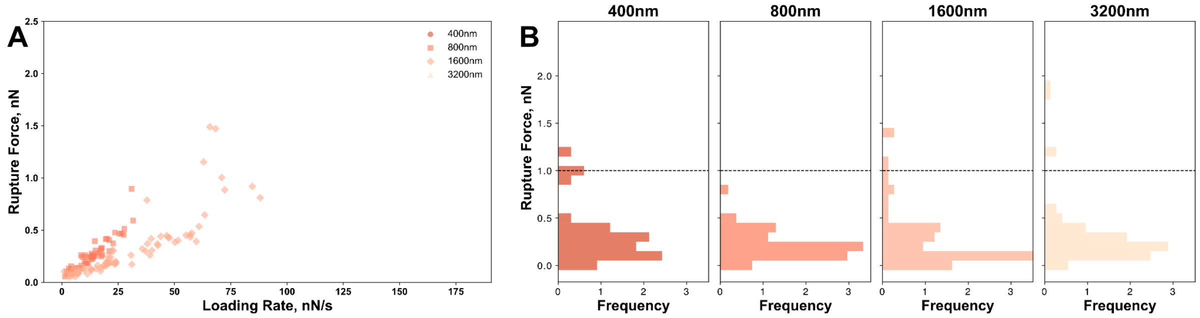


**Figure S2.8:** (**A**) Scatter plot of loading rate (nN/s) vs rupture force (nN) for each of the 4 pulling speeds (400, 800, 1600, 3200 nm/s), and (**B**) histograms of frequency vs rupture force at each pulling speed for an $N_{mut}$-SdrG-coated cantilever against an Fgβ-coated substrate in TBS buffer with 10 mM $Ca^{2+}$

#### *S.2.4 $N_{mut}$-SdrG:Fg expanded rupture force data*

These experiments are conducted to understand the effect of the binding pocket mutation on catch bond formation, essentially as a control to ensure that the effects observed in other experiments can be validated, as well as to determine if this is impacted by the specificity of the substrate. This is performed at four pulling speeds simultaneously – 400, 800, 1600, and 3200 nm/s using an $N_{mut}$-SdrG-coated cantilever and an Fg-coated substrate in 10 mM $Ca^{2+}$ in TBS buffer (Figure **S2.9**).

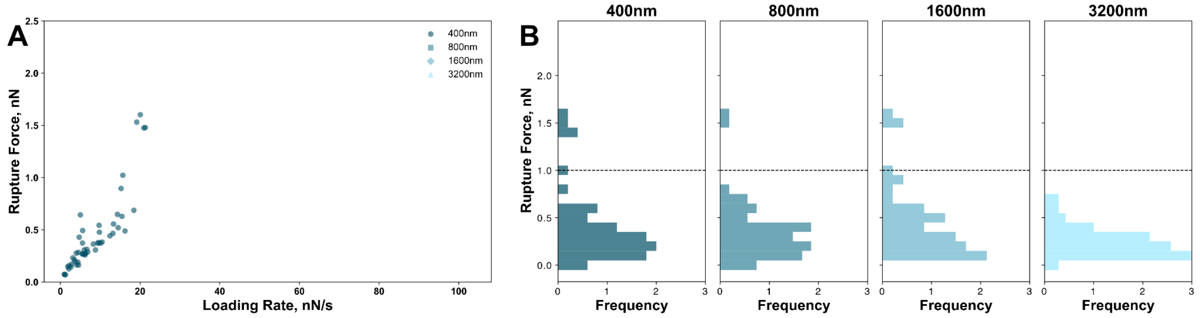


**Figure S2.9:** (**A**) Scatter plot of loading rate (nN/s) vs rupture force (nN) for each of the 4 pulling speeds (400, 800, 1600, 3200 nm/s), and (**B**) histograms of frequency vs rupture force at each pulling speed for an $N_{mut}$-SdrG-coated cantilever against an Fg-coated substrate in TBS buffer with 10 mM $Ca^{2+}$

#### *S.2.5 C-SdrG:Fgβ expanded rupture force data*

These experiments are conducted to understand the effect of the pulling geometry or protein orientation on the cantilever on catch bond formation. Again, this experiment is performed at four pulling speeds simultaneously – 400, 800, 1600, and 3200 nm/s using a C-SdrG-coated cantilever and an Fgβ-coated substrate in 1 mM (Figure **S2.10**), and 10 mM (Figure **S2.11**) $Ca^{2+}$ in TBS buffer.

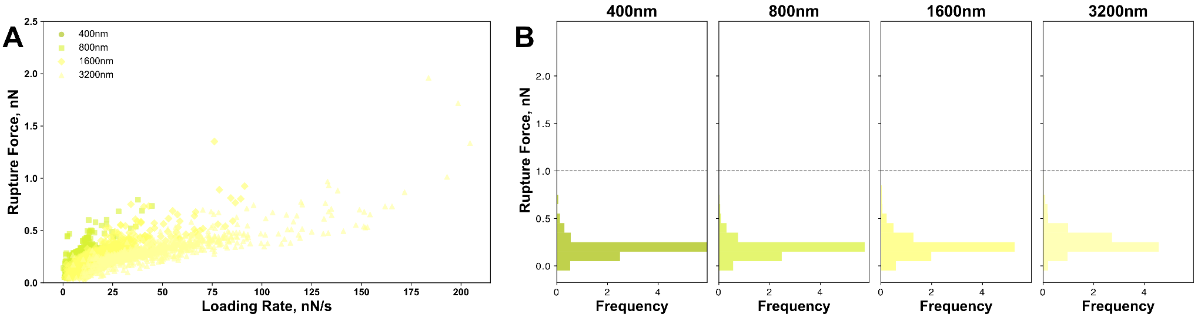


**Figure S2.10:** (**A**) Scatter plot of loading rate (nN/s) vs rupture force (nN) for each of the 4 pulling speeds (400, 800, 1600, 3200 nm/s), and (**B**) histograms of frequency vs rupture force at each pulling speed for a C-SdrG-coated cantilever against an Fgβ-coated substrate in TBS buffer with 1 mM $Ca^{2+}$

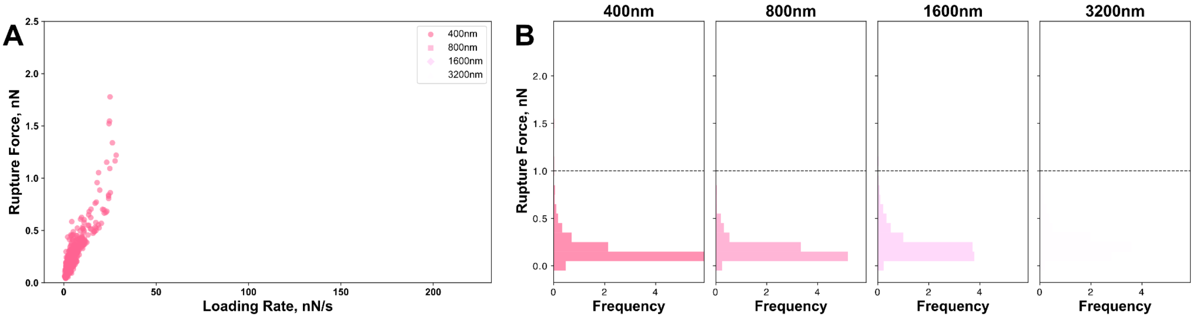


**Figure S2.11:** (**A**) Scatter plot of loading rate (nN/s) vs rupture force (nN) for each of the 4 pulling speeds (400, 800, 1600, 3200 nm/s), and (**B**) histograms of frequency vs rupture force at each pulling speed for a C-SdrG-coated cantilever against an Fgβ-coated substrate in TBS buffer with 10 mM $Ca^{2+}$

#### *S.2.6 C-SdrG:Fg expanded rupture force data*

These experiments are conducted to understand the effect of the pulling geometry or protein orientation on the cantilever on catch bond formation, alongside how this is impacted by the specificity of the substrate. This experiment is performed at four pulling speeds simultaneously – 400, 800, 1600, and 3200 nm/s using a C-SdrG-coated cantilever and an Fg-coated substrate in 1 mM (Figure **S2.12**), and 10 mM (Figure **S2.13**) $Ca^{2+}$ in TBS buffer.

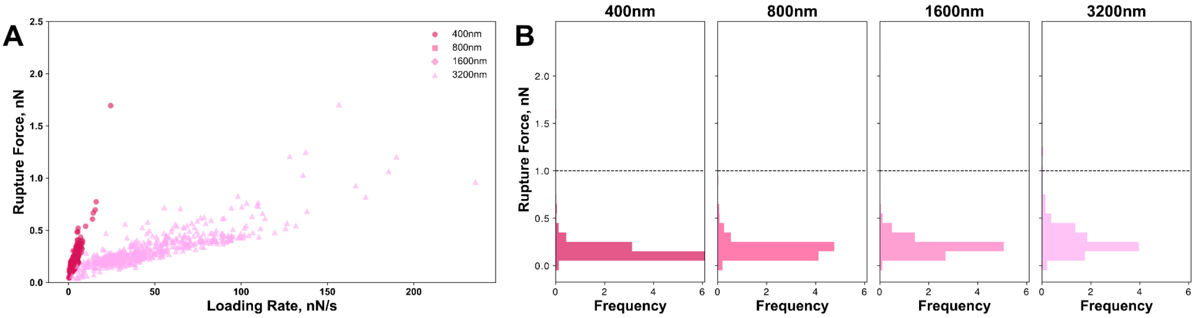


**Figure S2.12:** (**A**) Scatter plot of loading rate (nN/s) vs rupture force (nN) for each of the 4 pulling speeds (400, 800, 1600, 3200 nm/s), and (**B**) histograms of frequency vs rupture force at each pulling speed for a C-SdrG-coated cantilever against an Fg-coated substrate in TBS buffer with 1 mM $Ca^{2+}$

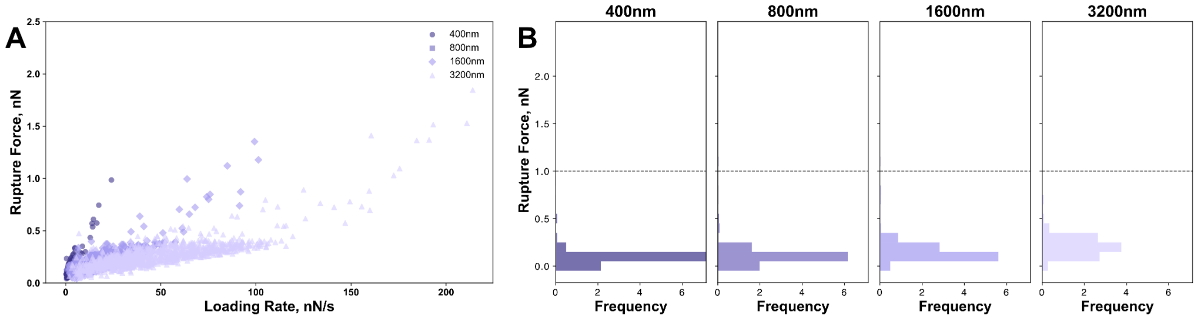


**Figure S2.13:** (**A**) Scatter plot of loading rate (nN/s) vs rupture force (nN) for each of the 4 pulling speeds (400, 800, 1600, 3200 nm/s), and (**B**) histograms of frequency vs rupture force at each pulling speed for a C-SdrG-coated cantilever against an Fg-coated substrate in TBS buffer with 10 mM $Ca^{2+}$

#### S3 Monte Carlo Simulations

Our datasets clearly show the presence of at least two distinct populations under most conditions, particularly when N-SdrG is investigated, indicating that binding kinetics should be understood with a two-pathway model, as for many other catch bonds analysed by this group and others. What is not clear is whether both pathways are accessible to each interaction or whether there are potentially more than two-pathways to follow. To understand more about the binding kinetics and how it might be impacted by the environment, Monte Carlo simulations were performed with 4 different models: reversible transition two-pathway (rttp), in which both pathways are accessible to each interaction, irreversible transition two-pathway (ittp), in which interactions that follow the high force rupture cannot transition back to the low force rupture pathway, multiple binding modes two-pathway (mbmtp), in which it is assumed that there are at least 2 pathways and transitions can be made between them, and finally a Ca-dependent mbmtp. The full raw data associated with Figure 3 of the manuscript are given in the following sections. Scripts for each of these models are given at the resources listed above.

##### *S3.1 Extraction of kinetic data from SMFS datasets*

In order to conduct Monte Carlo simulations, certain parameters must be extracted from the raw SMFS data. Firstly, the ratios of high force to low force rupture events, Table S3.1 (given as an average across loading rates and as a function of pulling speed and $Ca^{2+}$ concentration) must be obtained. As well as the intrinsic dissociation rate constant ($k_{off}$), the rate at which the complex would rupture without any applied force, and the distance to the transition state ($\Delta x$), which is force dependent. These are given as a function of buffer $Ca^{2+}$ concentration, in Tables **S3.2**-**S3.5**. We used two different scripts to extract these parameters; one based on a linear fitting (H in Figures **S3.1**-**S3.16**) and one based on a global fitting (G n Figures **S3.1**-**S3.16**) and evaluated the correlation with the raw datasets.

**Table S3.1**: High force/low force ratios extracted from the SMFS datasets of the N-SdrG:Fgβ interactions using the Bell Evans equation

| **Average High Force/Low Force Ratios** | **1 mM** | **2.5 mM** | **6.25 mM** | **10 mM** |
|---|---|---|---|---|
| | 0.0175 | 0.57 | 0.92 | 0.5738 |
| **Concentration** | **400 nm/s** | **800 nm/s** | **1600 nm/s** | **3200 nm/s** |
| **1 mM** | 0 | 0 | 0.01 | 0.06 |
| **2.5 mM** | 0.6 | 0.56 | 0.57 | 0.55 |
| **6.25 mM** | 0.95 | 0.96 | 0.91 | 0.86 |
| **10 mM** | 0.67 | 0.76 | 0.64 | 0.54 |

**Table S3.2**: Kinetic data extracted from the SMFS datasets of the N-SdrG:Fgβ interactions using the Bell Evans equation at 1 mM $Ca^{2+}$

| Model | Low Force *dx* | Low Force $k_{off}$ | High Force *dx* | High Force $k_{off}$ |
|---|---|---|---|---|
| **linear** | 0.3693443 | $5.366979\times10^{-10}$ | 100 | 100 |
| **Global (G)** | 0.09121784 | 0.6895718 | 0.01249848 | 0.03124917 |
| **in-house (H)** | 0.107857148 | 9.2715227 | 0.0218458632 | 45.77525 |

**Table S3.3**: Kinetic data extracted from the SMFS datasets of the N-SdrG:Fgβ interactions using the Bell Evans equation at 2.5 mM $Ca^{2+}$

| Model | Low Force *dx* | Low Force $k_{off}$ | High Force *dx* | High Force $k_{off}$ |
|---|---|---|---|---|
| **linear** | 0.1343916 | 0.04409844 | 0.08816555 | $5.81047\times10^{-16}$ |
| **Global (G)** | 0.02814467 | 6.671574 | 0.08107483 | $2.666974\times10^{-14}$ |
| **In-house (H)** | 0.117349618171034 | 8.52154455707324 | 0.0306635999284198 | 32.6119569239871 |

**Table S3.4**: Kinetic data extracted from the SMFS datasets of the N-SdrG:Fgβ interactions using the Bell Evans equation at 6.25 mM $Ca^{2+}$

| Model | Low Force *dx* | Low Force $k_{off}$ | High Force *dx* | High Force $k_{off}$ |
|---|---|---|---|---|
| **linear** | -0.007016891 | $-5.769065\times10^{-15}$ | 0.03854874 | $1.339456\times10^{-34}$ |
| **Global (G)** | 0.02350186 | 0.08989391 | 0.1196068 | $2.450007\times10^{-19}$ |
| **In-house (H)** | 0.049222466956076 | 20.3159260768534 | 0.015297546994764 | 65.3699577025178 |

**Table S3.5**: Kinetic data extracted from the SMFS datasets of the N-SdrG:Fgβ interactions using the Bell Evans equation at 10 mM $Ca^{2+}$

| Model | Low Force *dx* | Low Force $k_{off}$ | High Force *dx* | High Force $k_{off}$ |
|---|---|---|---|---|
| **linear** | 0.2060607 | 0.00282463 | 0.1153503 | $1.449808\times10^{-18}$ |
| **Global (G)** | 0.04223003 | 3.233904 | 0.1224587 | $1.89342\times10^{-20}$ |
| **In-house (H)** | 0.0981576900732965 | 10.1876888021028 | 0.0578283405860253 | 17.2925591477487 |

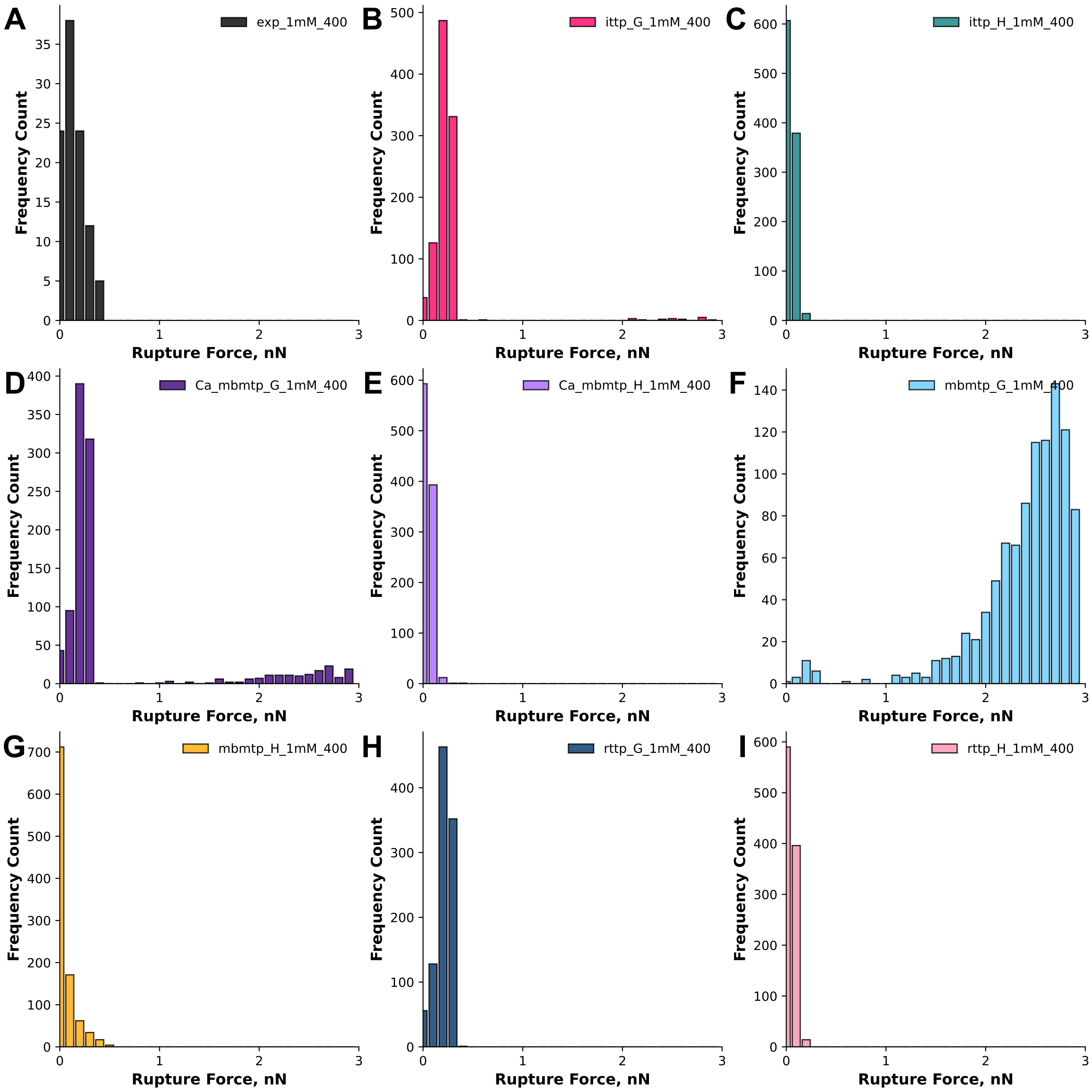


**Figure S3.1:** Histograms of rupture force (nN) vs frequency count for the raw data (black, **A**) obtained from the pulling the N-SdrG-coated cantilever from an Fgβ-coated substrate in TBS with 1 mM $Ca^{2+}$ at 400 nm/s and each of the Monte Carlo models, ittp-global (dark pink, **B**), ittp-linear (green, **C**), Ca-mbmtp-global (dark purple, **D**), Ca-mbmtp-linear (purple, **E**), mbmtp-global (blue, **F**), mbmtp-linear (yellow, **G**), rttp-global (dark blue, **H**), rttp-linear (pink, **I**).

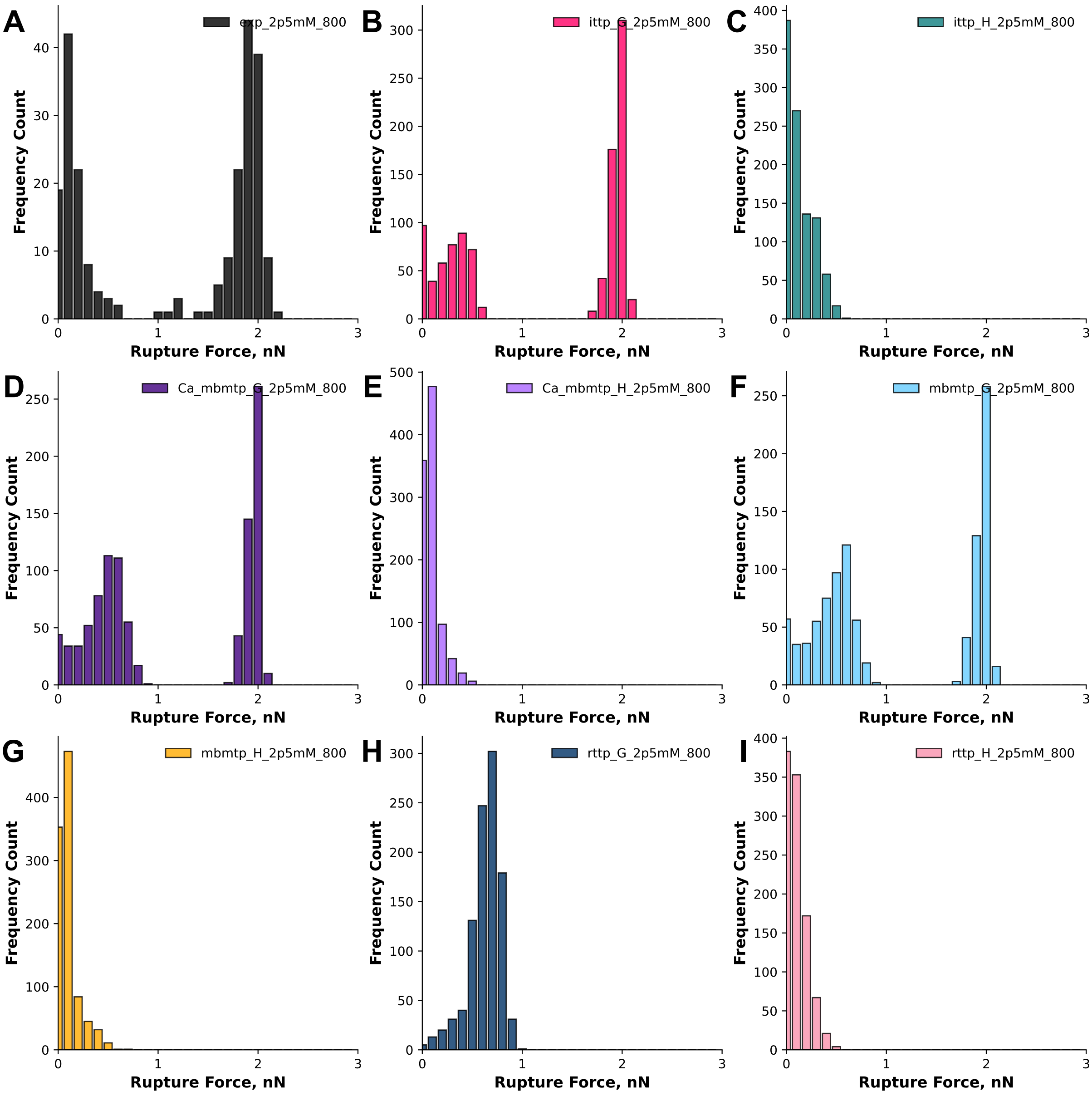


**Figure S3.2:** Histograms of rupture force (nN) vs frequency count for the raw data (black, **A**) obtained from the pulling the N-SdrG-coated cantilever from an Fgβ-coated substrate in TBS with 1 mM $Ca^{2+}$ at 800 nm/s and each of the Monte Carlo models, ittp-global (dark pink, **B**), ittp-linear (green, **C**), Ca-mbmtp-global (dark purple, **D**), Ca-mbmtp-linear (purple, **E**), mbmtp-global (blue, **F**), mbmtp-linear (yellow, **G**), rttp-global (dark blue, **H**), rttp-linear (pink, **I**).

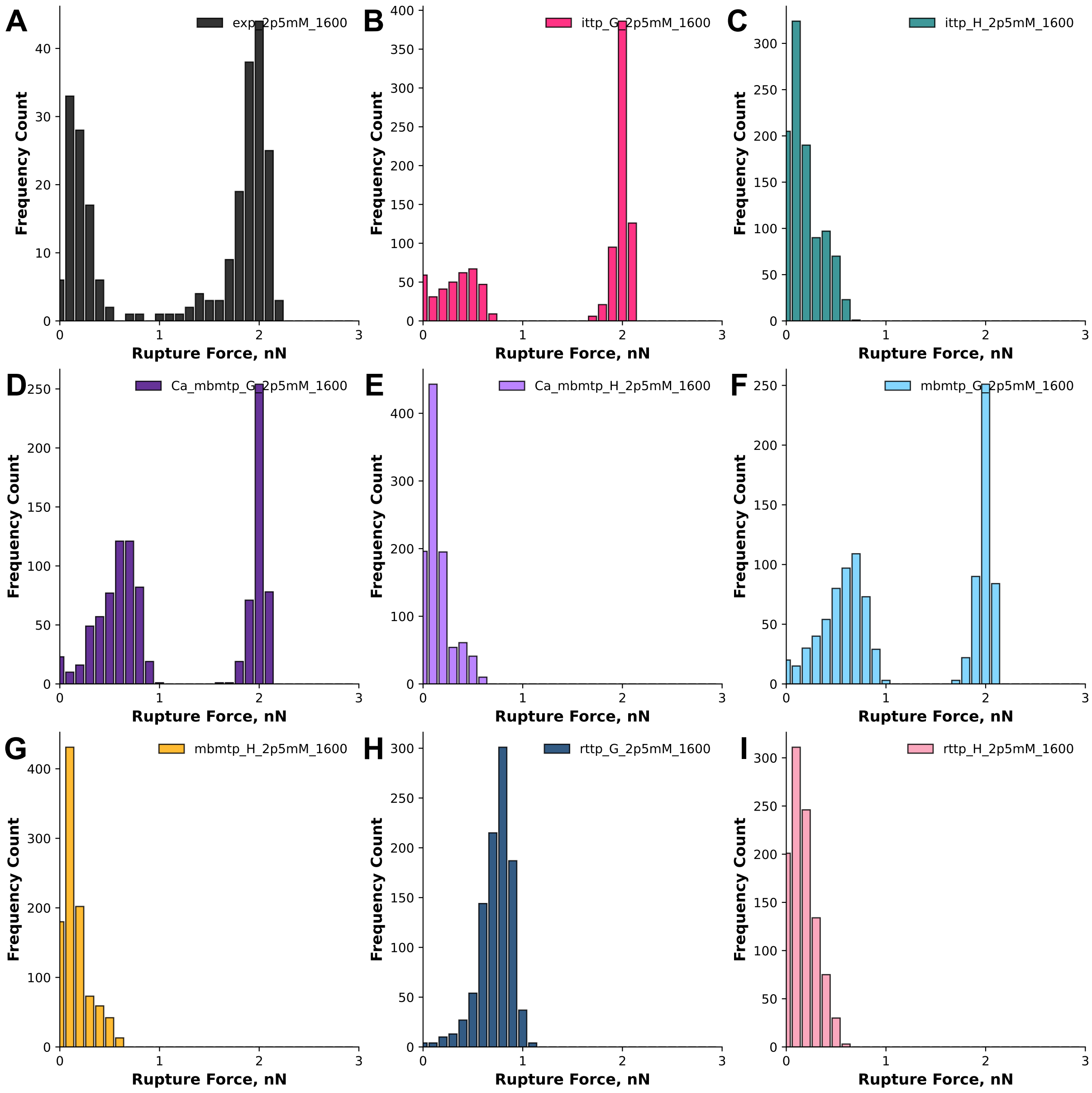


**Figure S3.3:** Histograms of rupture force (nN) vs frequency count for the raw data (black, **A**) obtained from the pulling the N-SdrG-coated cantilever from an Fgβ-coated substrate in TBS with 1 mM $Ca^{2+}$ at 1600 nm/s and each of the Monte Carlo models, ittp-global (dark pink, **B**), ittp-linear (green, **C**), Ca-mbmtp-global (dark purple, **D**), Ca-mbmtp-linear (purple, **E**), mbmtp-global (blue, **F**), mbmtp-linear (yellow, **G**), rttp-global (dark blue, **H**), rttp-linear (pink, **I**).

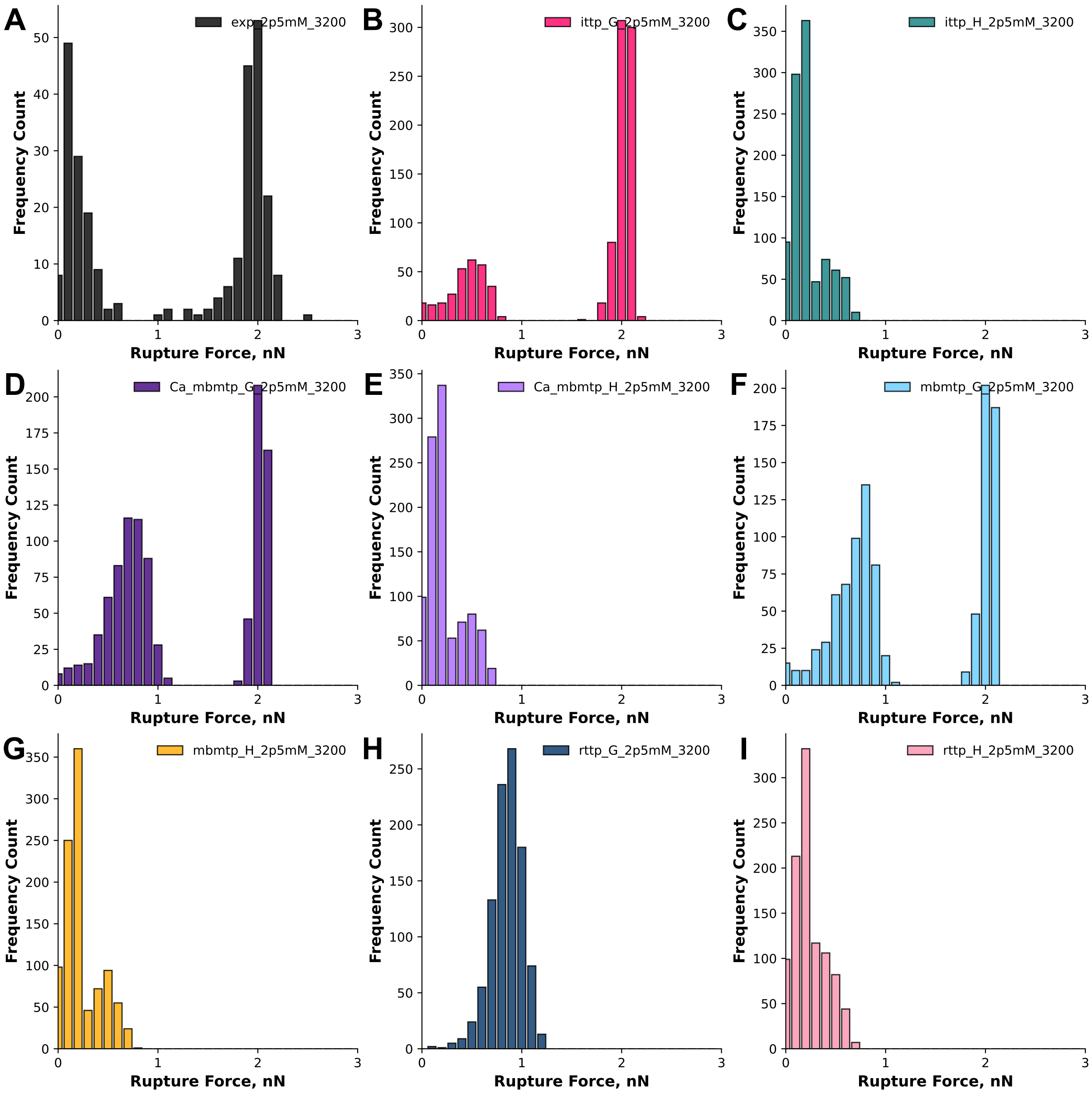


**Figure S3.4:** Histograms of rupture force (nN) vs frequency count for the raw data (black, **A**) obtained from the pulling the N-SdrG-coated cantilever from an Fgβ-coated substrate in TBS with 1 mM $Ca^{2+}$ at 3200 nm/s and each of the Monte Carlo models, ittp-global (dark pink, **B**), ittp-linear (green, **C**), Ca-mbmtp-global (dark purple, **D**), Ca-mbmtp-linear (purple, **E**), mbmtp-global (blue, **F**), mbmtp-linear (yellow, **G**), rttp-global (dark blue, **H**), rttp-linear (pink, **I**).

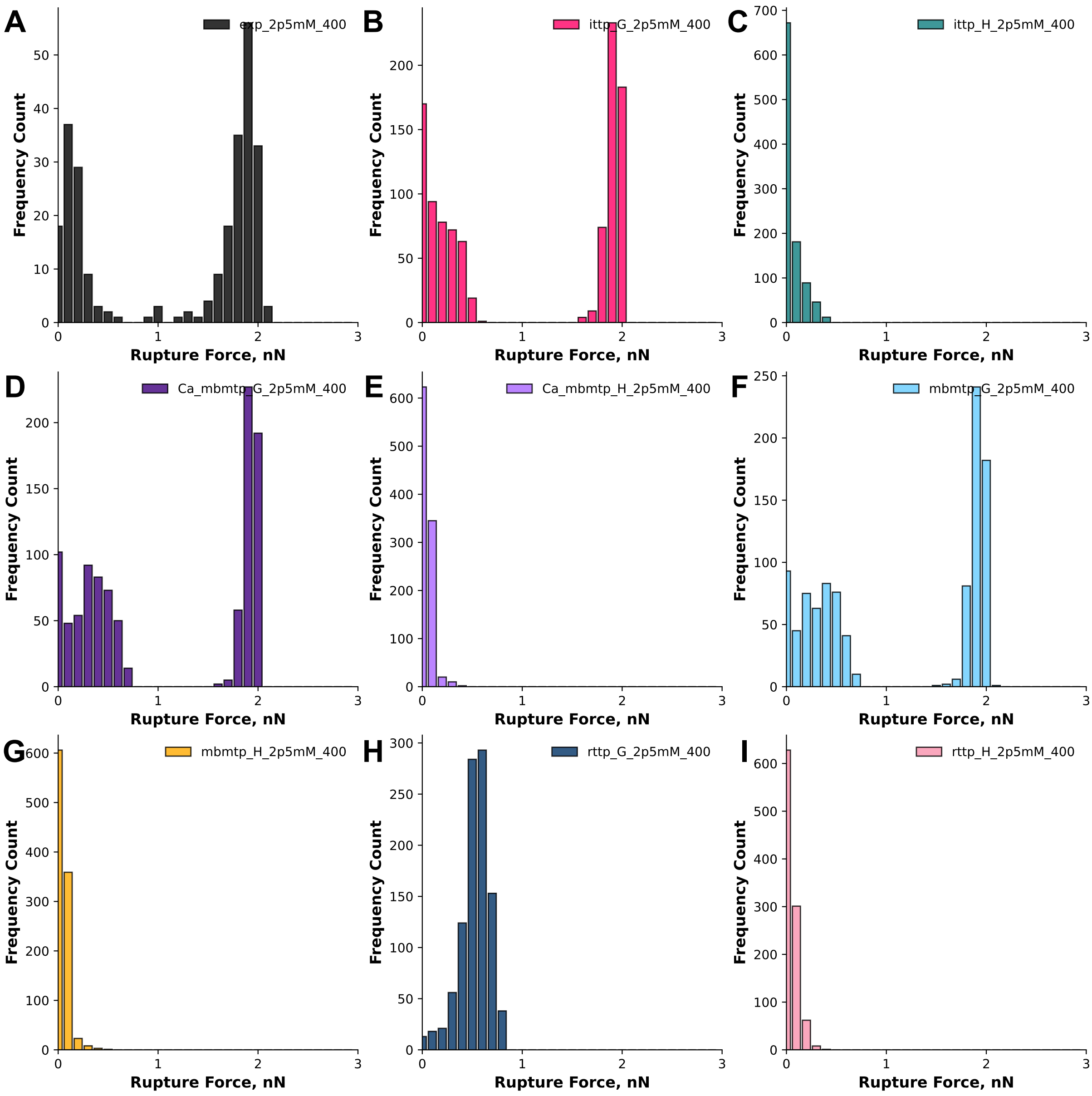


**Figure S3.5:** Histograms of rupture force (nN) vs frequency count for the raw data (black, **A**) obtained from the pulling the N-SdrG-coated cantilever from an Fgβ-coated substrate in TBS with 2.5 mM $Ca^{2+}$ at 400 nm/s and each of the Monte Carlo models, ittp-global (dark pink, **B**), ittp-linear (green, **C**), Ca-mbmtp-global (dark purple, **D**), Ca-mbmtp-linear (purple, **E**), mbmtp-global (blue, **F**), mbmtp-linear (yellow, **G**), rttp-global (dark blue, **H**), rttp-linear (pink, **I**).

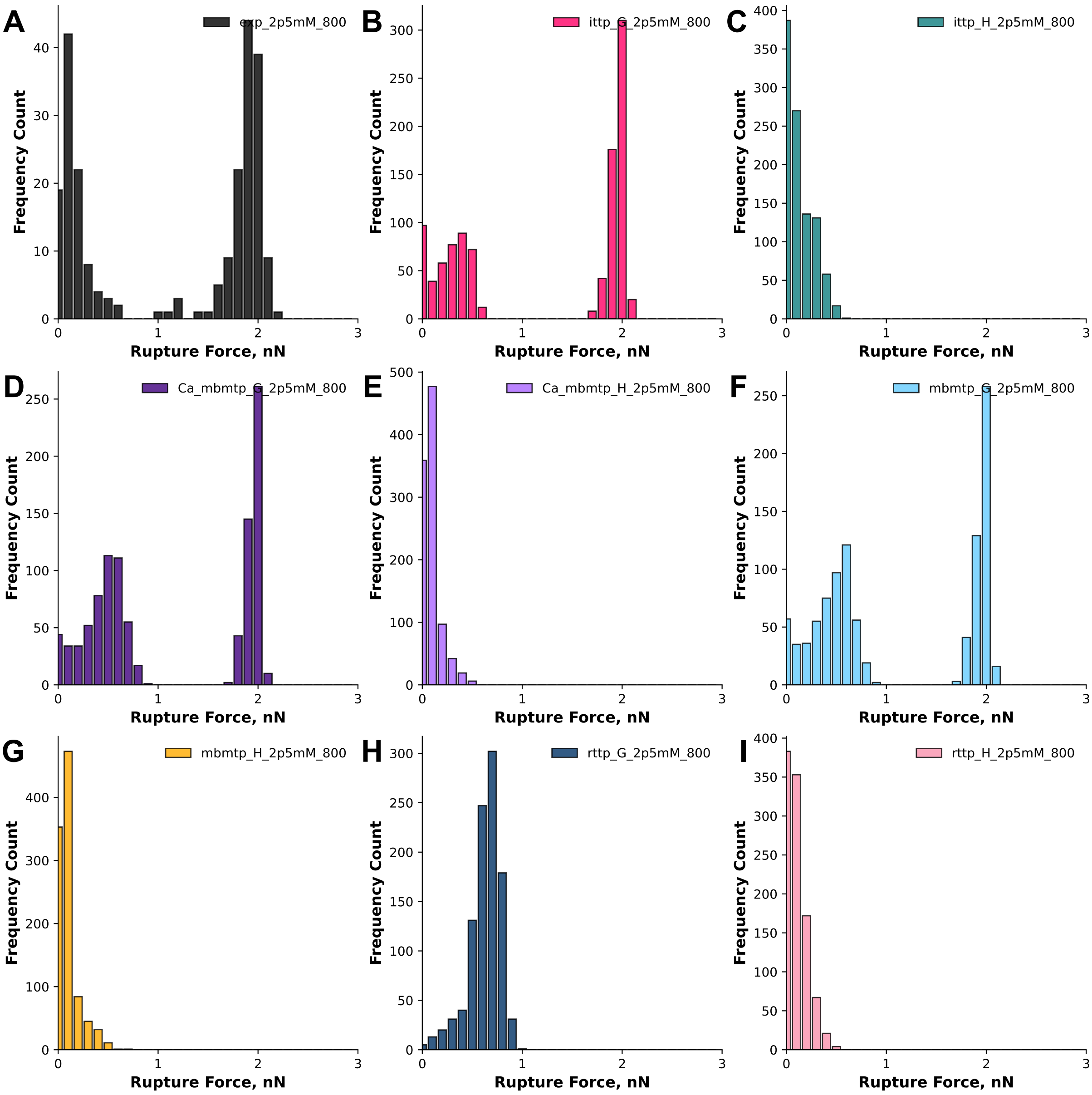


**Figure S3.6:** Histograms of rupture force (nN) vs frequency count for the raw data (black, **A**) obtained from the pulling the N-SdrG-coated cantilever from an Fgβ-coated substrate in TBS with 2.5 mM $Ca^{2+}$ at 800 nm/s and each of the Monte Carlo models, ittp-global (dark pink, **B**), ittp-linear (green, **C**), Ca-mbmtp-global (dark purple, **D**), Ca-mbmtp-linear (purple, **E**), mbmtp-global (blue, **F**), mbmtp-linear (yellow, **G**), rttp-global (dark blue, **H**), rttp-linear (pink, **I**).

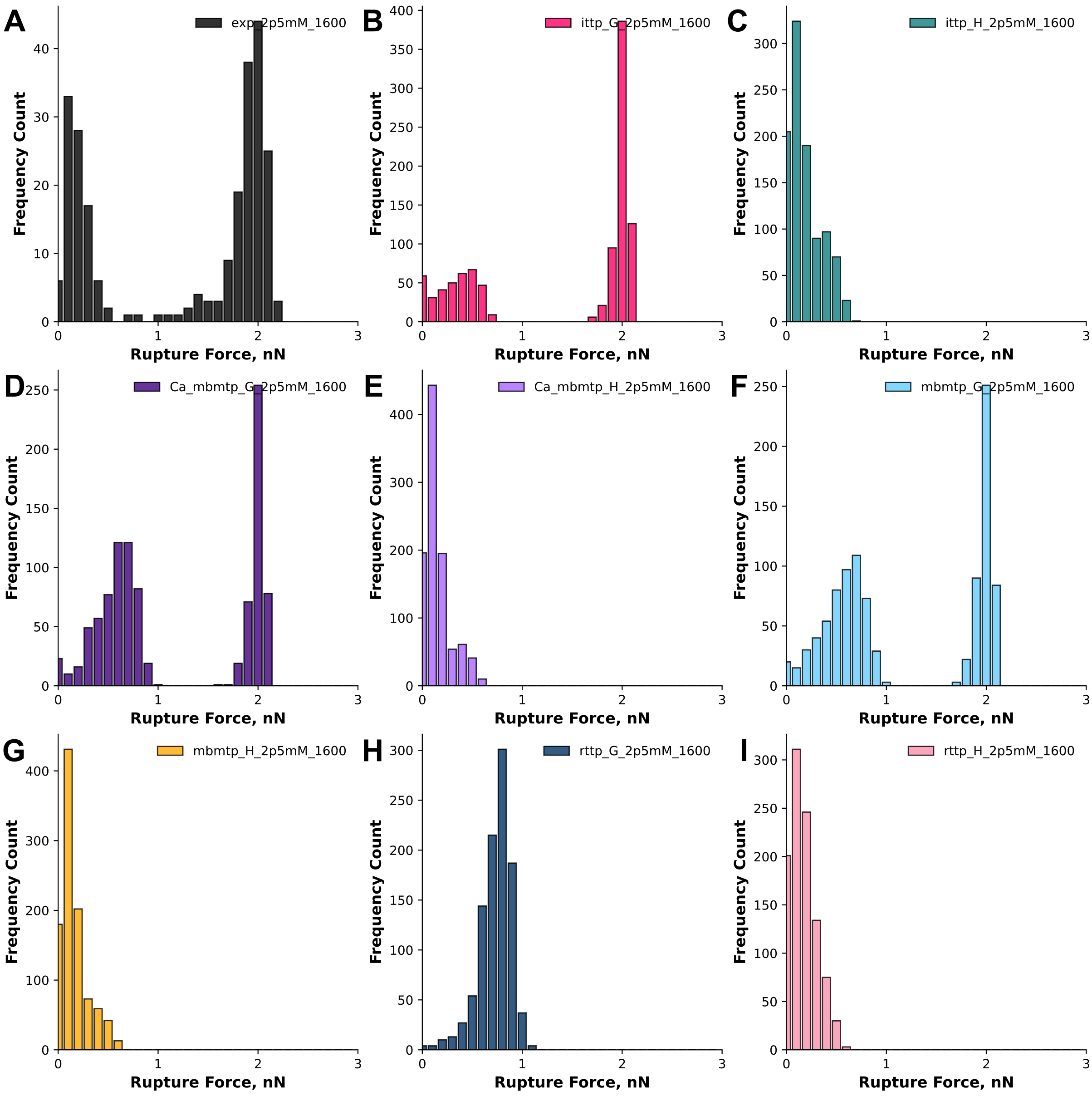


**Figure S3.7:** Histograms of rupture force (nN) vs frequency count for the raw data (black, **A**) obtained from the pulling the N-SdrG-coated cantilever from an Fgβ-coated substrate in TBS with 2.5 mM $Ca^{2+}$ at 1600 nm/s and each of the Monte Carlo models, ittp-global (dark pink, **B**), ittp-linear (green, **C**), Ca-mbmtp-global (dark purple, **D**), Ca-mbmtp-linear (purple, **E**), mbmtp-global (blue, **F**), mbmtp-linear (yellow, **G**), rttp-global (dark blue, **H**), rttp-linear (pink, **I**).

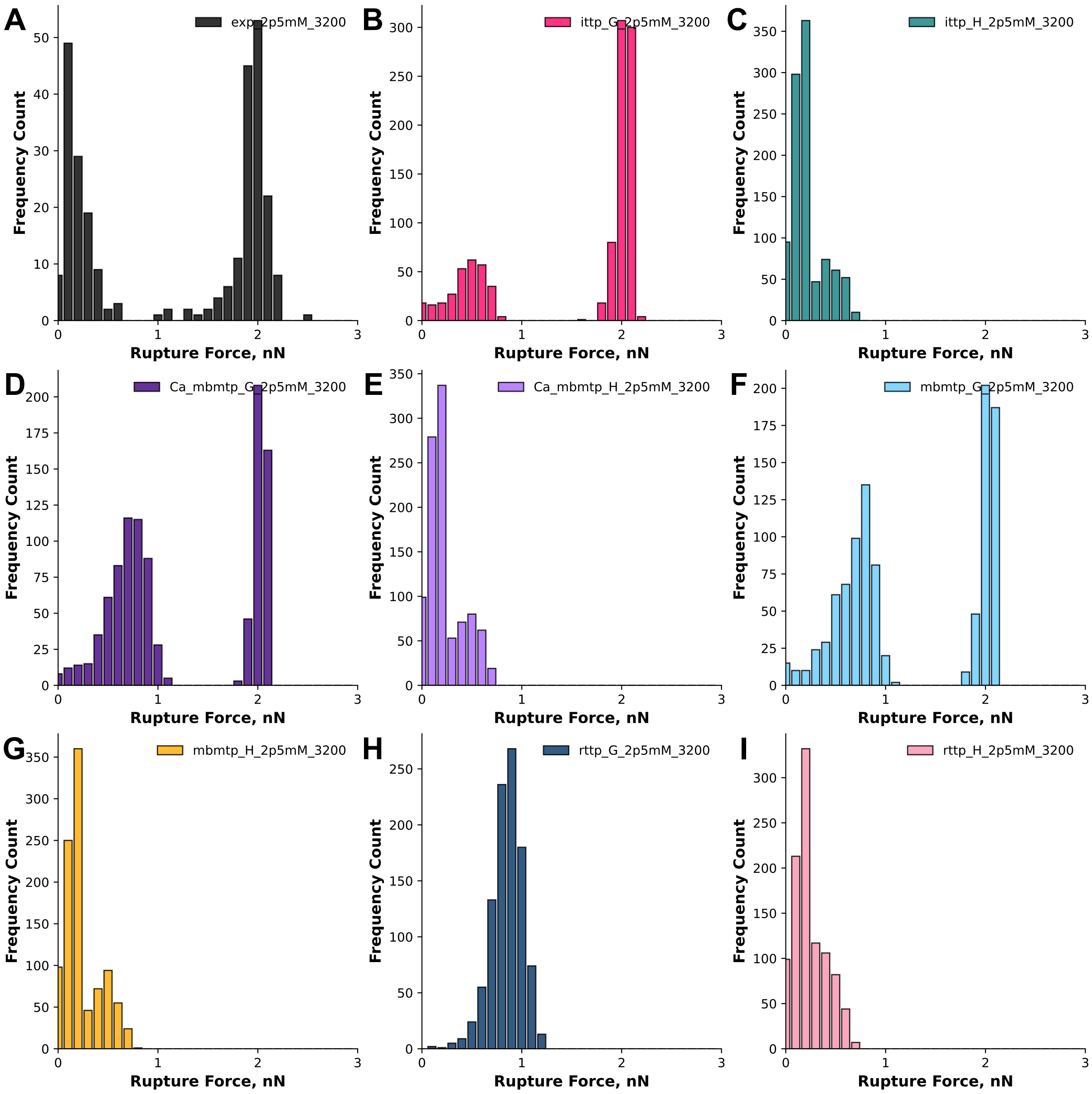


**Figure S3.8:** Histograms of rupture force (nN) vs frequency count for the raw data (black, **A**) obtained from the pulling the N-SdrG-coated cantilever from an Fgβ-coated substrate in TBS with 2.5 mM $Ca^{2+}$ at 3200 nm/s and each of the Monte Carlo models, ittp-global (dark pink, **B**), ittp-linear (green, **C**), Ca-mbmtp-global (dark purple, **D**), Ca-mbmtp-linear (purple, **E**), mbmtp-global (blue, **F**), mbmtp-linear (yellow, **G**), rttp-global (dark blue, **H**), rttp-linear (pink, **I**).

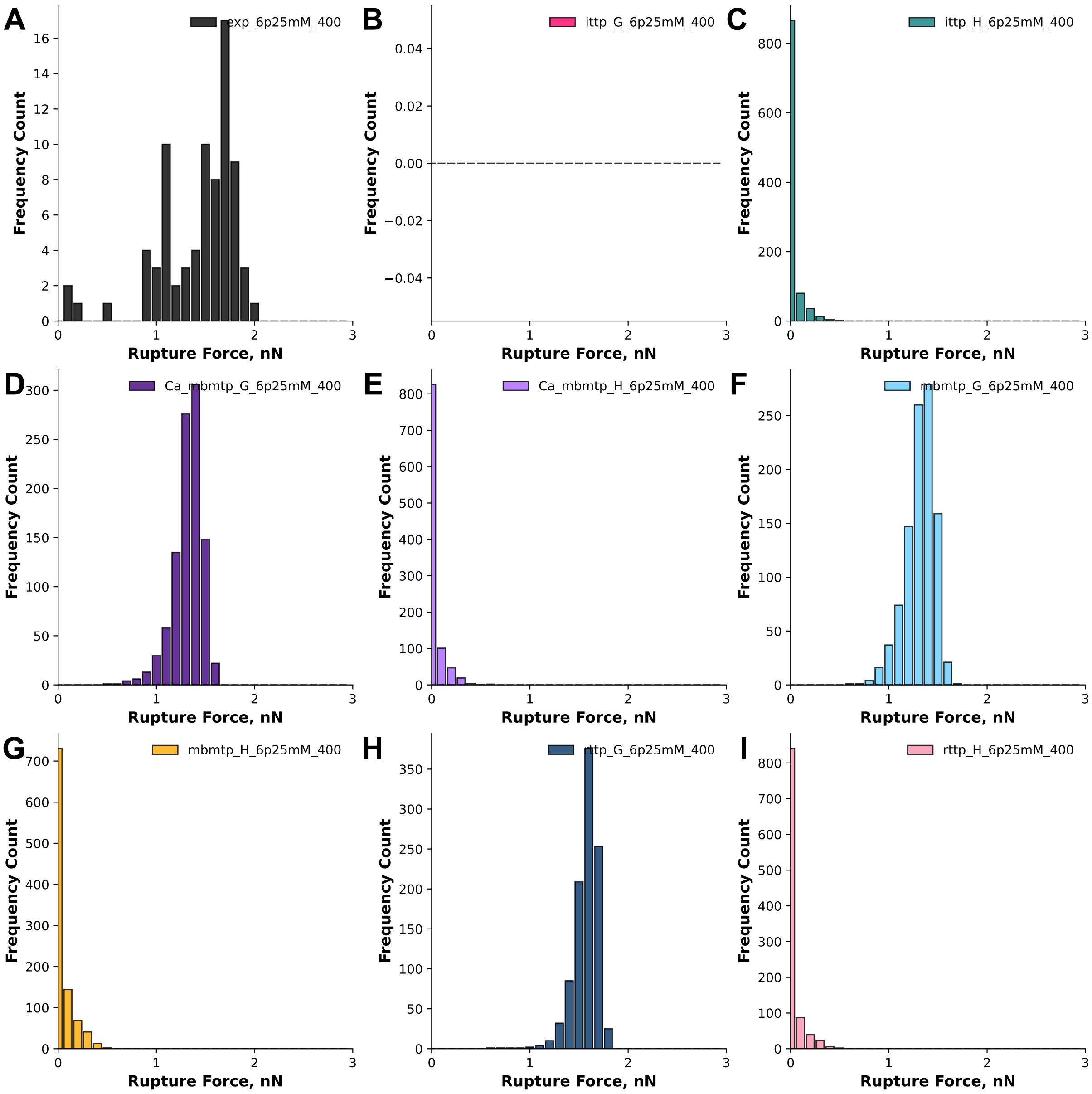


**Figure S3.9:** Histograms of rupture force (nN) vs frequency count for the raw data (black, **A**) obtained from the pulling the N-SdrG-coated cantilever from an Fgβ-coated substrate in TBS with 6.25 mM $Ca^{2+}$ at 400 nm/s and each of the Monte Carlo models, ittp-global (dark pink, **B**), ittp-linear (green, **C**), Ca-mbmtp-global (dark purple, **D**), Ca-mbmtp-linear (purple, **E**), mbmtp-global (blue, **F**), mbmtp-linear (yellow, **G**), rttp-global (dark blue, **H**), rttp-linear (pink, **I**).

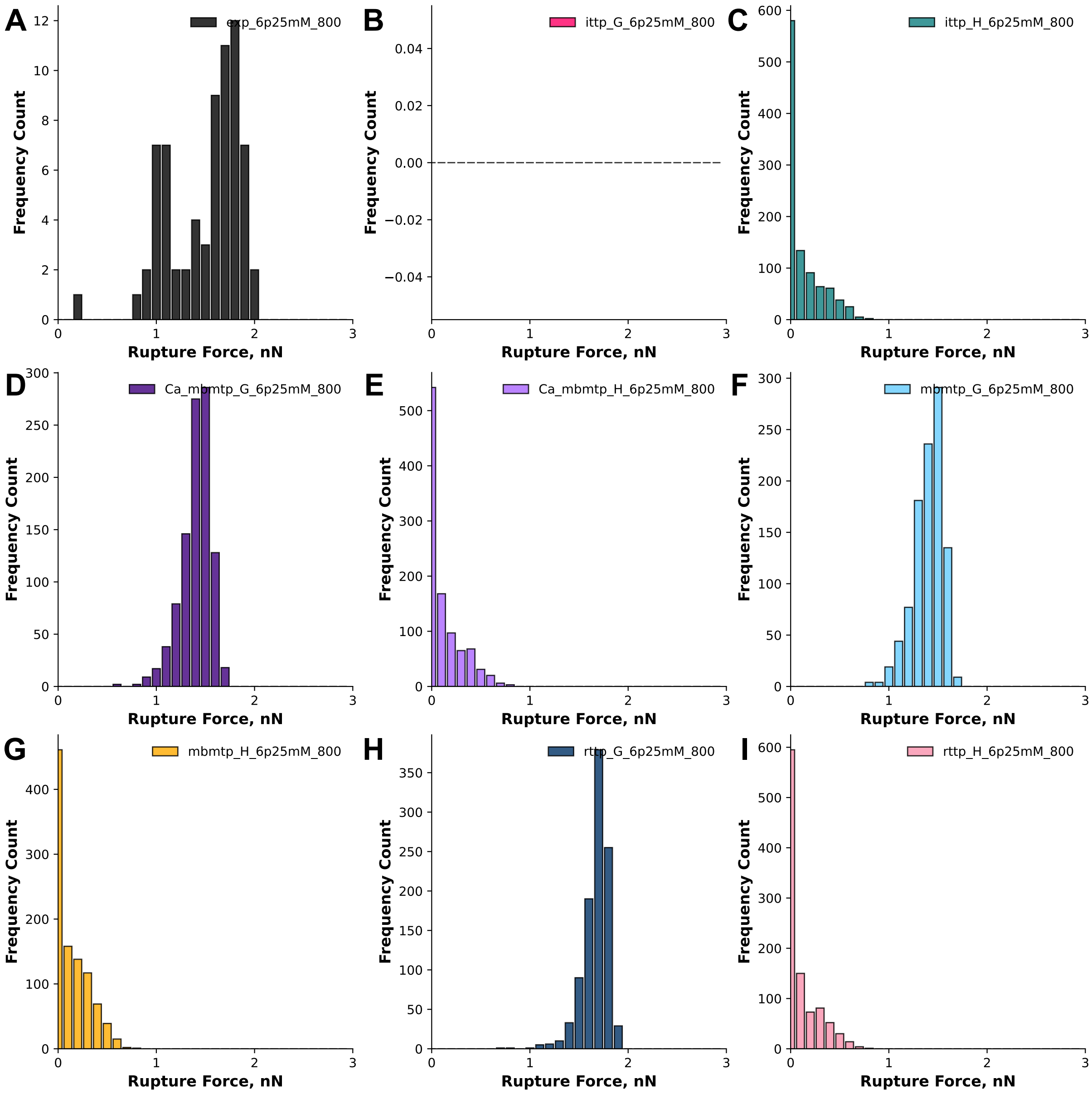


**Figure S3.10:** Histograms of rupture force (nN) vs frequency count for the raw data (black, **A**) obtained from the pulling the N-SdrG-coated cantilever from an Fgβ-coated substrate in TBS with 6.25 mM $Ca^{2+}$ at 800 nm/s and each of the Monte Carlo models, ittp-global (dark pink, **B**), ittp-linear (green, **C**), Ca-mbmtp-global (dark purple, **D**), Ca-mbmtp-linear (purple, **E**), mbmtp-global (blue, **F**), mbmtp-linear (yellow, **G**), rttp-global (dark blue, **H**), rttp-linear (pink, **I**).

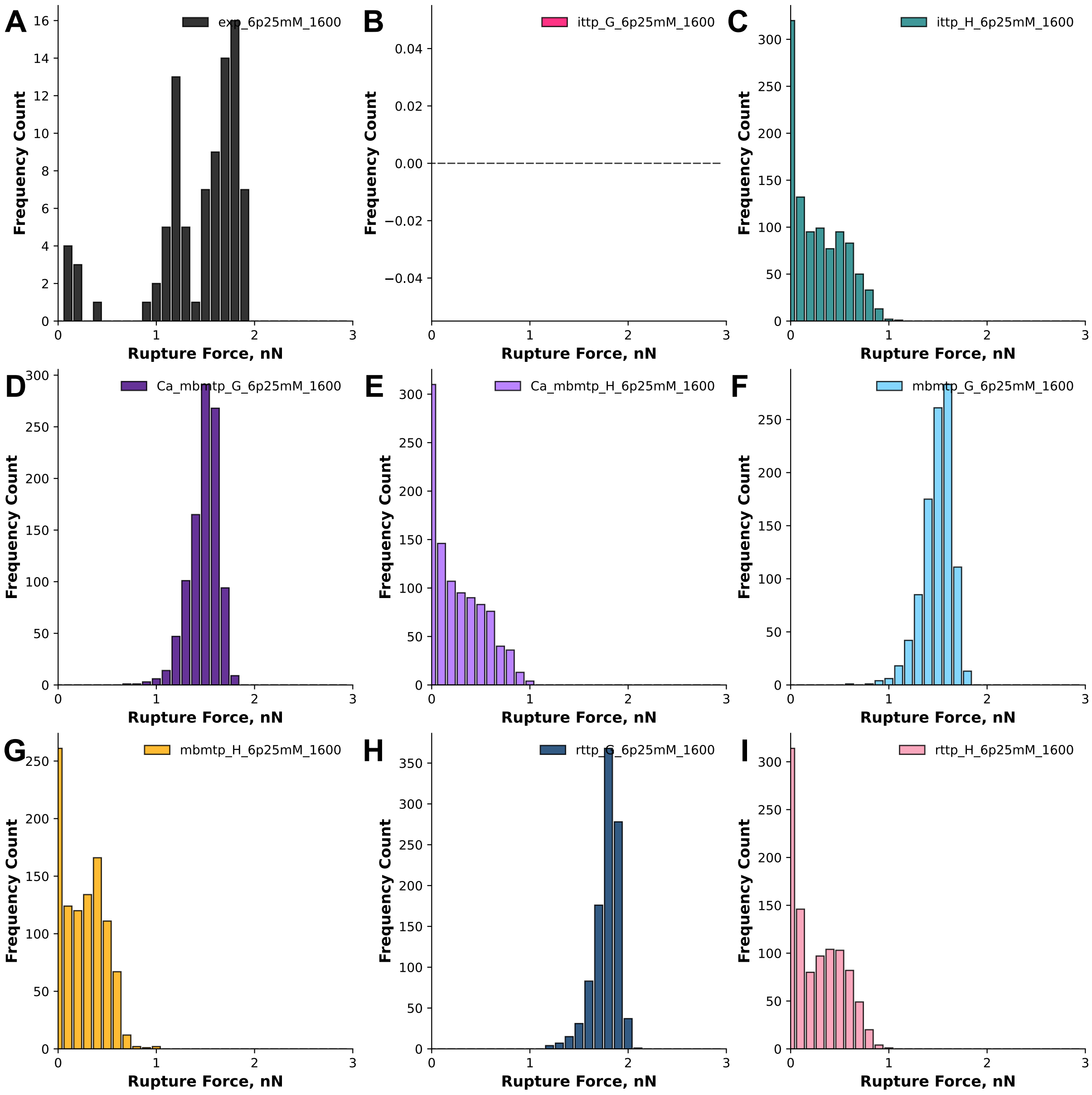


**Figure S3.11:** Histograms of rupture force (nN) vs frequency count for the raw data (black, **A**) obtained from the pulling the N-SdrG-coated cantilever from an Fgβ-coated substrate in TBS with 6.25 mM $Ca^{2+}$ at 1600 nm/s and each of the Monte Carlo models, ittp-global (dark pink, **B**), ittp-linear (green, **C**), Ca-mbmtp-global (dark purple, **D**), Ca-mbmtp-linear (purple, **E**), mbmtp-global (blue, **F**), mbmtp-linear (yellow, **G**), rttp-global (dark blue, **H**), rttp-linear (pink, **I**).

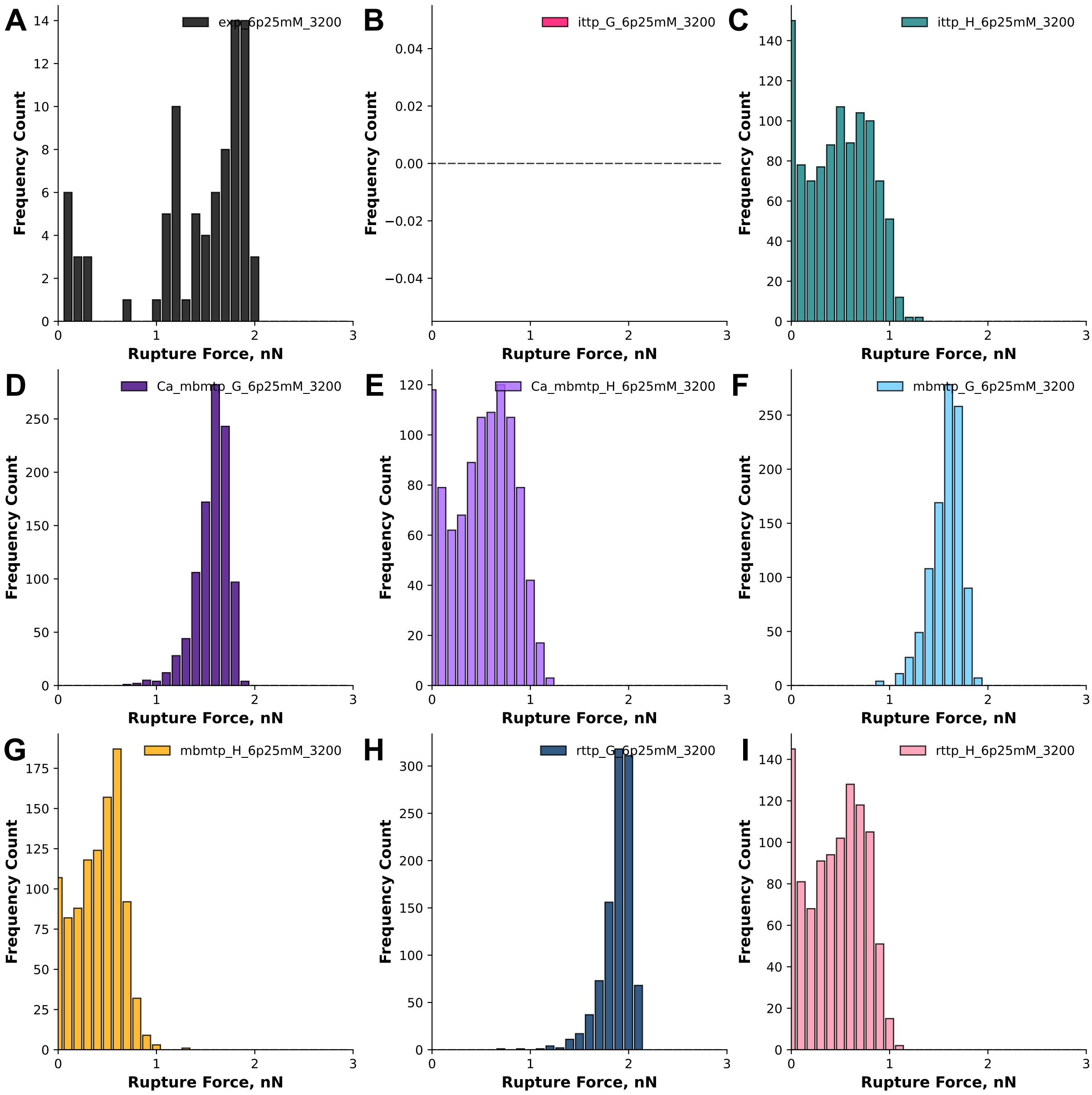


**Figure S3.12:** Histograms of rupture force (nN) vs frequency count for the raw data (black, **A**) obtained from the pulling the N-SdrG-coated cantilever from an Fgβ-coated substrate in TBS with 6.25 mM $Ca^{2+}$ at 3200 nm/s and each of the Monte Carlo models, ittp-global (dark pink, **B**), ittp-linear (green, **C**), Ca-mbmtp-global (dark purple, **D**), Ca-mbmtp-linear (purple, **E**), mbmtp-global (blue, **F**), mbmtp-linear (yellow, **G**), rttp-global (dark blue, **H**), rttp-linear (pink, **I**).

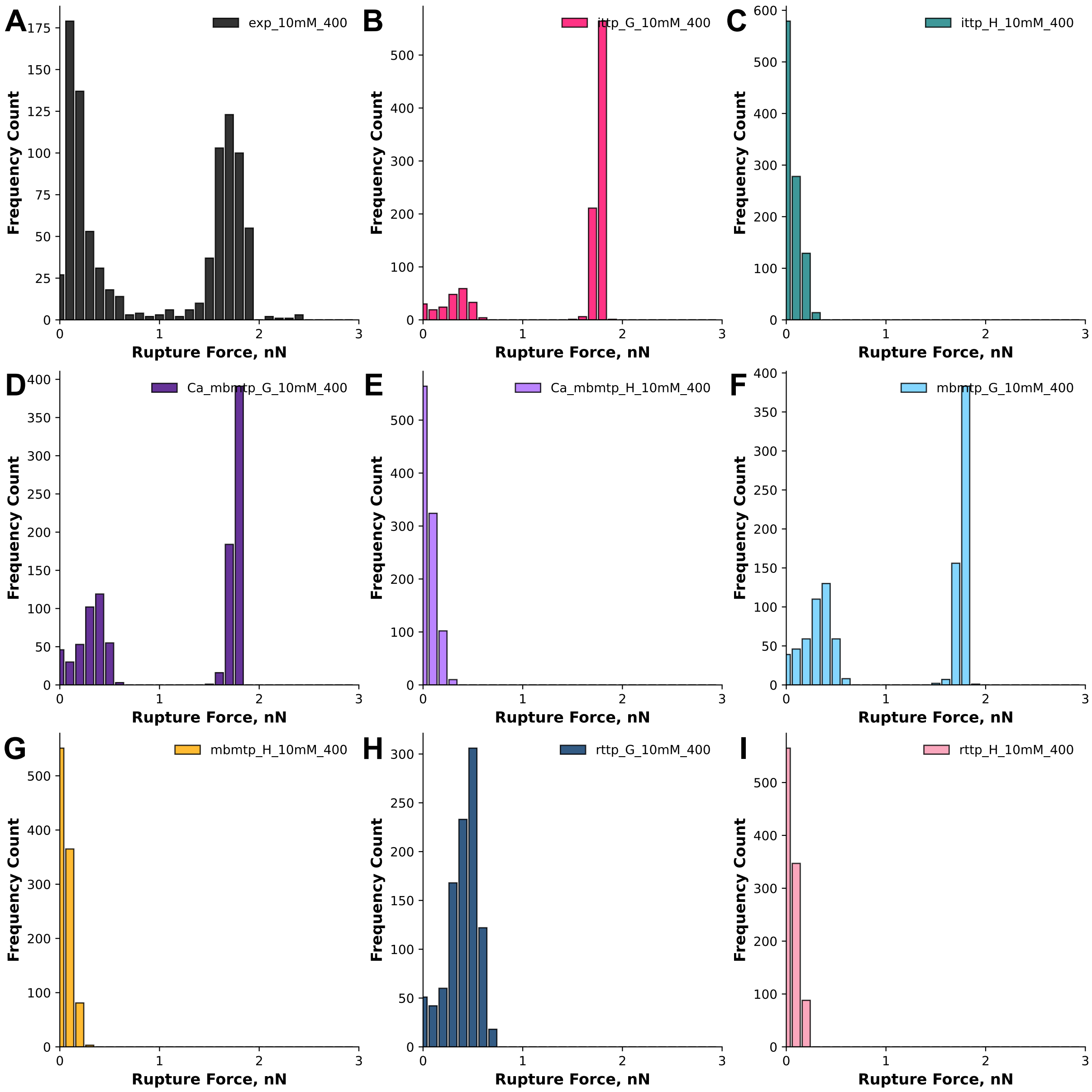


**Figure S3.13:** Histograms of rupture force (nN) vs frequency count for the raw data (black, **A**) obtained from the pulling the N-SdrG-coated cantilever from an Fgβ-coated substrate in TBS with 10 mM $Ca^{2+}$ at 400 nm/s and each of the Monte Carlo models, ittp-global (dark pink, **B**), ittp-linear (green, **C**), Ca-mbmtp-global (dark purple, **D**), Ca-mbmtp-linear (purple, **E**), mbmtp-global (blue, **F**), mbmtp-linear (yellow, **G**), rttp-global (dark blue, **H**), rttp-linear (pink, **I**).

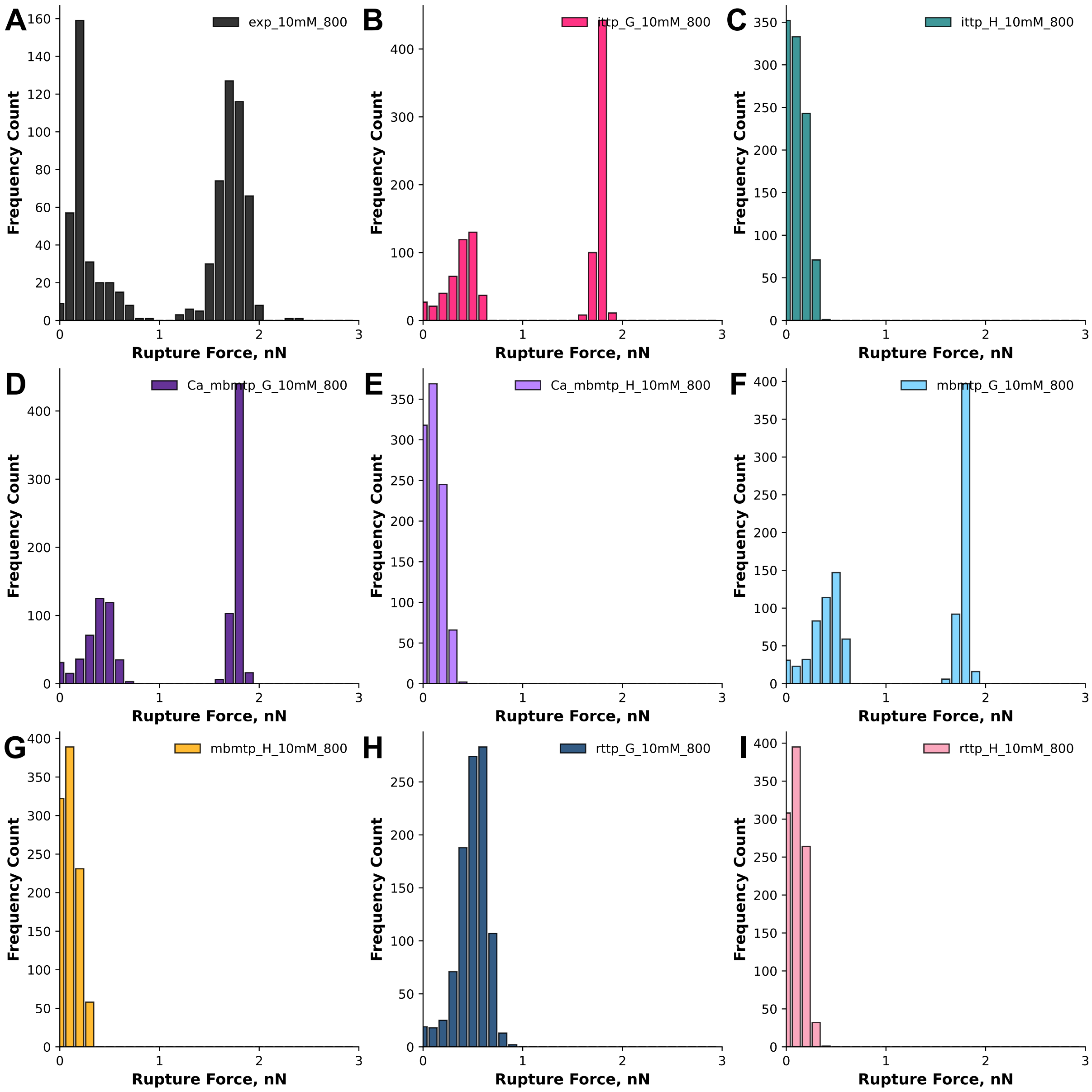


**Figure S3.14:** Histograms of rupture force (nN) vs frequency count for the raw data (black, **A**) obtained from the pulling the N-SdrG-coated cantilever from an Fgβ-coated substrate in TBS with 10 mM $Ca^{2+}$ at 800 nm/s and each of the Monte Carlo models, ittp-global (dark pink, **B**), ittp-linear (green, **C**), Ca-mbmtp-global (dark purple, **D**), Ca-mbmtp-linear (purple, **E**), mbmtp-global (blue, **F**), mbmtp-linear (yellow, **G**), rttp-global (dark blue, **H**), rttp-linear (pink, **I**).

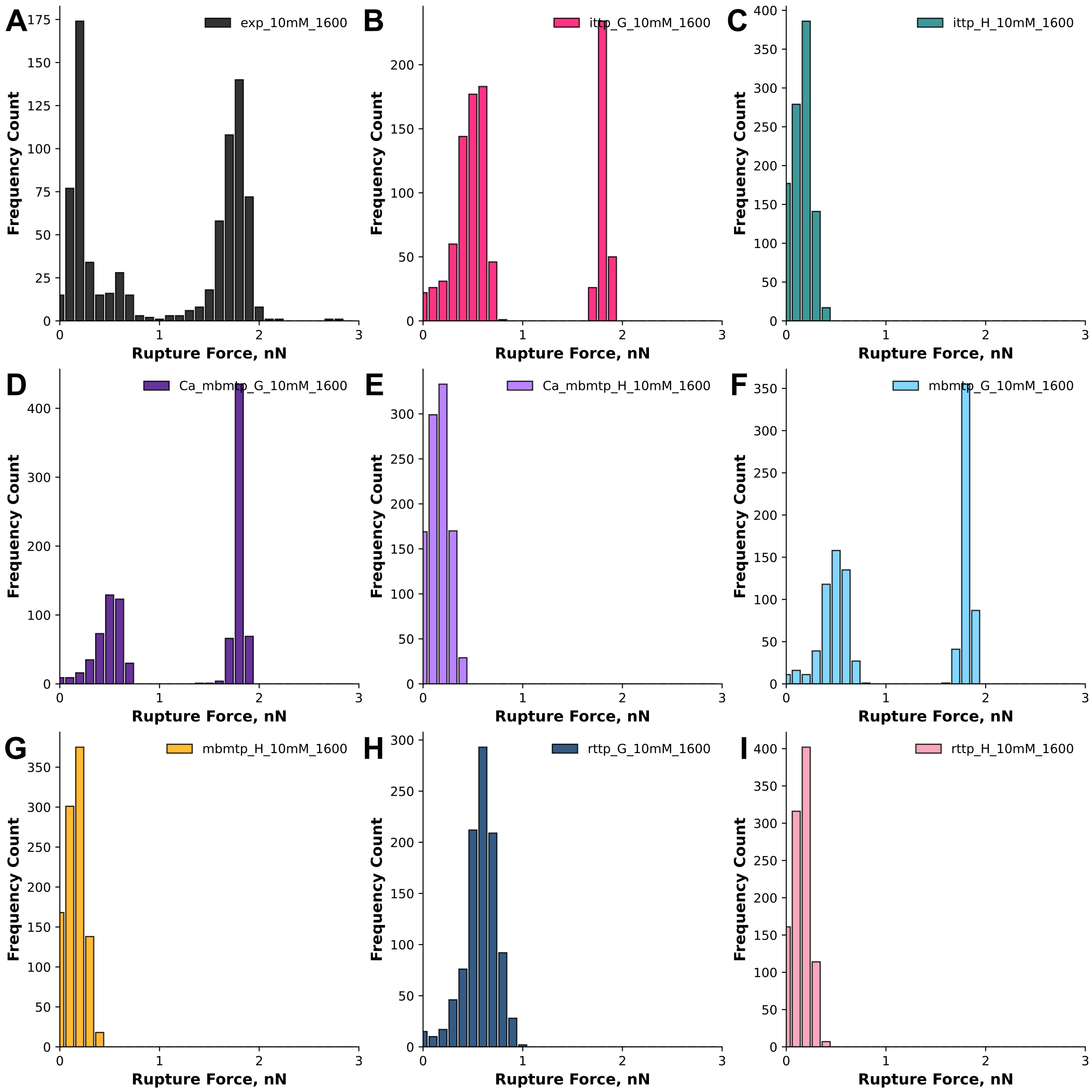


**Figure S3.15:** Histograms of rupture force (nN) vs frequency count for the raw data (black, **A**) obtained from the pulling the N-SdrG-coated cantilever from an Fgβ-coated substrate in TBS with 10 mM $Ca^{2+}$ at 1600 nm/s and each of the Monte Carlo models, ittp-global (dark pink, **B**), ittp-linear (green, **C**), Ca-mbmtp-global (dark purple, **D**), Ca-mbmtp-linear (purple, **E**), mbmtp-global (blue, **F**), mbmtp-linear (yellow, **G**), rttp-global (dark blue, **H**), rttp-linear (pink, **I**).

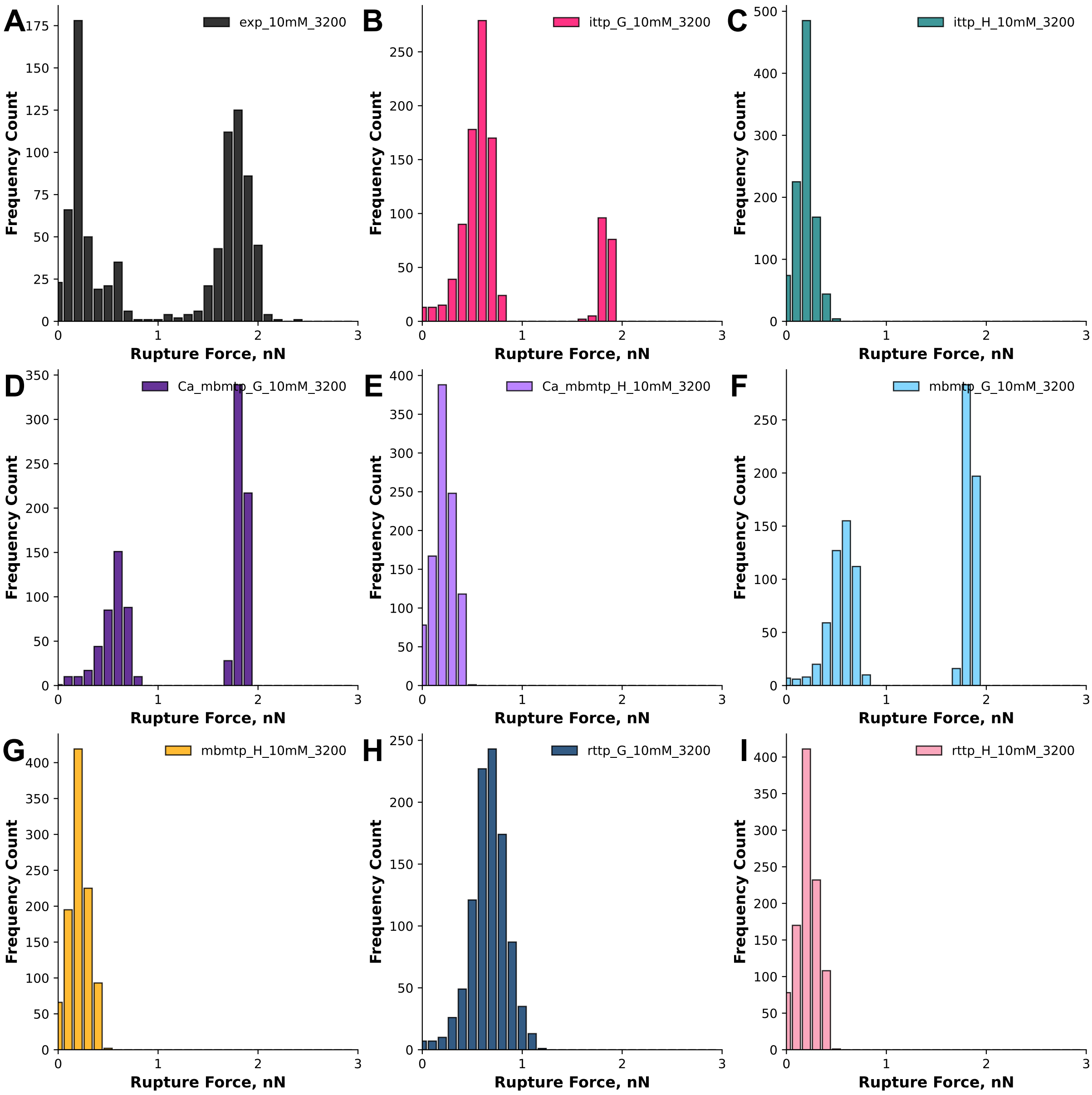


**Figure S3.16:** Histograms of rupture force (nN) vs frequency count for the raw data (black, **A**) obtained from the pulling the N-SdrG-coated cantilever from an Fgβ-coated substrate in TBS with 10 mM $Ca^{2+}$ at 3200 nm/s and each of the Monte Carlo models, ittp-global (dark pink, **B**), ittp-linear (green, **C**), Ca-mbmtp-global (dark purple, **D**), Ca-mbmtp-linear (purple, **E**), mbmtp-global (blue, **F**), mbmtp-linear (yellow, **G**), rttp-global (dark blue, **H**), rttp-linear (pink, **I**).

#### *S3.3 Statistical analysis of MC models vs raw data*

A variety of statistical methods were used to determine how similar the MC predictions were to the raw data but challenges existed in getting a reliable comparison due to the high number of different experimental conditions, the different extraction parameters and the different numbers of datapoints in both the predictions and the raw datasets. For this reason we explored the Wasserstein distance to compare datasets, as it is robust to sampling differences and is more sensitive to the geometry of distributions. The comparison of all the datasets is shown in Figure **S3.17.**

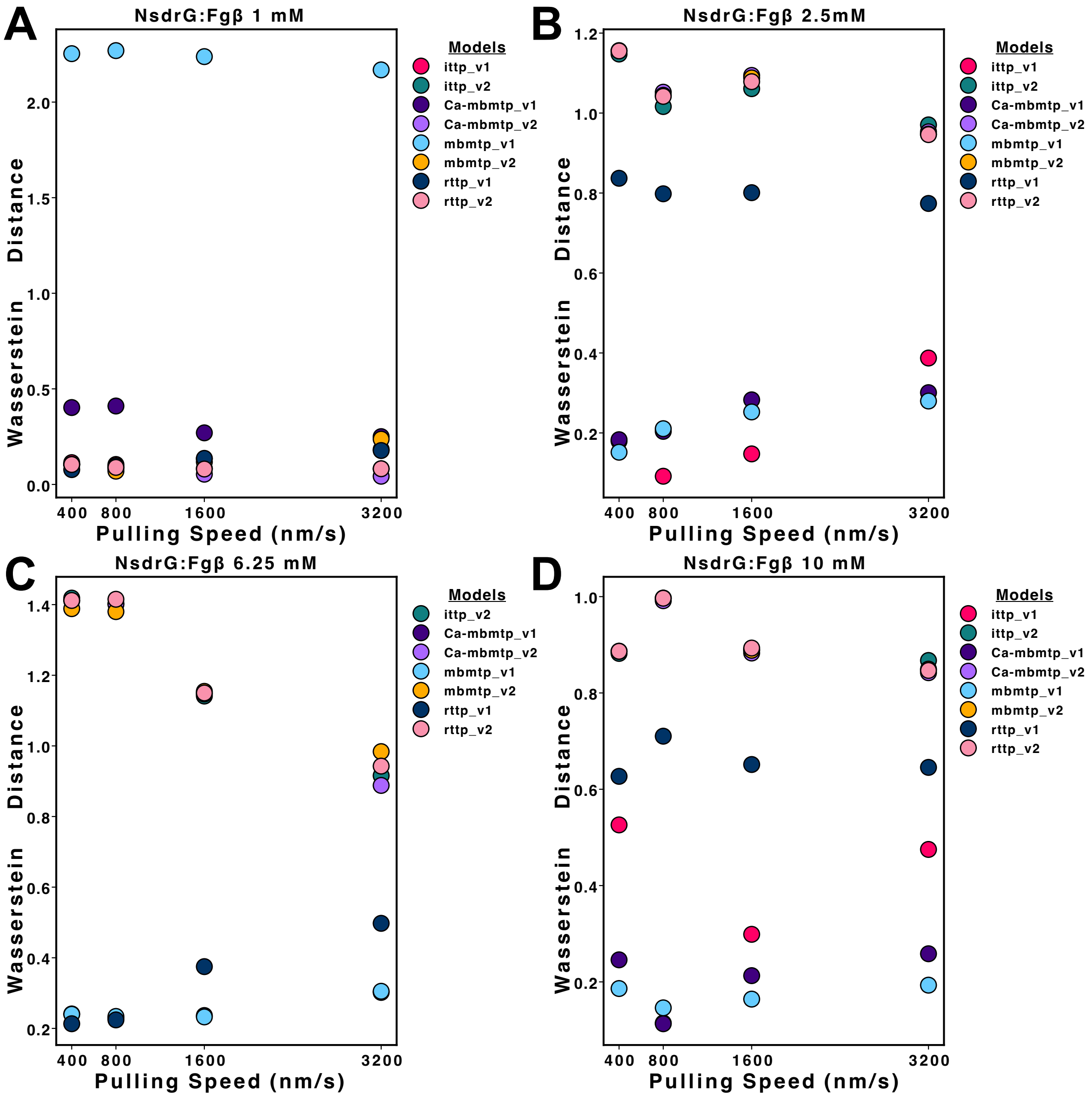


**Figure S3.17:** Wasserstein plots of pulling speed (nm/s) vs Wasserstein Distance for each of the Monte Carlo models for N-SdrG:Fgβ per calcium concentration (**A**) 1 mM, (**B**) 2.5 mM, (**C**) 6.25 mM, and (**D**) 10 mM, where each is labelled in the colour related to Figures **S3.1-S3.16**, ittp-global (dark pink), ittp-linear (green), Ca-mbmtp-global (dark purple), Ca-mbmtp-linear (purple), mbmtp-global (blue), mbmtp-linear (yellow), rttp-global (dark blue), rttp-linear (pink).

#### S4 Computational Fluid Dynamics

Computation fluid dynamics (CFD) is a numerical technique employed to solve fluid flow and heat transfer problems using the finite volume method (FVM) to determine approximate solutions. To investigate cell detachment and quantify the hydrodynamic forces exerted on mesoscale spherical particles (i.e., within the cell population), CFD models of the spinning disk adhesion (SDA) assays were employed, which have been well-studied over the past decade. For this study, a computer-aided design (CAD) model of the disk was generated in ANSYS Design Modeller. Spherical particles with a diameter of 5 μm were distributed along the disk's radial axis at uniform intervals, with one particle positioned at the centre. Additional particles were arranged in four rows aligned with both the positive and negative directions of the horizontal and vertical axes. A cylindrical fluid domain surrounding the disk was then created, after which the solid disk body was subtracted to define the fluid region. The computational setup is illustrated in a previous report, while the design specifications of the spinning disc, particles, and domain are provided in Table **S4.1**.

**Table S4.1.** Geometric parameters of the spinning disk system

| | |
|---|---|
| Disk diameter | 25 mm |
| Disk thickness | 196 μm |
| Particle diameter | 5 μm |
| Distance between particle and disc | 0.3 μm |
| Fluid domain diameter | 29 mm |
| Fluid domain depth | 21 mm |
| Particle spacing | 0.735 mm |

##### Meshing and CFD Setup

After the fluid domain was defined, it was discretised into control volumes for solving the Reynolds-Averaged Navier–Stokes (RANS) equations using the finite volume method. A tetrahedral mesh was applied, with local refinement near the disc surface and spherical particles. The mesh resolution was set to 2 μm over both the disc and the particles, with additional refinement in regions of curvature and close spacing. Each surface of the computational domain was labelled, and particles were sequentially numbered outward from the centre: the central particle was defined as sphere_0, followed by sphere_1 adjacent to it, and so on along the radial line toward the disk edge.

The CFD simulations were conducted under steady-state conditions with a pressure-based solver. Water was used as the working fluid, with density 998.5 kg/m$^3$ and viscosity 0.001 Pa·s. Flow was assumed to be laminar to reduce computational cost and runtime. The angular velocity of the spinning disc was imposed via a moving reference frame approach. The cell detachment and hydrodynamic forces on cells attached to the spinning disk were analysed at angular velocities of 3000 rpm and 4000 rpm. The magnitude of the liquid velocity over the cells was calculated as shown in Figure **S4.1**.

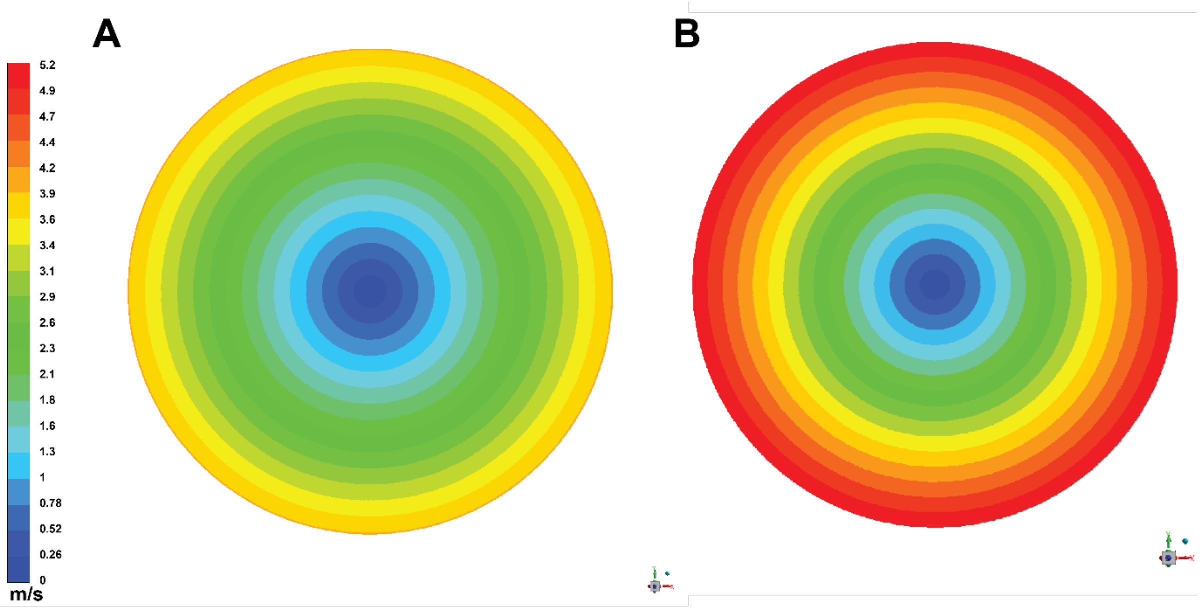


**Figure S4.1:** The magnitude of the liquid velocity over the disk surface in m/s for spinning at 3000 rpm (**A**) and 4000 rpm (**B**) used to calculate the drag and lift forces shown in the main text

#### S5 Yeast Expression and Display

##### *S5.1 Yeast Display*

Before investigation by SDA assays, it was necessary to express each protein construct described previously on yeast cells and optimise their display. N-SdrG, C-SdrG and $N_{mut}$-SdrG were displayed with the ddFLN4, yBBr and His-Tag domains intact, as for the AFM investigations, to avoid the (albeit unlikely) scenario that the absence of these domains would alter the observed behaviour of the SdrG. Maintaining the correct orientation of the SdrG (N- or C-terminus pulling), when expressed on the surface of *Saccharomyces cerevisiae*, necessitated the use of two different expression vectors. A pCHA expression vector was used for the N-SdrG and the $N_{mut}$-SdrG constructs, with both displayed at the C-terminus of the Aga2p domain ensuring their N-termini remained free to interact with Fg or Fgβ. A pYD1 expression vector was used for the C-SdrG construct. Here, C-SdrG is expressed at the N-terminus of the Aga2p domain, maintaining the free C-terminus orientation. Plasmid maps for the three constructs are provided in Figures **S5.1**, **S5.2** and **S5.3**.

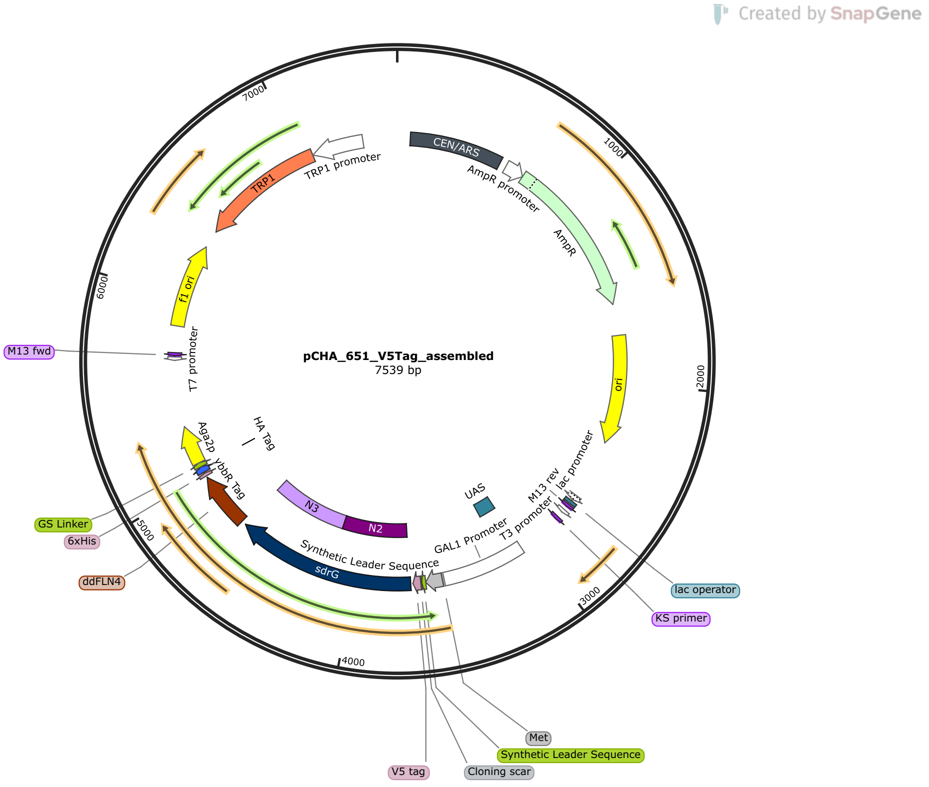


**Figure S5.1:** Plasmid map for the N-SdrG yeast display construct with ddFLN4, His-tag and yBBR domains in a pCHA yeast expression vector created in SnapGene

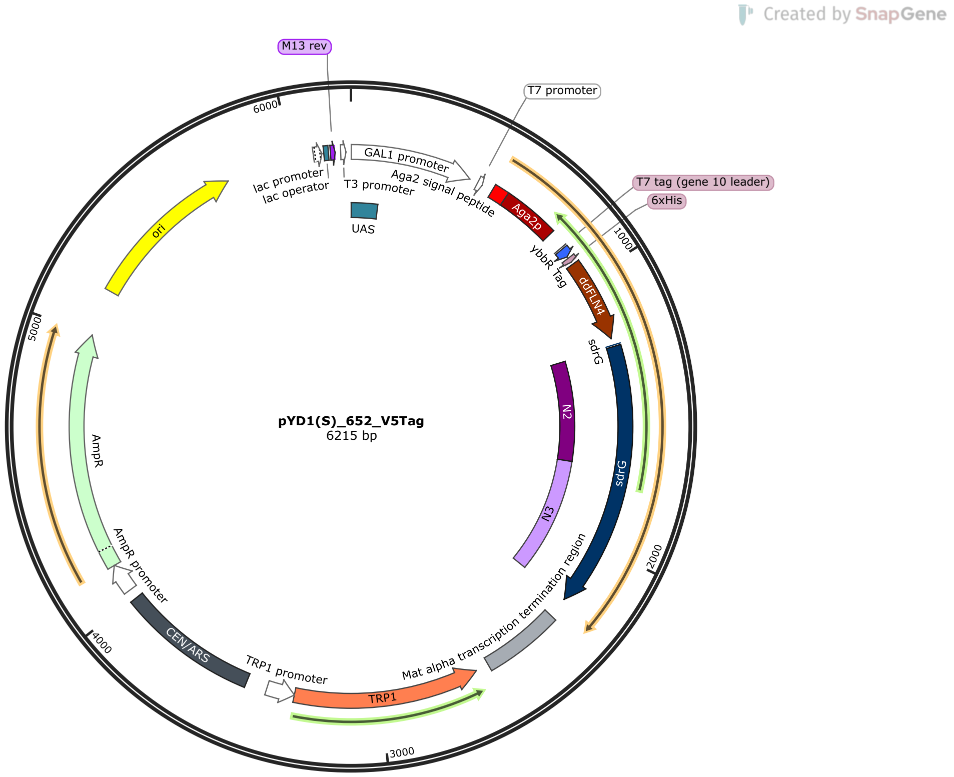


**Figure S5.2:** Plasmid map for the C-SdrG yeast display construct with ddFLN4, His-tag and yBBR domains in a pYD1 yeast expression vector created in SnapGene

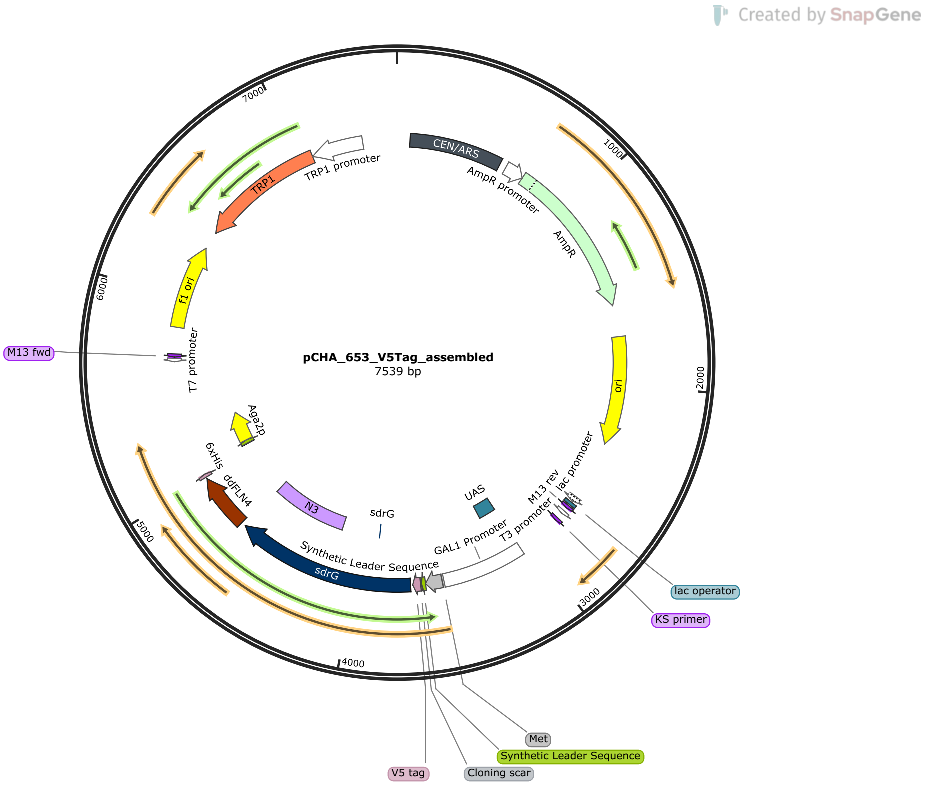


**Figure S5.3:** Plasmid map for the N$_{mut}$-SdrG yeast display construct with ddFLN4, His-tag and yBBR domains in a pCHA yeast expression vector created in SnapGene

To ensure optimal expression, the culture medium was buffered with citrate-phosphate buffer pH 3, pH 5 or pH 7 or cultured without buffer (NB). The degree of expression was determined using flow cytometry, before which cells were incubated with a primary mouse antibody specific for the His-tag, then stained with a goat anti-mouse secondary antibody conjugated to Alexa Fluor 594. Plotting the readout from the blue filter (BL1-H) against the yellow filter (YL2-H) allows us to quantify expression of each construct as a function of buffer pH. The flow cytometry plot showing display as a function of buffering is given in Figure **S5.4**.

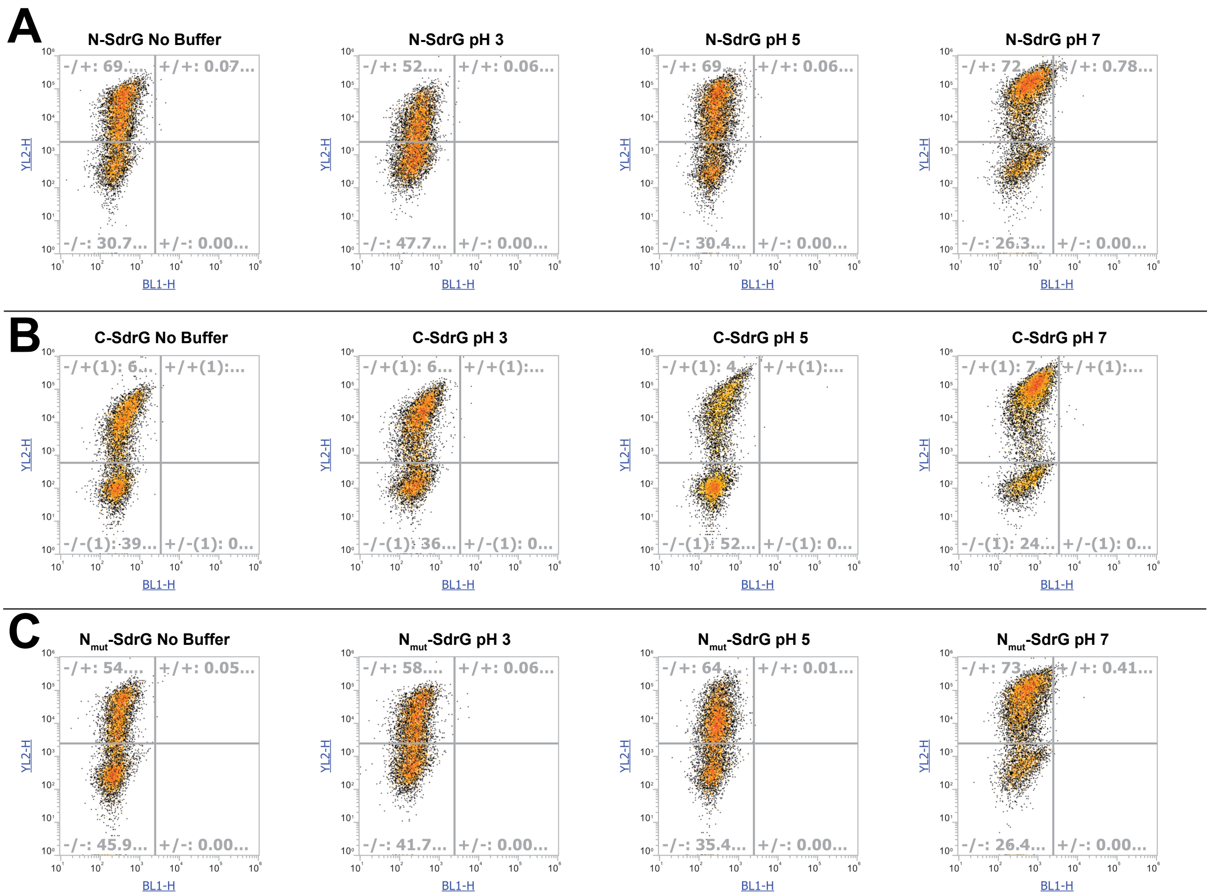


**Figure S5.4:** Plots of absolute signal intensity from the blue channel (BL1-H) vs absolute signal intensity from the yellow channel (YL2-H) are shown for (**A**) N-SdrG, (**B**) C-SdrG and (**C**) $N_{mut}$-SdrG, at each buffer concentration, with NB being the non-buffered cultures.

The median fluorescence for multiple display rounds (n = 5) are shown in Figure S5.3. This was also determined from the flow cytometry data and performed to ensure that similar absolute expression levels were obtained, or at least that the N-SdrG was not more expressed than other constructs. It was observed that the pYD1 vector expressed at twice the rate of the pCHA vectors (Figure **S5.5**), however, this did not appear to have a significant impact on the outcome of the SDA assay results described in the following section, in which the C-SdrG performed significantly worse than the N-SdrG construct.

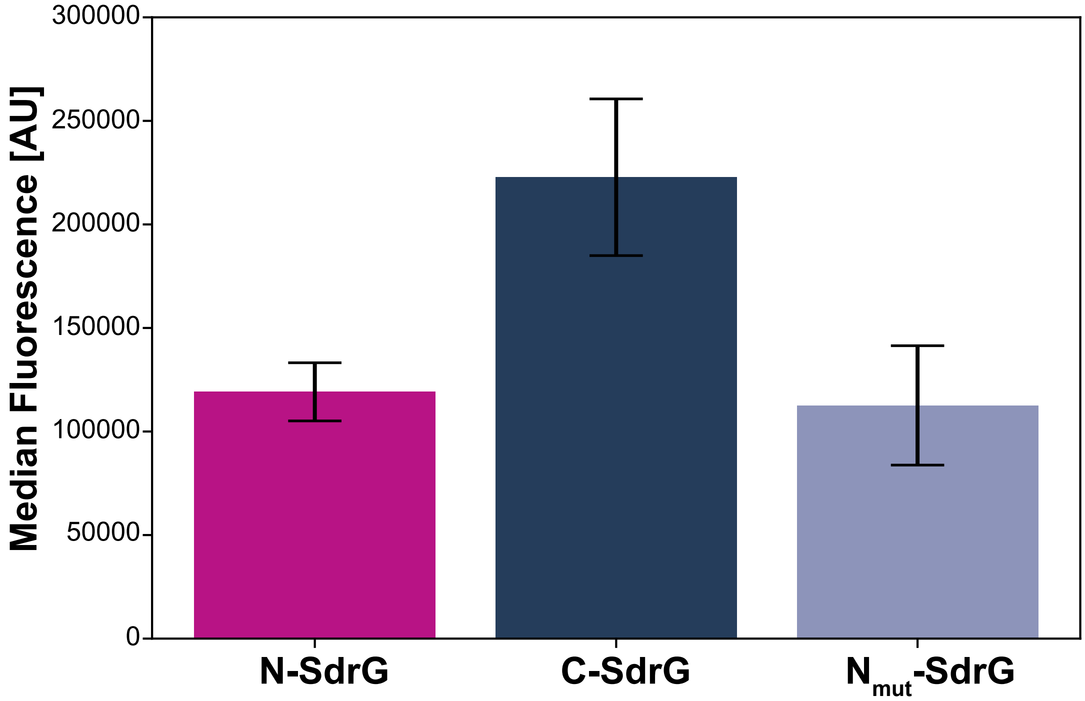


**Figure S5.5:** Median fluorescence intensity averaged from three samples for each construct; all measurements are from the optimal pH 7 buffering condition.

### S6 Spinning Disk Extended Data

#### *S6.1 Cumulative density plots*

For the spinning disk, each experiment was run between 6 and 13 times, with each dataset an average of 5 measurements representative of the cell distribution across the surface. To verify catch bond behaviour, cumulative density plots (Figure **S6.1** and **S6.2**) were prepared from the extracted cell coordinates from each disk under each condition. Catch bond behaviour is identified by the lack of decrease in cell distribution as a function of distance from the disk centre.

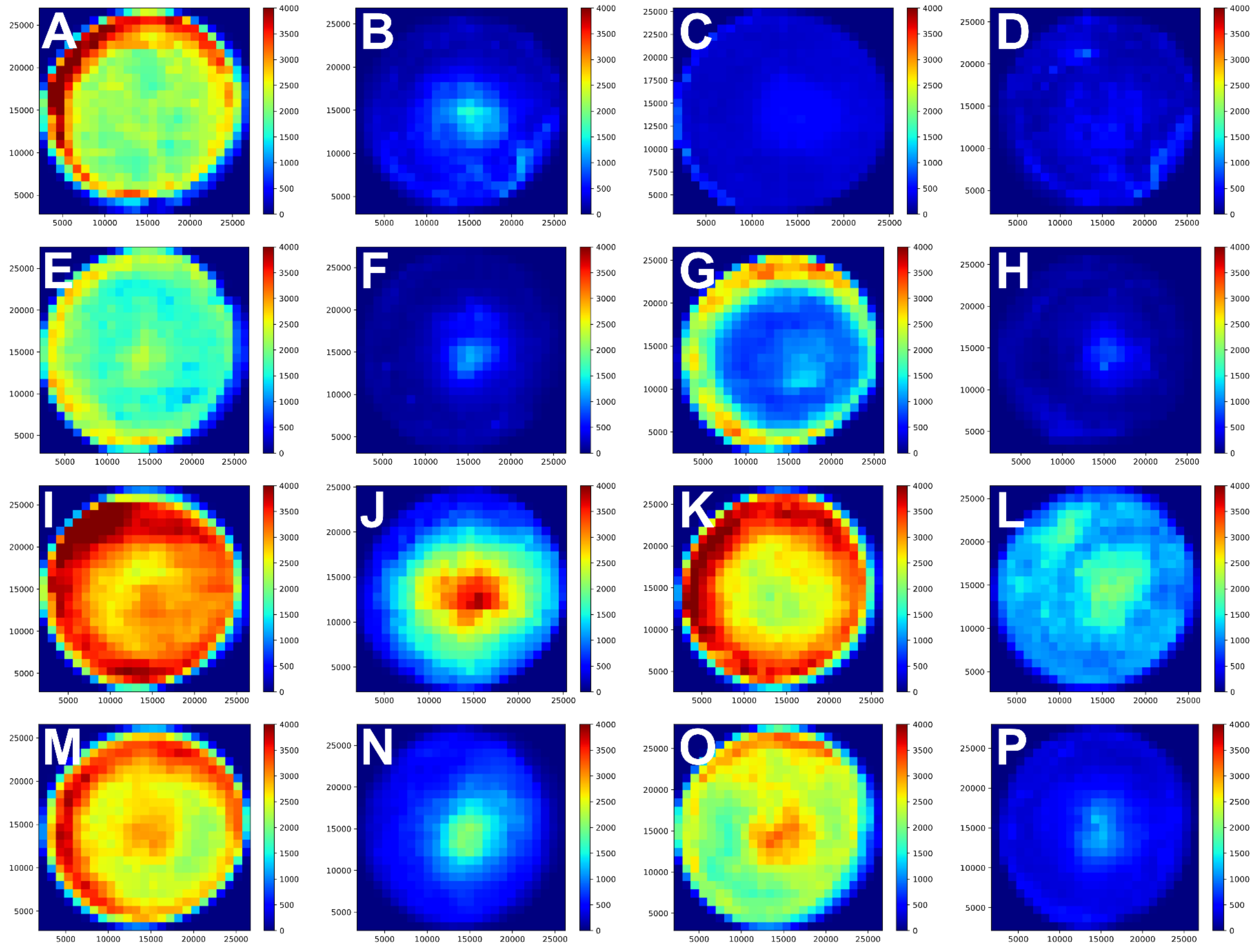


**Figure S6.1:** Cumulative density plot of cell distribution across disk surface for (**A**) N-SdrG, (**B**) C-SdrG against Fg substrate and (**C**) N-SdrG, (**D**) C-SdrG against Fgβ substrate, at 3000 rpm in TBS with 1 mM $Ca^{2+}$, (**E**) N-SdrG, (**F**) C-SdrG against Fg substrate and (**G**) N-SdrG, (**H**) C-SdrG against Fgβ substrate, at 3000 rpm in TBS with 10 mM $Ca^{2+}$, (**I**) N-SdrG, (**J**) C-SdrG against Fg substrate and (**K**) N-SdrG, (**L**) C-SdrG against Fgβ substrate, at 4000 rpm in TBS with 1 mM $Ca^{2+}$, (**M**) N-SdrG, (**N**) C-SdrG against Fg substrate and (**O**) N-SdrG, (**P**) C-SdrG against Fgβ substrate, at 4000 rpm in TBS with 10 mM $Ca^{2+}$.

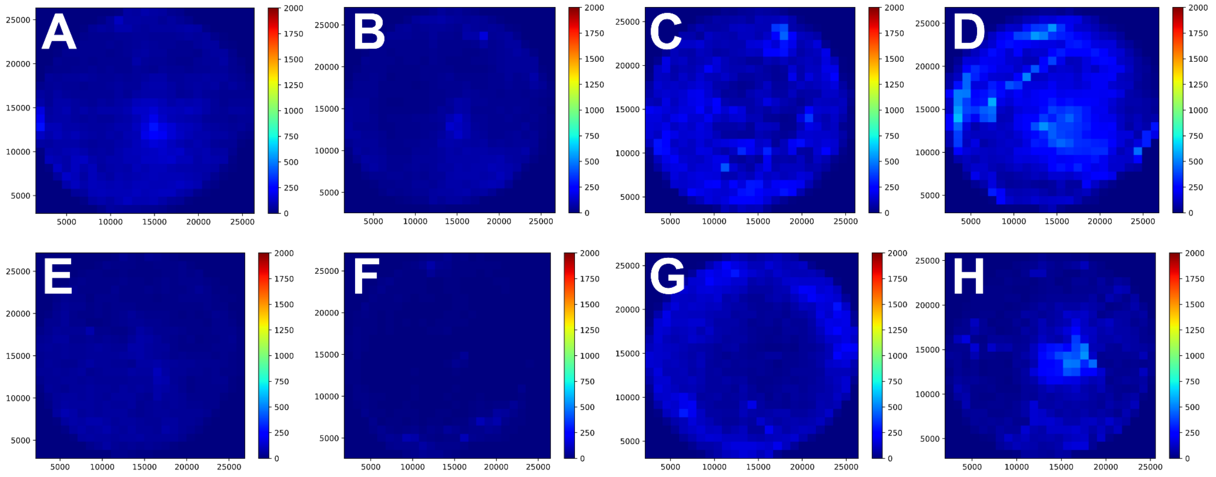


**Figure S6.2:** Cumulative density plot of cell distribution across disk surface for N$_{mut}$-SdrG against Fg substrate in TBS with (**A**) 1 mM $Ca^{2+}$, (**B**) 10 mM $Ca^{2+}$ and against Fgβ substrate in TBS with (**C**) 1 mM $Ca^{2+}$, (**D**) 10 mM $Ca^{2+}$, all at 3000 rpm, and for N$_{mut}$-SdrG against Fg substrate in TBS with (**E**) 1 mM $Ca^{2+}$, (**F**) 10 mM $Ca^{2+}$ and against Fgβ substrate in TBS with (**G**) 1 mM $Ca^{2+}$, (**H**) 10 mM $Ca^{2+}$, all at 4000 rpm.

#### S6.2 Actual cell density after spinning

From each disk the absolute number of cells remaining is calculated from the coordinates of the cell locations on the surface, as is used to calculate the cumulative density plots. The average remaining cell density for each condition is given in Figure **S6.9**, this is then normalised to fit the data and extract the mechanism of adhesion, only the data from the N$_{mut}$-SdrG cannot be normalised and fitted as the final cell density is too low to extract any meaningful patterns in the distribution.

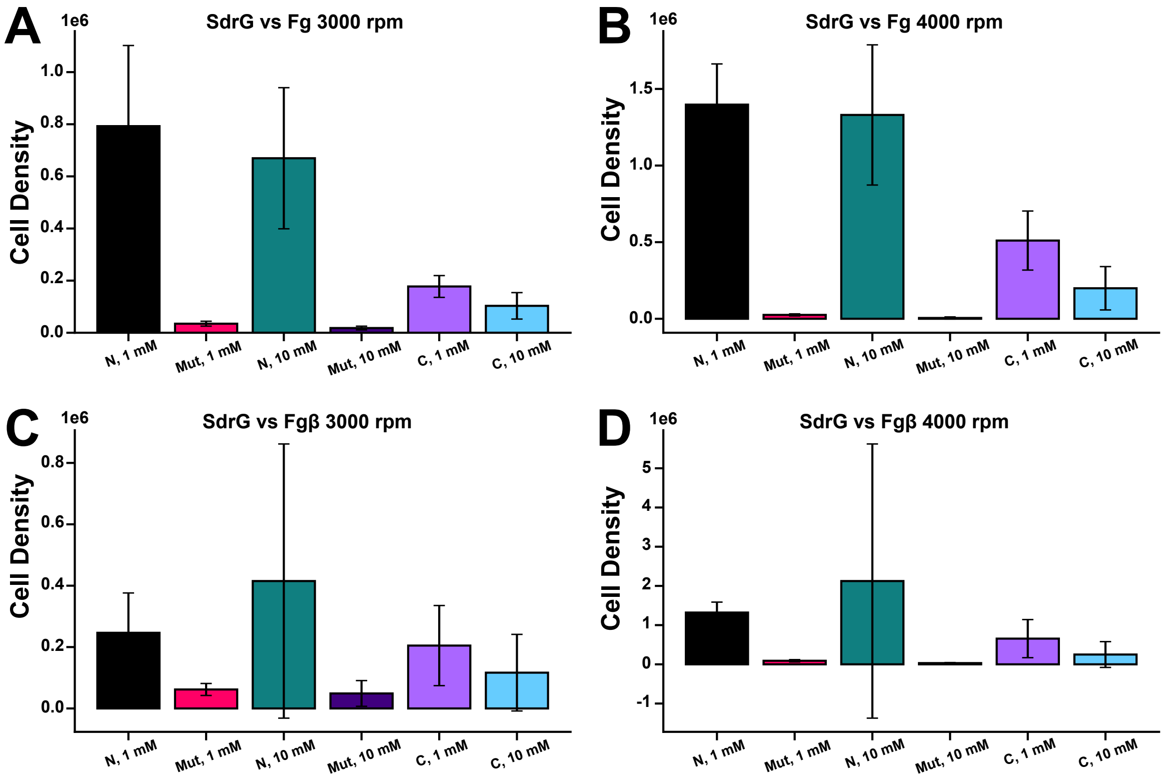


**Figure S6.3:** Actual cell adhesion after spinning at (**A**) 3000 rpm and (**B**) 4000 rpm against an Fg-coated substrate and at (**C**) 3000 rpm and **(D**) 4000 rpm against an Fgβ-coated substrate for N-SdrG (black = 1 mM, and green = 10 mM), N$_{mut}$-SdrG (pink = 1 mM, and dark purple = 10 mM), and C-SdrG (purple = 1 mM, and blue = 10 mM)

#### S6.3 Fitting of cell distribution data

Data from the 3000-rpm spinning is fit using either a combined sigmoid-gaussian (Figure **S6.4A**) or a pure sigmoid (Figure **S6.4B**), some datasets are not easily fit with the sigmoid due to their concave nature of the distribution, the $\tau_{50}$ for these populations is then estimated from the point on the x-axis at which the y-distribution has reached 0.5.

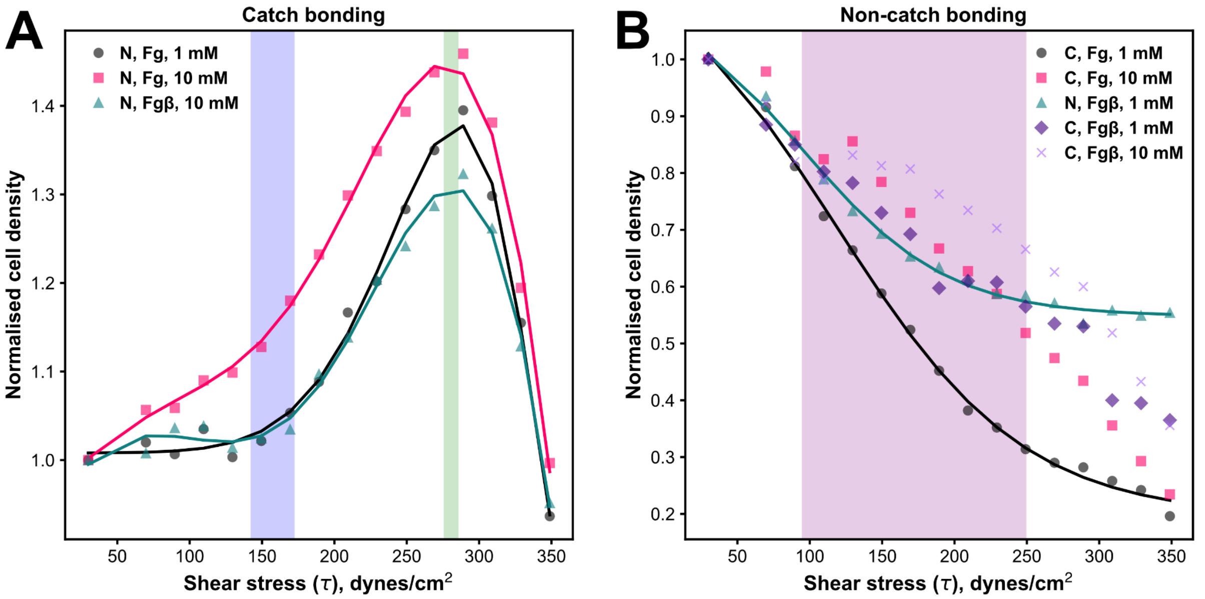


**Figure S6.4:** Plots of shear stress in dynes/cm$^2$ vs normalised cell density at 3000 rpm for (**A**) N-SdrG vs Fg at 1 mM or 10 mM $Ca^{2+}$ or Fgβ at 10 mM $Ca^{2+}$ and (**B**) N-SdrG vs Fgβ with 1 mM $Ca^{2+}$, and C-SdrG under all conditions.

Detailed fitting of each of the catch bond-displaying datasets with the sigmoid-gaussian fit, showing the activation of the catch bond and the catch bond peak are shown in Figure **S6.5**.

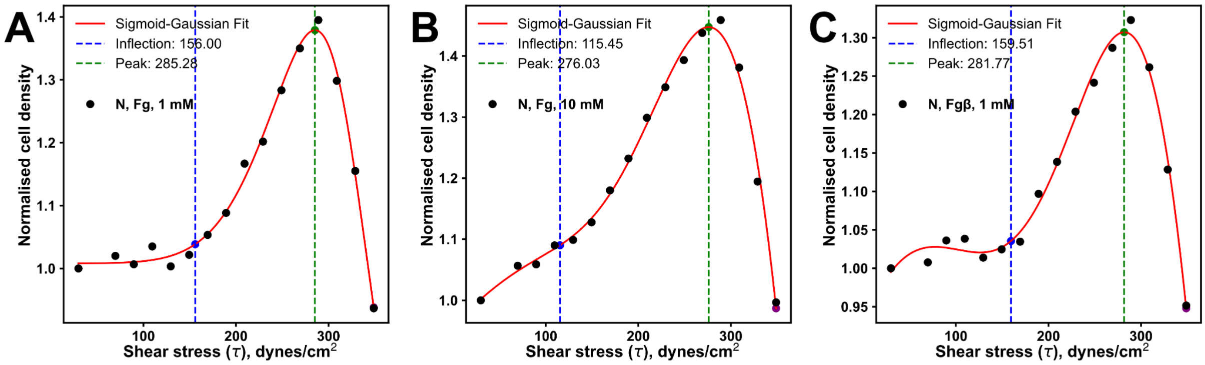


**Figure S6.5:** Plots of shear stress in dynes/cm$^2$ vs normalised cell density at 3000 rpm for (**A**) N-SdrG vs Fg with 1 mM $Ca^{2+}$ and (**B**) N-SdrG vs Fg with 10 mM $Ca^{2+}$, and (**C**) N-SdrG vs Fgβ with 10 mM $Ca^{2+}$ fitted with a combined sigmoid-gaussian fit, in which the activation of the catch bond (inflection) is shown with the blue dashed line and the catch bond peak is shown with a green dashed line.
